# CpG island turnover events predict evolutionary changes in enhancer activity

**DOI:** 10.1101/2023.05.09.540063

**Authors:** Acadia A. Kocher, Emily V. Dutrow, Severin Uebbing, Kristina M. Yim, María F. Rosales Larios, Marybeth Baumgartner, Timothy Nottoli, James P. Noonan

**Affiliations:** Department of Genetics, Yale School of Medicine, New Haven CT 06510, USA; Department of Comparative Medicine, Yale School of Medicine, New Haven, CT 06510, USA; Yale Genome Editing Center, Yale School of Medicine, New Haven, CT 06510, USA; Department of Ecology and Evolutionary Biology, Yale University, New Haven, CT 06520, USA; Department of Neuroscience, Yale School of Medicine, New Haven, CT 06510, USA; Wu Tsai Institute, Yale University, New Haven, CT 06510, USA; Cancer Genetics and Comparative Genomics Branch, National Human Genome Research Institute, National Institutes of Health, Bethesda, MD 20892, USA

## Abstract

Genetic changes that modify the function of transcriptional enhancers have been linked to the evolution of biological diversity across species. Multiple studies have focused on the role of nucleotide substitutions, transposition, and insertions and deletions in altering enhancer function. Here we show that turnover of CpG islands (CGIs), which contribute to enhancer activation, is broadly associated with changes in enhancer activity across mammals, including humans. We integrated maps of CGIs and enhancer activity-associated histone modifications obtained from multiple tissues in nine mammalian species and found that CGI content in enhancers was strongly associated with increased histone modification levels. CGIs showed widespread turnover across species and species-specific CGIs were strongly enriched for enhancers exhibiting species-specific activity across all tissues and species we examined. Genes associated with enhancers with species-specific CGIs showed concordant biases in their expression, supporting that CGI turnover contributes to gene regulatory innovation. Our results also implicate CGI turnover in the evolution of Human Gain Enhancers (HGEs), which show increased activity in human embryonic development and may have contributed to the evolution of uniquely human traits. Using a humanized mouse model, we show that a highly conserved HGE with a large CGI absent from the mouse ortholog shows increased activity at the human CGI in the humanized mouse diencephalon. Collectively, our results point to CGI turnover as a mechanism driving gene regulatory changes potentially underlying trait evolution in mammals.

## Introduction

Genetic variation in transcriptional enhancers has been associated with trait variation across species (1–8). Sequence changes in enhancers are hypothesized to modify the regulatory information enhancers encode by changing transcription factor binding site (TFBS) composition, thereby altering recruitment of transcription factors and co-activators (9–12). Such molecular changes may alter the spatiotemporal pattern and degree of enhancer activation and lead to corresponding changes in the expression of their target genes (13, 14). Numerous studies have characterized the contribution of nucleotide substitutions (3, 4, 6, 15–22), transposable elements (23–26), and insertions and deletions (27, 28) to evolutionary changes in enhancer function. However, despite these advances, understanding the specific mechanisms by which genetic variation modifies enhancer activity during evolution remains challenging.

One approach to identifying changes in enhancer function across species relies on comparing levels of histone modifications, such as histone H3 lysine 27 acetylation (H3K27ac), that are strongly associated with enhancer activity (29–32). These histone modifications can be mapped across the genome in order to identify regions, termed peaks, that show enriched levels of each modification. Several recent studies have compared levels of H3K27ac and other enhancer-associated histone modifications across multiple tissues in multiple mammalian species. These studies identified changes in histone modification levels at thousands of enhancers across species, suggesting abundant evolutionary turnover in enhancer activity across mammals (33–35).

This comparative approach has also been used to identify changes in enhancer activity relevant to human evolution. Two studies in developing human, rhesus macaque, and mouse cortex (36) and limb (31) identified Human Gain Enhancers (HGEs), defined as putative enhancers with higher levels of enhancer-associated histone modifications in human compared to the other two species. Subsequent Massively Parallel Reporter Assays (MPRAs) have identified sequence changes within HGEs that drive differential activity between human and chimpanzee orthologs (18). Chromatin interaction maps in mid-fetal human brain indicate that many HGEs target neurodevelopmental and neuronal genes (37), suggesting that altered HGE activity may have contributed to the evolution of uniquely human brain features.

Although previous studies have hypothesized that such changes in enhancer activity are due to genetic variation that altered transcription factor binding, CpG islands (CGIs) have also recently been found to contribute to enhancer activation. CGIs are genomic intervals with high GC-content and CpG dinucleotide frequency (38). They are frequently unmethylated, unlike the majority of CpG dinucleotides in the genome, and are associated with 70% of annotated promoters (39). However, CGIs are also located in intronic and intergenic regions. These “orphan CGIs (oCGIs)” are often located within enhancers that show higher levels of histone modifications, transcription factor binding, and three-dimensional interactions with other genomic regions than non-oCGI containing enhancers (40, 41).

Several mechanisms link CGIs with transcription-factor independent recruitment of chromatin modifiers. First, unmethylated CpG dinucleotides recruit proteins containing ZF-CxxC finger domains, which in turn recruit histone methyltransferase complexes that deposit H3K4me3. These include CFP1, a subunit of the SETD1 histone methyltransferase (42), and MLL2, a member of the SET1A/B complex (43). The presence of H3K4me3 promotes open, active chromatin by several mechanisms, including recruitment of histone acetylases, exclusion of factors that deposit repressive histone modifications, recruitment of chromatin remodelers, exclusion of DNA methylation, and direct recruitment of the transcriptional machinery (44). The recruitment of H3K4me3 and consequently all of these downstream effects are directly mediated by the CpG dinucleotides within CGIs and are independent of transcription factor binding events.

Given the abundance of evidence linking oCGIs to enhancer activity, we chose to examine oCGI turnover as a potential mechanism driving differences in enhancer activity across mammals. We identified oCGIs in nine mammalian genomes and integrated these maps with changes in the deposition of several enhancer-associated histone modifications. We first found that oCGIs are significantly enriched for H3K27ac, H3K4me3, H3K4me2, and H3K4me1 peaks in multiple tissues and species (31, 34, 36). We also found extensive turnover of oCGIs across species, and that species-specific oCGIs in putative enhancers were associated with species-specific increases in enhancer-associated histone modification levels. Orphan CGI turnover was associated with changes in enhancer activity even within ancient, highly conserved enhancers.

Our findings also support that oCGIs contribute to the increased activity of many HGEs. In light of these results, we selected one HGE to study *in vivo*, which has both an oCGI and prior evidence of increased regulatory activity in human compared to mouse. We generated a humanized mouse line for this locus and found that the oCGI-containing human enhancer exhibits increased H3K27ac and H3K4me3 in the developing mouse brain, although we did not observe associated changes in gene expression. However, using published data, we found that species-specific oCGIs with species-specific histone modifications were generally associated with increased expression of nearby genes. Finally, we found that turnover in transcription factor (TF) binding co-occurred at enhancers with oCGI turnover. We propose that oCGI turnover contributes to enhancer evolution both directly via altering recruitment of chromatin modifiers, and also indirectly by generating permissive chromatin that reveals TFBSs which may then be functionally integrated into nascent or existing enhancers. Our findings identify oCGI turnover as a novel class of sequence change, in addition to copy number changes and studies of evolutionary acceleration focused on nucleotide substitutions, that may be leveraged in comparative studies to identify gene regulatory innovations in mammalian genomes.

## Results

### oCGIs are significantly enriched for enhancer-associated histone modifications

We first examined a potential link between oCGIs and enhancer activity-associated histone modifications across a range of species and tissues. We identified genome-wide CGIs in nine mammalian genomes for which histone modification maps were previously generated (Fig. 1A). We used the canonical definition of CGIs: length ≥ 200bp, GC content ≥ 50%, and observed/expected CpG dinucleotides ≥ 0.6 (38). We restricted our analysis to oCGIs by excluding CGIs overlapping exons or located within regions 2kb upstream of a transcription start site, as annotated by RefSeq in every genome (45) (Fig. S1). For human and mouse, whose genomes have been more extensively annotated, we also removed CGIs overlapping ENCODE blacklist regions and regions 2kb upstream of transcription start sites identified by the FANTOM Consortium (46, 47). To further ensure that the identified oCGIs in each mammal were not part of unannotated exons or promoters, we only considered CGIs in each genome that had orthologous sequence in human that did not overlap human RefSeq, ENCODE blacklist, and FANTOM annotations (Fig. S1). After filtering, we found thousands of oCGIs in each mammalian genome, ranging from 11,067 in mouse to 77,199 in cat (Fig. 1A).

**Figure 1.**
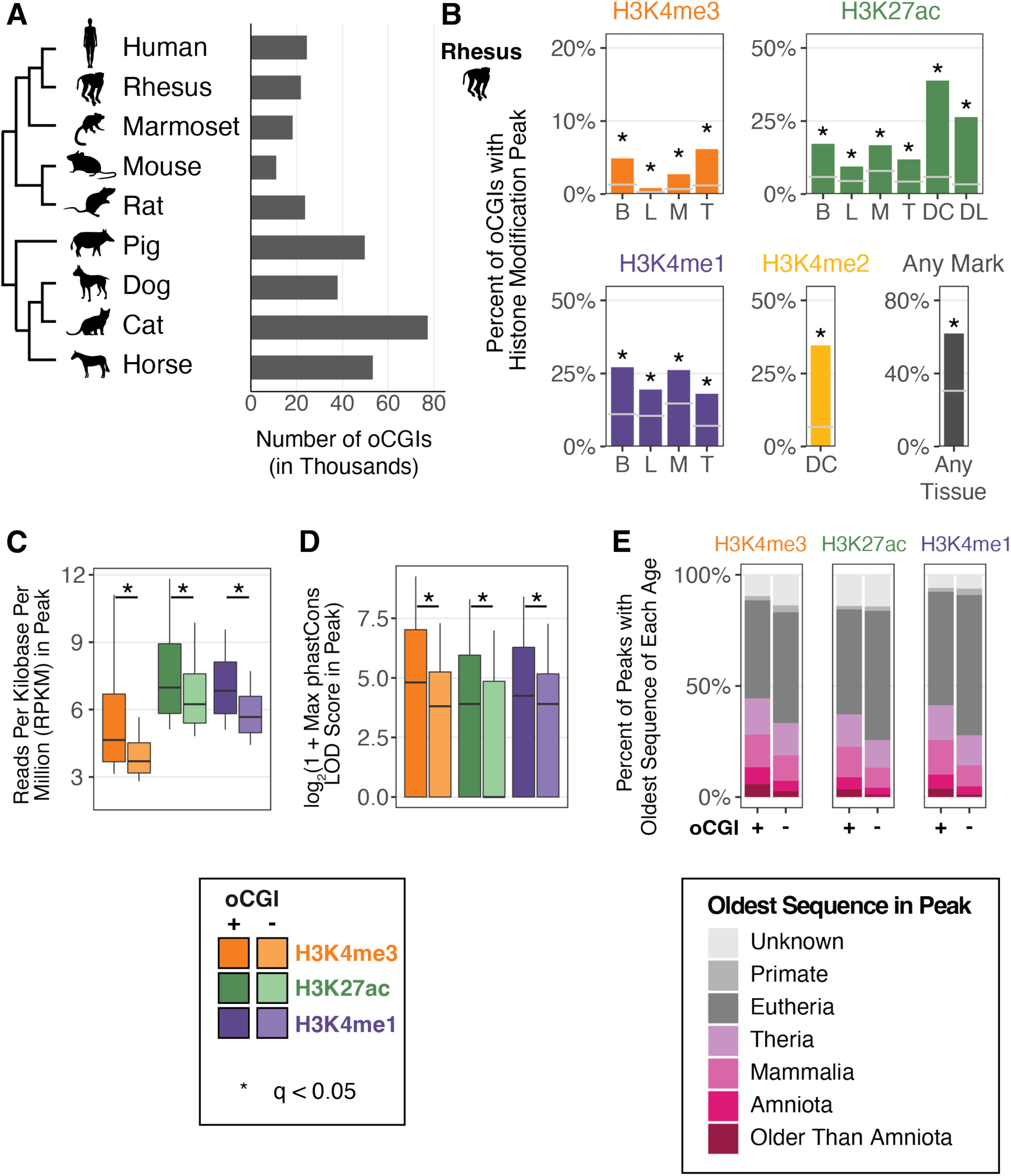
oCGIs are enriched for enhancer-associated histone modifications. (A) The number of oCGIs identified in nine mammalian genomes considered in this study. (B) Percent of oCGIs overlapping a histone modification peak for each indicated histone modification and tissue in rhesus macaque (31, 34, 36). B = adult brain, L = adult liver, M = adult muscle, T = adult testis, DC = developing cortex, DL = developing limb. Gray horizontal lines indicate the expected overlap and stars indicate significant enrichment (q < 0.05, BH-corrected, determined by permutation test; see Methods). (C) The level of each indicated histone modification in peaks with and without oCGIs, measured in Reads per Kilobase per Million (RPKM). Box plots show the interquartile range and median, and whiskers indicate the 90% confidence interval. Stars indicate a significant difference between peaks with and without an oCGI (q < 0.05, Wilcoxon rank-sum test, BH-corrected). (D) Maximum phastCons LOD (log-odds) scores in peaks with and without oCGIs. Box plots show the interquartile range and median, and whiskers indicate the 90% confidence interval. Stars indicate a significant difference between peaks with and without an oCGI (q < 0.05, Wilcoxon rank-sum test, BH-corrected). (E) Evolutionary origin of peaks with and without oCGIs. Bar plots show the percentage of peaks with and without oCGIs whose oldest sequence belongs to each age category. The results shown in panels (C) through (E) were generated using peaks from adult brain in rhesus macaque; see Fig. S5 and Fig. S7 for results from additional species and tissues.

The association between H3K27ac enrichment and enhancer activity has been previously established (29–32). We sought to evaluate whether other histone modifications included in this study were also associated with enhancer activity. We collected all peaks identified by the ENCODE consortium for the marks included in this study (H3K27ac, H3K4me3, H3K4me2, and H3K4me1) in five tissues (embryonic day 11.5 mouse forebrain, midbrain, hindbrain, heart, and limb) (30). We then intersected these peaks with sequences obtained from the VISTA Enhancer Browser that were tested for activity in a LacZ transgenic enhancer assay (termed VISTA elements; Fig. S2A) (48). We performed a separate analysis of VISTA elements that overlap an oCGI and those that do not overlap an oCGI in order to assess the predictive power of histone modifications in both cases. For all histone modifications in the analysis, we found that among VISTA elements lacking an oCGI, those that overlap a histone modification peak were significantly more likely to show transgenic enhancer activity than those not overlapping a peak (Fisher’s exact test, Benjamini Hochberg (BH)-corrected; Fig. S2B (*left*), Table S1). For VISTA elements that do contain an oCGI, those overlapping a histone modification peak were also more likely to show transgenic reporter activity for H3K27ac, H3K4me2, and H3K4me1, although the sample sizes in this analysis were smaller and did not always reach significance for H3K4me2 and H3K4me1 (Fisher’s exact test, BH-corrected; Fig. S2B (*right*), Table S1). These results support that the histone modification peaks included in this study are predictive of enhancer function, both for enhancers with and without oCGIs.

We next quantified the overlap of oCGIs in each species with published maps of each histone modification in multiple tissues (31, 34, 36). For all datasets, oCGIs were enriched for peaks from all histone modifications (permutation test, see Methods; Fig. 1B, Fig. S3). In all species, a large percentage (ranging from 36.7% in cat to 78.5% in mouse) of oCGIs overlap histone modification peaks in at least one dataset (“Any Mark” and “Any Tissue” in Fig. 1B, Fig. S3). This result supports that oCGIs are strongly associated with histone modifications indicative of enhancer activity. In light of this finding, we next sought to determine the proportion of putative enhancers that included an oCGI, and whose activity may thus be influenced by oCGI-mediated regulatory mechanisms. To address this question, we calculated the percentage of all histone modification peaks in each species and tissue that contained an oCGI. H3K4me3 peaks showed the highest overlap with oCGIs (in rhesus macaque, ranging from 21.9% of regions in testis to 49.5% in liver), consistent with the role of oCGIs in directly recruiting factors involved in H3K4me3 deposition (Fig. S4) (44). Although we found that H3K27ac, H3K4me1, and H3K4me2 peaks were less frequently associated with oCGIs, we still identified thousands of regions for each modification that contained oCGIs, revealing putative enhancers with possible oCGI-dependent functions (Fig. S4).

We next examined the association between oCGIs and the activity and evolutionary constraint of putative enhancers. We found that histone modification peaks containing an oCGI exhibited higher levels of each modification than peaks without an oCGI (Fig. 1C, Fig. S5), suggesting that oCGI-containing enhancers may show stronger activity than enhancers lacking an oCGI. Histone modification peaks with an oCGI were also longer than peaks without an oCGI, consistent with an oCGI-associated increased recruitment of histone modifiers across a broader region (Fig. S6). Peaks with an oCGI were more constrained, as measured using phastCons (49), compared to peaks lacking oCGIs. For example, in adult rhesus macaque brain, H3K4me3 peaks with an oCGI had higher maximum phastCons LOD (log-odds) scores compared to H3K4me3 peaks without an oCGI (log_2_-transformed median of 4.81 versus 3.81; log_2_-transformed upper quartile of 7.03 versus 5.25; significance determined by Wilcoxon rank-sum test, BH-corrected, see Methods) (Fig. 1D, Fig. S7). Peaks with an oCGI also had higher aggregate phastCons LOD scores and more bases included within constrained regions defined by phastCons (Fig. S8). We obtained further support for this finding using an age segmentation map of the human genome, in which the evolutionary origin of a human genomic region is inferred based on the most distantly related species in which an orthologous sequence can be identified (50). Peaks containing oCGIs were more likely to include ancient sequences compared to peaks lacking oCGIs (Fig. 1E). These trends generalized across all marks tested and across all species in our analysis (Fig. S9), supporting that oCGIs are components of ancient, constrained and active enhancers.

### Extensive turnover of oCGIs across mammalian species

To examine whether oCGI turnover is a widespread mechanism contributing to evolutionary changes in enhancer activity, we first asked how conserved oCGIs are by comparing species pairs within our dataset. For each species pair, we identified all oCGIs located within orthologous sequences. We then classified each oCGI as called in only one species (defined as species-specific within the scope of each pairwise comparison and labeled as “A-only” or “B-only” throughout the subsequent figures) or called in both species (labeled as “shared”) (Fig. 2A, Fig. S10). We found that, in every species pair we considered, the majority of oCGIs were present in only one species in the pair (Fig. 2B, see Fig. S11 and Table S2 for all species pairs). This was true both in closely-related and more distantly-related species. For example, in comparing the primate species rhesus macaque and marmoset, we found that 43.5% of oCGIs were rhesus-specific (A-only), while 37.4% were marmoset-specific (B-only), and 19.1% were shared (Fig. 2B). In comparing rat versus horse, which are more distantly related, we found that 36.6% of oCGIs were rat-specific (A-only), 56.0% were horse-specific (B-only), and 7.5% were shared (Fig. 2B).

**Figure 2.**
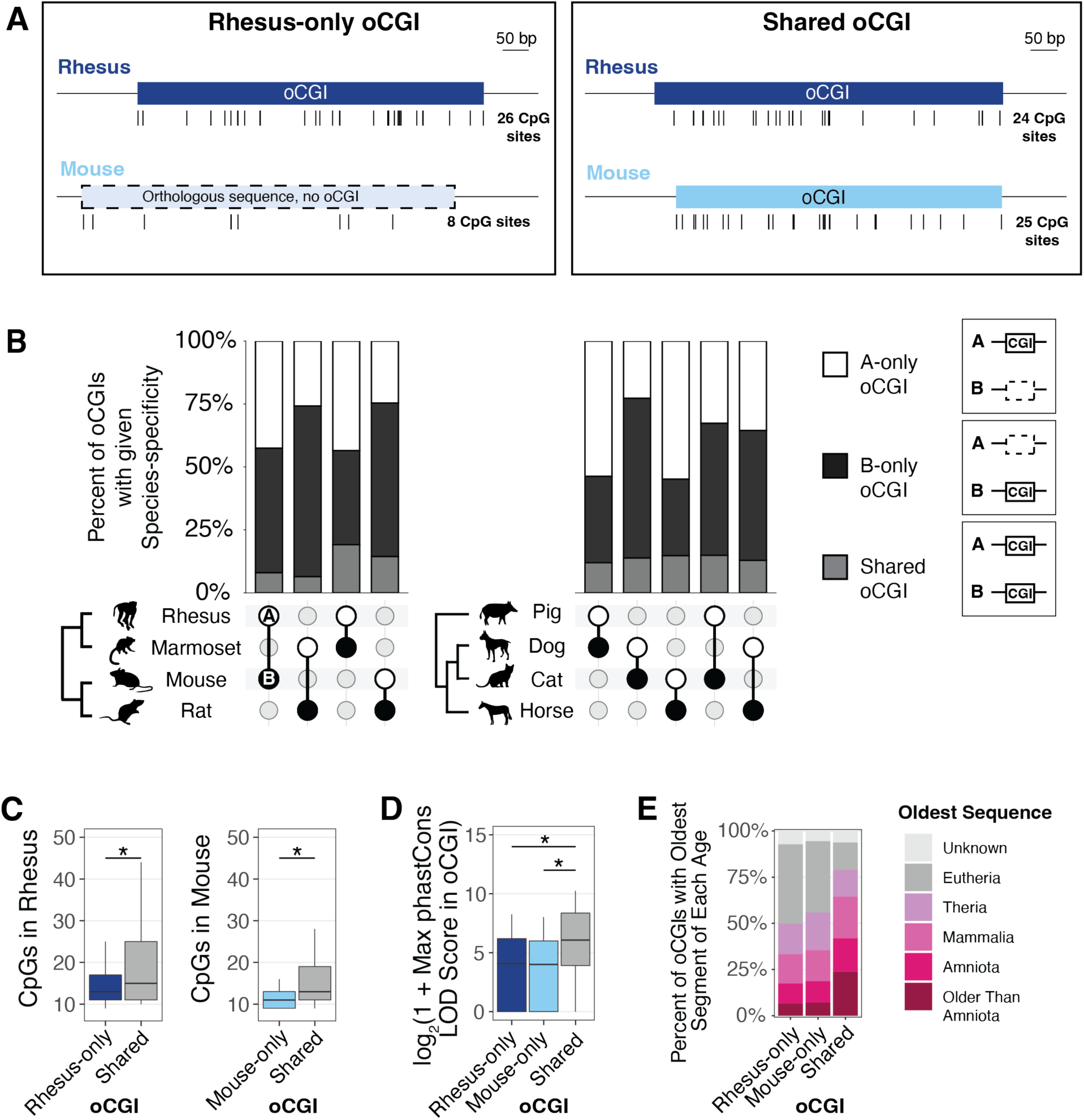
oCGIs show extensive turnover across species. (**A**) Schematic illustrating how we defined species-specific oCGIs in pairwise comparisons, using rhesus macaque and mouse as an example. *Left*: a rhesus-only oCGI (the sequence is present in both rhesus and mouse, but the oCGI is only present in rhesus). *Right*: a shared oCGI (both the sequence and oCGI are present in both rhesus and mouse). Ticks under each oCGI represent the locations of CpG dinucleotides. (B) Percent of oCGIs across the indicated species pairs (species A versus species B) that are “A-only,” “B-only,” or “shared” as described in the main text. The species pair is shown under each bar, with species A denoted by a white circle and species B denoted by a black circle. Percentages of oCGIs that are species A-only (white), species B-only (black), or shared (gray) are shown. (C) Number of CpG dinucleotides in rhesus-only (dark blue) or mouse-only (light blue) oCGIs compared to shared (gray) oCGIs. Box plots show the interquartile range and median, and whiskers indicate the 90% confidence interval. Stars indicate significant differences (q < 0.05 Wilcoxon rank-sum test, BH-corrected). (D) Maximum phastCons LOD scores in rhesus-only, mouse-only, and shared oCGIs. Box plots show the interquartile range and median, and whiskers indicate the 90% confidence interval. Stars indicate significant differences (q < 0.05, Wilcoxon rank-sum test, BH-corrected). (E) Evolutionary origins of rhesus-only, mouse-only, and shared oCGI sequences.

We next asked whether species-specific and shared oCGIs differ in their composition and their constraint. As a representative example, we show a comparison of rhesus macaque (species A) versus mouse (species B) in Figure 2C-E. We found that shared oCGIs had more CpG dinucleotides than rhesus-specific (A-only) or mouse-specific (B-only) oCGIs (Fig. 2C, see Fig. S12 for all species pairs), and were longer (Fig. S13). Shared oCGIs were more constrained, in that they were more likely to contain a higher scoring phastCons element (for rhesus versus mouse, log_2_-transformed median of 8.42 for shared oCGIs, compared to 4.95 for A-only oCGIs and 5.49 for B-only oCGIs; log_2_-transformed upper quartile of 10.14 for shared oCGIs compared to 7.11 for A-only oCGIs and 7.61 for B-only oCGIs) (Fig. 2D, see Fig. S14 for all species pairs). Shared oCGIs were more likely to have higher aggregate phastCons LOD scores and a greater percentage of bases covered by a phastCons element (Fig. S15). Shared oCGIs were also more likely to be located within more ancient sequences compared to species-specific oCGIs (Fig. 2E, see Fig. S16 for all species pairs).

Although shared oCGIs generally showed evidence of higher constraint, a substantial proportion of species-specific oCGIs did overlap with phastCons elements, evidence that they are located in sequences under constraint. In the rhesus macaque versus mouse comparison, 67.0% of rhesus-specific (A-only) oCGIs and 77.6% of mouse specific (B-only) oCGIs overlapped a phastCons element, compared to 99.2% of shared oCGIs. Additionally, many species-specific oCGIs were located in ancient enhancers: in the rhesus versus mouse comparison, 17.4% of rhesus-specific (A-only) oCGIs & 18.7% of mouse-specific (B-only) oCGIs were located in enhancers conserved among Amniotes or older clades, compared to 41.9% of shared oCGIs (Fig. 2E). These results support that oCGI turnover has occurred even within ancient, highly constrained enhancers.

### Species-specific oCGIs are significantly enriched for species-specific histone modification peaks

In light of the relationship we identified between oCGIs and enhancer activity-associated histone modifications, coupled with the extensive turnover of oCGIs we observed, we next examined whether species differences in oCGIs were associated with species differences in histone modification levels. Using pairwise species comparisons as described above, we sorted oCGIs based on their species-specificity and the species-specificity of co-localized histone modification peaks, performing a separate analysis for each species pair, tissue, and modification. Using a permutation test (Fig. 3A-B, Fig. S17, Methods), we found that species-specific oCGIs were significantly enriched for species-specific histone modification peaks; three representative pairwise comparisons are shown in Figure 3C-D. We also found that oCGIs specific to one species in the pair were depleted for histone modification peaks specific to the other species, and for peaks that were shared between both species. Shared oCGIs present in both species were also enriched for shared histone modification peaks, and were depleted for species-specific peaks. We observed the greatest enrichment of species-specific oCGIs for species-specific H3K4me3 peaks, but we observed significant enrichment for species-specific H3K27ac and H3K4me1 peaks as well (Fig. 3C-D). This trend was consistent across all species pairs and tissues, as demonstrated by the representative examples shown in Figures 3C and 3D and by all 28 species pairwise comparisons in adult brain, liver, muscle, and testis for H3K4me3 (Fig. S18-19), H3K27ac (Fig. S20-21), and H3K4me1 (Fig. S22-23) (Table S3). We also performed a peak-centric analysis (as opposed to the oCGI-centric analysis described above) in order to evaluate whether the patterns we observed were consistent between the two approaches (Methods). In this reciprocal analysis, we found that species-specific peaks were enriched for species-specific oCGIs (Fig. S24, Table S4), supporting the patterns observed in the oCGI-centric analysis.

**Figure 3.**
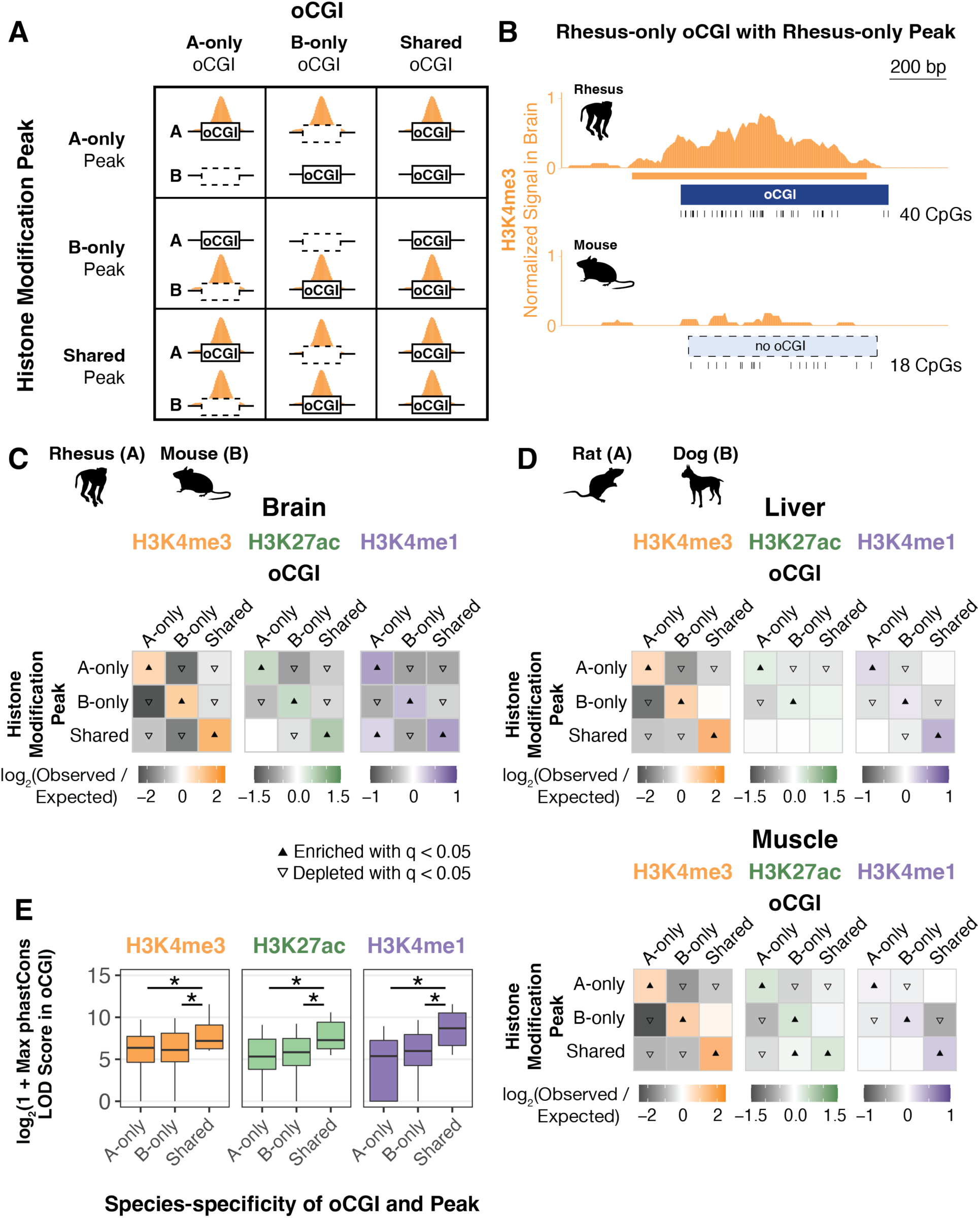
Species-specific oCGIs are significantly enriched for species-specific histone modification peaks. (A) Schematic illustrating how we defined species-specific and shared oCGIs and peaks. In each pairwise species comparison for each histone modification and tissue, we sorted oCGIs throughout the genome based on their species-specificity (designated as A-only, B-only, or shared as in Figure 2) and the species-specificity of their histone modification peaks (also designated as A-only, B-only, or shared, shown in orange in the schematic). (B) An example of a rhesus macaque-specific oCGI overlapping a rhesus-specific H3K4me3 peak in a pairwise comparison of rhesus macaque and mouse. Ticks show the location of CpG dinucleotides. The normalized H3K4me3 signal at this locus is shown in orange, measured as read counts per million in adjacent 10-bp bins. (C) Enrichment and depletion in each indicated comparison of species-specific and shared oCGIs (*top:* A-only, B-only, Shared) and species-specific and shared peaks (*left*: A-only, B-only, Shared), compared to a null expectation of no association between oCGI turnover and peak turnover. Each 3 x 3 grid shows the results for a specific test examining oCGIs and their overlap with three histone modifications in adult rhesus macaque brain: H3K4me3 (*left*), H3K27ac (*middle)*, and H3K4me1 (*right*). Each box in each grid is colored according to the level of enrichment over expectation (orange for H3K4me3, green for H3K27ac, or purple for H3K4me1) or depletion (gray for all marks) of genome-wide sites that meet the criteria for that box. The color bar below each plot illustrates the level of enrichment or depletion over expectation. The filled upward-pointing triangles denote significant enrichment and open downward-pointing triangles denote significant depletion (q < 0.05, permutation test, BH-corrected, see Fig. S17 and Methods). (D) Enrichment and depletion in an additional species comparison, rat versus dog, and in additional tissues (liver, *top,* and muscle, *bottom*), shown as described in (C). (E) Maximum LOD score in species-specific oCGIs in species-specific peaks and shared oCGIs in shared peaks, using data from adult rhesus macaque brain. Box plots show the interquartile range and median, and whiskers indicate the 90% confidence interval. Stars indicate significance (q < 0.05, Wilcoxon rank-sum test, BH-corrected).

We next examined whether species-specific oCGIs associated with species-specific changes in histone modification marking were under evolutionary constraint. Overall, shared oCGIs in shared peaks exhibited higher levels of constraint as measured by maximum phastCons LOD scores (Fig. 3E, see Fig. S25 for more species pairs). Shared oCGIs in shared peaks also more frequently overlapped sequences of greater evolutionary age (Fig. S26) than species-specific oCGIs within species-specific peaks. However, we also found that a substantial fraction of species-specific oCGIs located within species-specific peaks were under constraint (Fig. 3E). For example, in a comparison of species-specific rhesus macaque and mouse oCGIs and H3K27ac peaks in adult brain, 75.5% of rhesus-specific oCGIs in rhesus-specific peaks and 83.7% of mouse-specific oCGIs in mouse-specific peaks overlapped a constrained region annotated by phastCons, compared to 94.7% of shared oCGIs located in shared peaks. In this same comparison, we also found that 20.9% of rhesus-specific oCGIs in rhesus-specific peaks and 18.2% of mouse-specific oCGIs in mouse-specific peaks overlapped with a sequence conserved within Amniota or more ancient clades, compared to 29.5% of shared active oCGIs in shared peaks, suggesting that many species-specific oCGIs associated with species-specific peaks are components of ancient, constrained enhancers.

In order to assess whether the enrichment of species-specific oCGIs for species-specific peaks we observed in adult tissues was consistent in other contexts, we performed the same analysis on two datasets generated by independent studies of the human, rhesus macaque, and mouse developing cortex (36) and developing limb (31). Both studies compared H3K27ac levels across species, and the developing cortex dataset examined an additional mark associated with enhancer activity, H3K4me2. Both studies also generated histone modification maps across four developmental time points (Table S5), which allowed us to perform an independent comparison for each time point, species pair, tissue, and mark (Fig. S17, Methods). In each case we found that species-specific oCGIs were enriched for species-specific histone modification peaks and shared oCGIs were enriched for shared peaks, consistent with our previous findings in adult tissues (Fig. 4A, see Fig. S27 for all species pairs and time points).

**Figure 4.**
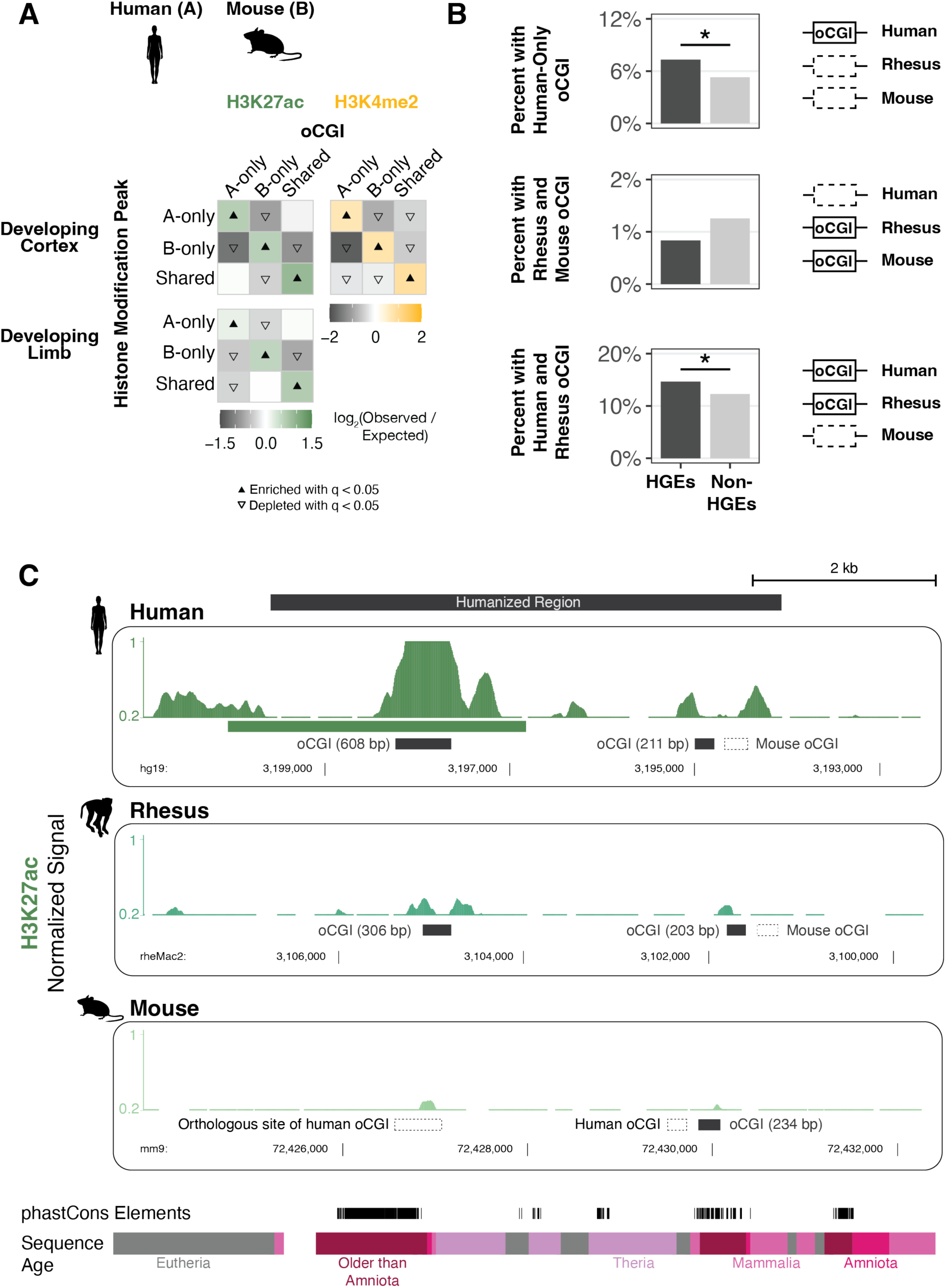
Association of species-specific oCGIs with species-specific histone modification peaks and HGEs in the developing human cortex and limb. (A) Enrichment and depletion in each indicated comparison of species-specific and shared oCGIs (*top:* A-only, B-only, Shared) and species-specific and shared peaks (*left*: A-only, B-only, Shared), compared to a null expectation of no association between oCGI turnover and peak turnover. As in Figure 3C-D, each 3 x 3 grid shows the results for a specific test examining oCGIs and their overlap with two histone modifications: H3K27ac (*left)*, and H3K4me2 (*right*). Each box in each grid is colored according to the level of enrichment over expectation (green for H3K27ac or yellow for H3K4me2) or depletion (gray for all marks) of genome-wide sites that meet the criteria for that box. The color bar below each plot illustrates the level of enrichment or depletion over expectation. The filled upward-pointing triangles denote significant enrichment and open downward-pointing triangles denote significant depletion (q < 0.05, permutation test, BH-corrected; see Fig. S17 and Methods). One representative comparison is shown for developing cortex (8.5 post-conception weeks (p.c.w.) in human versus embryonic day 14.5 in mouse) and developing limb (embryonic day 41 in human versus embryonic day 12.5 in mouse). (B) Enrichment of specific oCGI species patterns in HGEs compared to non-HGE enhancers in human cortex at 8.5 p.c.w. Bar plots show the percentage of HGEs (*left*) or non-HGE enhancers (*right*) that overlap an oCGI with the specified species pattern: oCGI in human only, oCGI in rhesus & mouse but absent in human, and oCGI in human & rhesus but absent in mouse. Significance was determined using a resampling test comparing HGEs to non-HGE human enhancers matched for overall histone modification levels (resampling test, BH-corrected; see Fig. S28 and Methods). (C) One representative HGE, hs754. H3K27ac levels are shown in developing cortex at human 8.5 p.c.w., rhesus embryonic day 55, and mouse embryonic day 14.5. H3K27ac signal tracks show the number of sequenced fragments per million overlapping each base pair. Black bars denote the locations of oCGIs in each species, and empty bars with dotted lines denote the locations where an orthologous sequence in another species contains an oCGI. Additional tracks show the locations of phastCons elements and sequences of the indicated evolutionary origin. For the purposes of visualization, features in rhesus and mouse have been aligned to the location of an orthologous base pair within the human peak due to overall differences in orthologous sequence lengths.

Both of these developmental studies also identified Human Gain Enhancers (HGEs) based on increased histone modification levels in human cortex or limb compared to rhesus macaque and mouse (31, 36). HGEs have been shown to exhibit human-specific changes in their activity and have been implicated in the regulation of neurodevelopmental genes (37), suggesting they may have contributed to human cortical evolution. Therefore, we examined whether oCGI turnover may have contributed to HGE evolution. We classified HGEs based on whether they had an oCGI in human, rhesus, and mouse, and compared them to non-HGE enhancers, defined as all other putative human enhancers that were mapped in each tissue and time point in each study, but that were not called as HGEs. We performed this comparison using a resampling test to randomly draw sets of non-HGE enhancers with activity level distributions matched to the activity level distribution of HGEs, as measured by total histone modification levels (Fig. S28, Methods). We found that cortex HGEs were significantly more likely to contain a human-only oCGI or an oCGI shared between human and rhesus, but absent in mouse (resampling test, BH-corrected; Fig. 4B, Fig. S29, Table S6). This finding suggests that changes in oCGI content may have contributed to the increased activity of HGEs in the human cortex. Consistent with this hypothesis, cortex HGEs were depleted for mouse-only oCGIs and oCGIs shared between rhesus and mouse, but absent in human. Limb HGEs were depleted across most oCGI categories and were enriched for having no oCGI at all, suggesting that oCGIs may be less relevant to enhancer activity in the developing limb than the developing cortex (Fig. S29).

### Changes in enhancer oCGI content are associated with changes in histone modification levels in a humanized mouse model

We next sought to study the effect of species differences in oCGI content on enhancer activity using an experimental approach. We selected one candidate HGE, named hs754 (36). This enhancer is highly constrained and includes sequences that are inferred to have originated in the stem lineage of jawed vertebrates (Fig. 4C, *bottom*). The human sequence is marked by an H3K27ac peak in the developing human cortex that was not called in the rhesus macaque or mouse cortex. The sequence underlying the human peak contains a 608-bp oCGI that has orthologous sequences in both rhesus and mouse. However, the rhesus orthologous sequence contains a smaller, 306-bp oCGI and the mouse orthologous sequence contains no oCGI (Fig. 4C). In order to determine whether this species difference in oCGI content is associated with changes in enhancer activity during development, we generated a mouse model in which a 5.5-kb human sequence containing hs754 replaced the mouse orthologous locus via CRISPR/Cas9-mediated homology-directed repair in C57BL/6 mouse embryos (Methods, Fig. S30, Table S7-8).

We first examined whether the humanized sequence containing an oCGI (Fig. 5A) showed increased levels of H3K27ac and H3K4me3 compared to the mouse sequence. We focused on developing diencephalon, based on a reported regulatory interaction between hs754 and the gene *Irx2* (51), which is expressed during diencephalon development (52). We included both an early (embryonic day 11.5, E11.5) and late (embryonic day 17.5, E17.5) time point.

**Figure 5.**
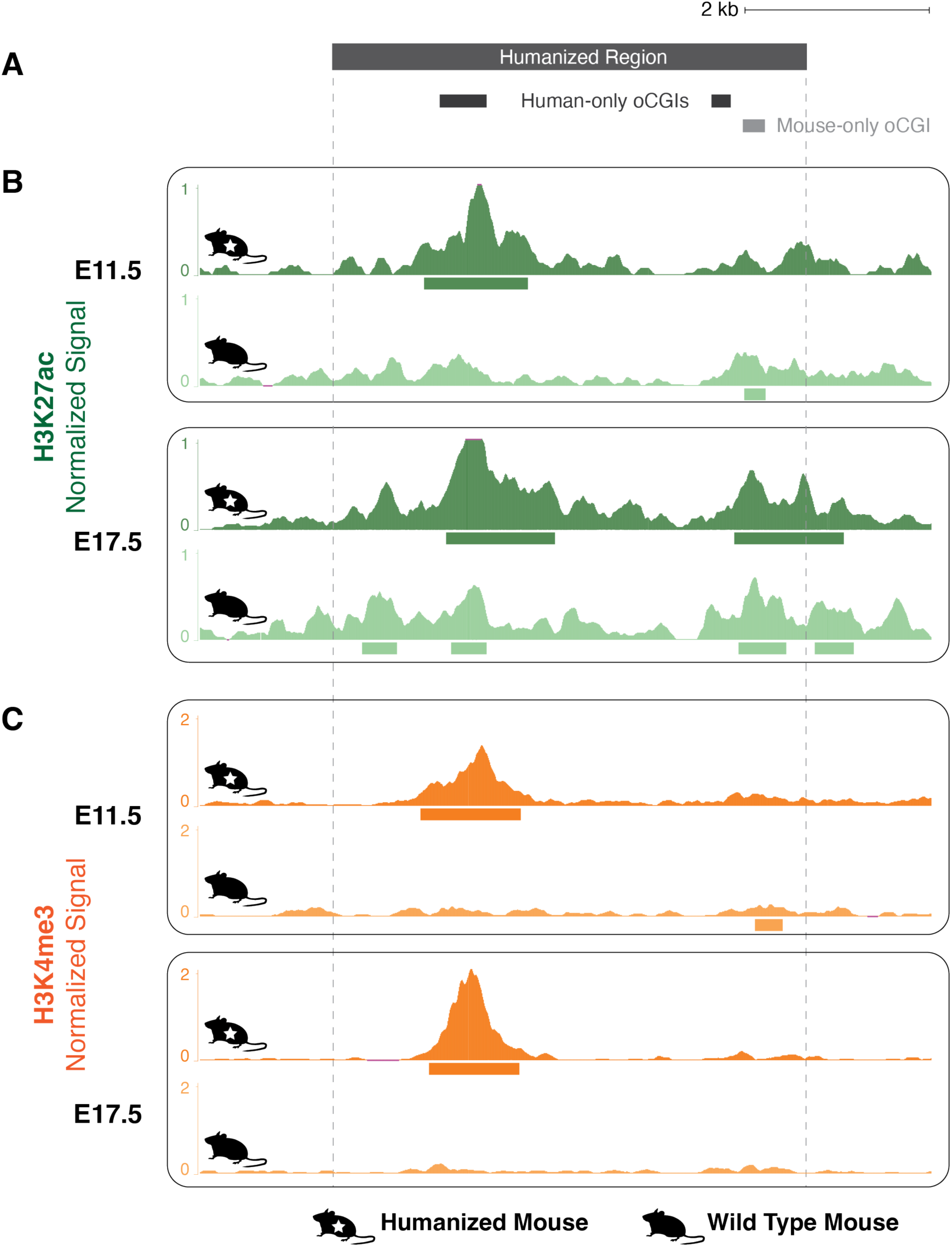
Gain of H3K27ac and H3K4me3 associated with a human oCGI in a humanized mouse model. (A) Locations of oCGIs within hs754 and its mouse ortholog. Dark gray boxes indicate the locations of two human oCGIs not present in the mouse sequence, and the light gray box indicates the location of a mouse oCGI not present in the human sequence. (B) H3K27ac levels in developing diencephalon at the humanized hs754 (*top*) or wild type (*bottom*) mouse locus at E11.5 and E17.5. Dark green (humanized) and light green (wild type) tracks show normalized H3K27ac levels as counts per million reads calculated in adjacent 10-bp bins. Peak calls are shown as boxes below the signal tracks. Nominal p-values were obtained by DESeq2 using a Wald test, then BH-corrected for multiple testing across all peaks genome-wide to generate q-values (see values in main text and in Fig. S31). (C) H3K4me3 levels in developing diencephalon at the humanized hs754 (*top*) or wild type (*bottom*) mouse locus at E11.5 and E17.5. Data are shown as in (B) but with H3K4me3 signal in dark orange (humanized) or light orange (wild type). The humanized hs754 locus is larger than the wild type locus, so for the purposes of visualization all humanized tracks have been shifted 190 bp to the left, bringing orthologous regions within the oCGI into alignment.

The humanized locus showed high levels of H3K27ac overlapping the human oCGI at both E11.5 and E17.5 (Fig. 5B, Fig. S31A-B). At both time points, the difference in H3K27ac level between the wild type and humanized orthologs was nominally statistically significant as measured using DESeq2, but did not reach significance after genome-wide multiple-testing correction (E11.5: p < 1.10 x 10^-5^, q = not significant; E17.5: p < 9.83 x 10^-4^, q = not significant; Wald test, BH-corrected). We also found a strong H3K4me3 peak overlapping the human oCGI at both time points, in contrast to the near absence of H3K4me3 in the wild type mouse (Figure 5C, Fig. S31C-D). These H3K4me3 peaks at both time points did reach significance after genome-wide correction (E11.5: p < 6.14 x 10^-20^, q < 4.23 x 10^-16^; E17.5: p < 1.66 x 10^-48^, q < 1.48 x 10^-44^; Wald test, BH-corrected). The full differential analysis results for all genome-wide peaks are shown in Figure S32 and Table S9. The overlap of the human oCGI with increased H3K27ac and H3K4me3 levels is consistent with the role of oCGIs in mediating changes in chromatin activity, and consistent with our genome-wide results across species pairs.

In order to determine whether these changes in enhancer activity were associated with downstream effects on gene expression, we carried out RNA-seq on E11.5 and E17.5 diencephalon (4 replicates per genotype at each time point). Using DESeq2, we found only two differentially expressed genes reaching genome-wide significance, both at embryonic day E11.5: the gene *Ppp3cc* which is on a different chromosome than hs754 (q < 1.98 x 10^-7^; Wald test, BH-corrected), and the gene *Serinc5* which is 20 Mb away from hs754 on the same chromosome (q < 4.37 x 10^-2^; Wald test, BH-corrected) (Fig. S33, Table S10). As reported *cis*-regulatory interactions over distances greater than several Mb are rare, as are *trans*-chromosome interactions (53), we consider it unlikely that either gene is a direct target of hs754.

Finally, we sought to understand whether the changes in histone modification levels at hs754 could stem not only from changes in oCGI content, but also from differences in the recruitment of transcription factors. The observed gains in histone modification levels both in human development and in the humanized mouse model coincide with a large oCGI that is present in the human ortholog, but not in mouse. However, the rhesus ortholog also contains an oCGI. Two mechanisms may explain why the rhesus sequence is not active during rhesus development. First, the size of the oCGI could play a role given that the rhesus oCGI is shorter than the human oCGI (306 bp compared to 608 bp). Second, there may be changes to TFBSs, either involving the oCGI or independent of it, in the human ortholog compared to rhesus. To assess this possibility, we performed ChIP-seq for the factor CTCF, which has several predicted motifs in hs754 and is involved in chromatin looping between enhancers and promoters (54–56). We identified two additional CTCF binding events in the hs754 humanized locus (Fig. S34). These changes in CTCF binding suggest that the gain of putative enhancer activity in the humanized mouse model may be due to a combination of oCGI-mediated mechanisms and changes in TF binding. We will return to the implications of this finding for enhancer evolution in the Discussion.

### Enhancers exhibiting oCGI and histone modification peak turnover are associated with gene expression changes across species

Although we did not observe changes in the expression of any potential target gene due to the oCGI-associated increase in enhancer activity in our hs754 humanized mouse model, such increases may nevertheless be generally associated with changes in enhancer activity. To evaluate this hypothesis, we used transcriptome datasets generated in (34) to determine whether species-specific oCGIs were correlated with increased expression of potential target genes across species pairs. We first identified all species-specific oCGIs in species-specific peaks, performing a separate analysis for each species pair, histone modification, and tissue. We associated these sites with potential target genes based on proximity (Methods; Fig. S35A) (57). We then focused on the subset of genes that are annotated as 1:1 orthologs between each species pair by ENSEMBL.

Taking as an example the comparison between rat and pig using adult brain H3K27ac peaks shown in Figure 6A, we sorted genes into a “rat-only set” which were associated with rat-only oCGIs in rat-only H3K27ac peaks, a “pig-only set” which are associated with pig-only oCGIs in pig-only H3K27ac peaks, and a “background set” not included in either category (Fig. S35A). For each gene in each set, we compared expression as the ratio of TPM (transcripts per million) in the two species. We then compared the log_2_-transformed TPM ratios in the rat-only set and the pig-only set to the background set (resampling test matching gene expression level, BH-corrected; see Methods and Fig. S35B-E). We found that genes associated with a rat-only active oCGI in a rat-only H3K27ac peak were more highly expressed in rat (median log_2_-transformed TPM ratio of 0.39), and genes associated with a pig-only active oCGI in a pig-only H3K27ac peak were more highly expressed in pig (median log_2_-transformed TPM ratio of -0.64). These results generalized across species pairs and tissues (Fig. 6, Fig. S36-S38), and reached statistical significance most often for species-specific oCGIs in species-specific H3K27ac peaks (Fig. S36, q < 0.05 in 62 of 120 tests), followed by those in species-specific H3K4me1 peaks (Figure S37, q < 0.05 in 52 of 120 tests) and those in species-specific H3K4me3 peaks (Figure S38, q < 0.05 in 24 of 120 tests). This finding suggests that oCGI turnover is not only associated with changes in enhancer activity, but also with downstream changes in gene expression.

**Figure 6.**
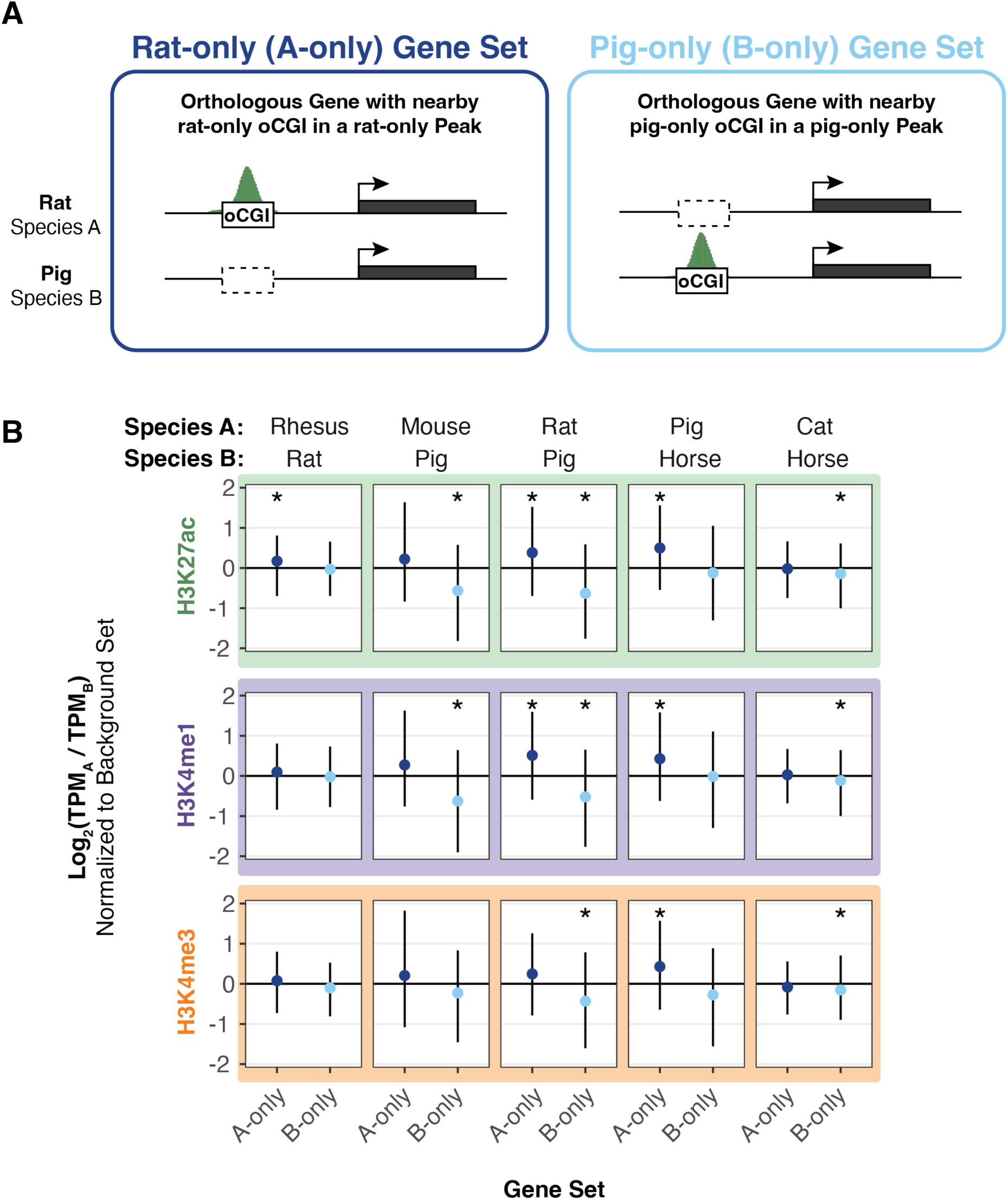
Species-specific oCGIs in species-specific peaks are associated with gene expression changes. (A) Schematic illustrating our method for assigning oCGIs and peaks to genes as described in the text and Figure S35, using a pairwise comparison of rat and pig as an example. *Left*: A gene associated with a rat-only oCGI in a rat-only H3K27ac peak, which means the gene is assigned to the “rat-only set” (A-only set) of genes. *Right*: A gene associated with a pig-only oCGI in a pig-only H3K27ac peak, which means the gene is assigned to the “pig-only set” (B-only set) of genes. (B) The log_2_-transformed TPM ratio for genes in the A-only set and the B-only set for each indicated species pair and histone modification using data from adult brain. Points indicate median values for the A-only set (dark blue) and the B-only set (light blue) and lines indicate the interquartile range. All values in the A-only set and B-only set were normalized to the median TPM ratio across resampling rounds from the background set. Stars indicate a significant difference between the observed median and the expected median (q < 0.05, resampling test to compare to the background set, BH-corrected; see Fig. S35 and Methods).

### Species-specific oCGIs are significantly enriched for species-specific transcription factor binding events

Orphan CGIs may contribute to enhancer activity by two mechanisms. The first is by direct recruitment of ZF-CxxC domain proteins such as CFP1 and MLL2, leading to increases in H3K4me3 and subsequent downstream effects (44). The second is via the TFBSs that they contain, which enable binding of TFs that then recruit coactivators. Many TFs have motifs with high GC content or that contain CpG dinucleotides, and could therefore be components of oCGIs (39).

To examine this second mechanism, we assessed whether species-specific oCGIs were enriched for species-specific transcription factor binding events. Using previously generated genome-wide binding data for several transcription factors in adult liver (58–60), we classified oCGIs based on their species-specificity and the species-specificity of co-localized TF peaks (Fig. 7A). For CTCF, we found that species-specific oCGIs were enriched for species-specific CTCF peaks (Fig. 7B, see Fig. S39A for all species pairs). Shared oCGIs were also enriched for shared CTCF peaks. This result is consistent with the sequence composition of the CTCF motif, which is GC-rich and contains a CpG dinucleotide, both features of oCGIs. We found similar enrichment patterns for an additional TF with a GC-rich motif (HNF4A, Fig. S39B) and a TF with a motif containing one CpG dinucleotide (HNF6, Fig. S39C), also consistent with oCGIs enabling binding of factors with GC-rich motifs and motifs containing CpGs.

**Figure 7.**
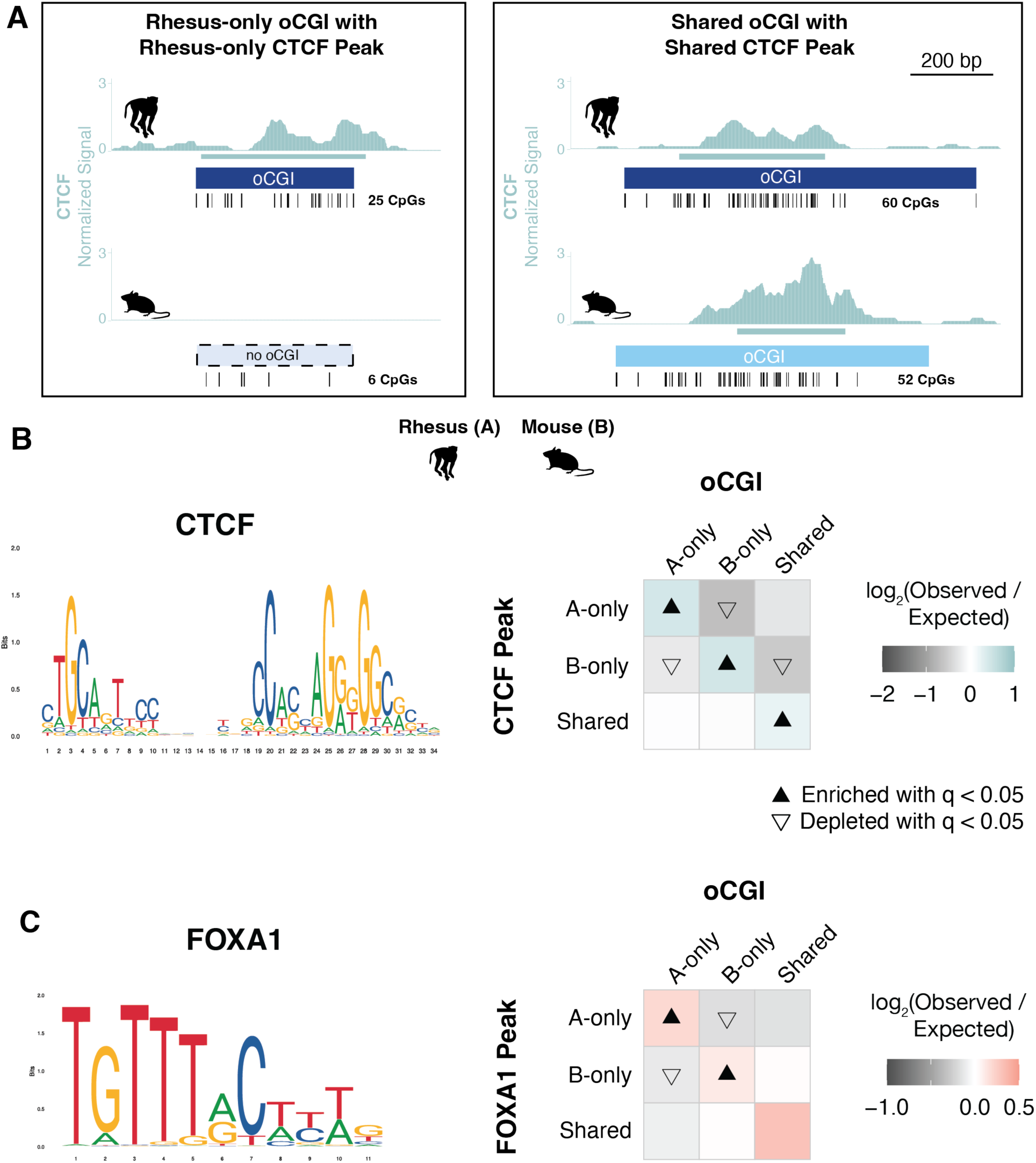
oCGI turnover is associated with changes in transcription factor binding. (A) Schematic illustrating how we compared species-specific oCGIs with species-specific transcription factor binding events in adult liver, using rhesus macaque and mouse as an example case. *Left*: a rhesus-only (species A-only) oCGI with a rhesus-only (species A-only) CTCF peak. *Right*: a shared oCGI with a shared CTCF peak. Ticks show the locations of CpG dinucleotides. (B) *Left*: the consensus motif for CTCF (MA1929.1 from the JASPAR database). *Right*: Enrichment and depletion in each indicated comparison of species-specific and shared oCGIs (*top:* A-only, B-only, Shared) and species-specific and shared CTCF peaks (*left*: A-only, B-only, Shared), compared to a null expectation of no association between oCGI turnover and peak turnover. Each 3 x 3 grid shows the results for a specific test examining oCGIs and their overlap with CTCF peaks. Each box in each grid is colored according to the level of enrichment over expectation (teal) or depletion (gray) of genome-wide sites that meet the criteria for that box. The color bar below each plot illustrates the level of enrichment or depletion over expectation. The filled upward-pointing triangles denote significant enrichment and open downward-pointing triangles denote significant depletion (q < 0.05, permutation test, BH-corrected; see Fig. S17 and Methods). (C) *Left*: the consensus motif for FOXA1 (MA0148.1 from the JASPAR database). *Right*: Enrichment and depletion in each indicated comparison of species-specific and shared oCGIs (*top:* A-only, B-only, Shared) and species-specific and shared FOXA1 peaks (*left*: A-only, B-only, Shared). Shown as in (B) but with boxes colored according to the level of enrichment over expectation (red) and depletion (gray) of genome-wide sites that meet the criteria for that box.

We next asked whether oCGIs were associated with the binding of other transcription factors with GC-poor motifs, namely FOXA1, a factor with an AT-rich motif containing no CpG sites. We found that species-specific oCGIs were enriched for species-specific FOXA1 binding (Fig. 7C, see Fig. S39D for all species pairs). Because the FOXA1 motif is AT-rich and contains no CpG sites, this enrichment is unlikely to be due to the sequence features of the oCGI and suggests that other oCGI-related mechanisms promote FOXA1 binding. Additionally, FOXA1 is a pioneer factor that is able to bind to its motif within closed chromatin (61, 62). Therefore, FOXA1 would not necessarily require open chromatin, such as the chromatin state generated by oCGIs, in order to bind. However, its binding is still enriched at oCGIs, suggesting that oCGIs do favor FOXA1 binding. Another pioneer factor with an AT-rich motif, CEBPA, was not associated with oCGIs, (Fig. S39E), suggesting that there is functional heterogeneity in oCGI recruitment of TFs. Nonetheless, our analysis of TF binding turnover suggests that evolutionary changes in the binding of multiple TFs is associated with oCGI turnover, and we will return to this finding in the Discussion.

## Discussion

Understanding the genetic and molecular mechanisms that drive evolutionary changes in gene regulation is essential for understanding how such changes contribute to the evolution of novel traits. Previous studies have focused on the role of nucleotide substitutions, transposable elements, and insertions and deletions to identify enhancers that may encode lineage-specific functions (3, 4, 20, 24, 25, 27, 28, 63). Here, we investigated the contribution of orphan CpG islands (oCGIs) to changes in transcriptional enhancer activity across species. Our findings support that oCGI turnover is associated with changes in the levels of several enhancer-associated histone modifications in mammals. We first found that oCGIs are enriched for histone modification peaks in all mammals we investigated, in line with previous findings in human (40, 41). Additionally, we identified extensive turnover of oCGIs across species. We then found that species-specific oCGIs were enriched for species-specific histone modification peaks in multiple developing and adult tissues, and this result was consistent in comparisons of both closely and distantly related species. We also found evidence that oCGI turnover is associated with changes in transcription factor binding events and changes in gene expression. Collectively, our results point to oCGI turnover as a major driver of gene regulatory innovation in mammalian evolution.

Our results also support that oCGIs contribute to the increased activity of Human Gain Enhancers (HGEs) in the developing cortex, which are hypothesized to alter gene expression during human brain development and contribute to uniquely human brain features. We experimentally modeled one such HGE, hs754, using humanized mice, and found that changes in oCGI content in the human ortholog were associated with *in vivo* changes in the levels of H3K27ac and H3K4me3, both associated with enhancer activity. These results are consistent with the role of oCGIs in active enhancers, and with the contribution of oCGI turnover to changes in enhancer activity we observed in our comparisons across mammals. However, we also identified correlated changes in CTCF binding at the humanized locus, which will need to be experimentally isolated from the effects of the human oCGI to determine the relative contributions of each to the increased enhancer activity of hs754.

Given the oCGI-associated changes in CTCF binding in our humanized mouse model, we sought to assess whether changes in TF binding might be coincident with changes in oCGIs. We found that species-specific oCGIs were enriched for species-specific TF binding for TFs with both GC- and AT-rich motif sequence compositions. As might be expected from the sequence characteristics of oCGIs, species-specific oCGIs were enriched for species-specific binding of TFs with GC-rich motifs and motifs containing CpG sites (CTCF, HNF4A, HNF6/ONECUT1). However, species-specific oCGIs were also enriched for FOXA1 binding events. FOXA1 has an AT-rich motif lacking any CpG site. This result suggests that CpG islands are associated with increased TF binding in a manner that is independent of their high GC- and CpG-content. We hypothesize that the gain of an CGI in an enhancer may not only add TFBSs due to their CpG-content, but that they also may promote the incorporation of TFBSs more generally. FOXA1 is a pioneer transcription factor with the ability to bind to its motif even within closed, nucleosomal DNA (61, 62). However, species-specific FOXA1 binding is still enriched at species-specific oCGIs, suggesting that oCGIs provide a favorable locus for FOXA1 binding to occur even given its function as a pioneer factor. We hypothesize that the presence of oCGIs that promote active, open chromatin is also likely to facilitate the binding of non-pioneer factors with both GC- and AT-rich motifs.

Given these results, we propose a model by which gain or loss of oCGIs influences the evolution of both new (Fig. 8A) and existing enhancers (Fig. 8B). Gain of oCGIs may occur via several mechanisms, including individual nucleotide substitution events or transposable element insertions. Transposable element insertion has been proposed as a way that new germ line differentially methylated regions, also CGIs, can arise (64). Another mechanism that may generate oCGIs is GC-biased gene conversion (gBGC), a process by which GC base pairs are preferentially fixed during meiotic recombination (65). Increased GC content would increase the number of CpG sites at a locus. To assess the potential impact of gBGC in our datasets, we identified gBGC tracts in each species in our analysis using the program phastBias (66). We found that a substantial proportion of oCGIs in each species overlaps a gBGC tract, especially in pig, dog, cat, and horse (Fig. S40). This analysis is consistent with the possibility that gBGC is a mechanism that generates oCGIs.

**Figure 8.**
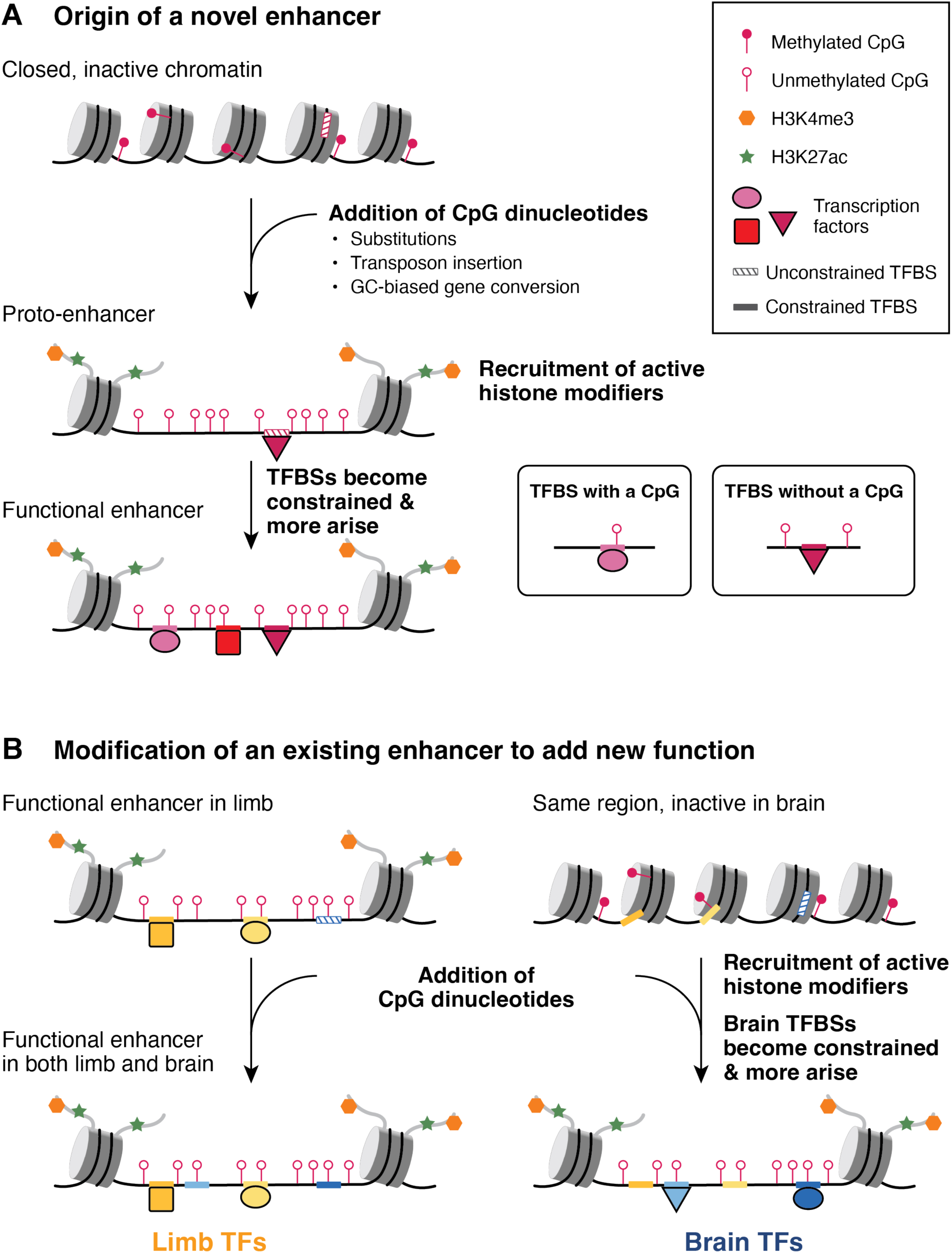
Model of enhancer evolution via oCGI turnover. (A) Evolution of a new enhancer from a locus in a closed chromatin state. This locus may include unconstrained, inaccessible TFBSs (striped boxes on DNA). DNA is depicted as a black line wrapped around cylindrical nucleosomes. After oCGI gain by several potential mechanisms (indicated in the figure), the site now acts as a proto-enhancer located within open, active chromatin recruited by the oCGI (44), which allows TFs to bind previously inaccessible TFBSs. A subset of histone tails (curved gray lines) with H3K4me3 (orange hexagons) and H3K27ac (green stars) modifications are shown. Filled lollipops indicate methylated CpGs, and unfilled lollipops indicate unmethylated CpGs. Over time, TFBSs become constrained (filled boxes on DNA) and additional TFBSs may arise and become fixed, resulting in the evolution of an enhancer with a constrained biological function. (B) Co-option of an existing enhancer in a novel biological context via oCGI gain. In an ancestral species, the enhancer is active in the developing limb and inactive in the developing brain, where the chromatin at the locus is closed. After oCGI gain, CpG-related mechanisms generate open chromatin in the developing brain, which allows existing unconstrained brain TFBSs to be bound. Over time, these and additional TFBSs may gain biological functions and be maintained by selection. The locus becomes a functional enhancer in the developing brain.

We propose that oCGIs may contribute to the evolution of novel enhancers via several mechanisms (Fig. 8A). Our model builds on a previously described model of *de novo* enhancer birth in the genome, which focused on the role of TFBSs in this process (50) and proposed that enhancers arise from “proto-enhancers,” which are regions of the genome containing TFBSs that are able to recruit TFs and subsequently histone modifiers, leading to deposition of enhancer-associated histone modifications. Although biochemically active, these proto-enhancers do not have biological functions, are not under evolutionary constraint and are rapidly gained or lost over time. However, in some cases these proto-enhancers may serve as nucleation points for novel enhancers to evolve via genetic changes that generate additional TFBSs, producing more complex enhancers with functional effects that may be favored and maintained by selection.

Gain of oCGIs may have similar effects, due to newly introduced CpG dinucleotides recruiting ZF-CxxC domain-containing proteins and associated histone H3K4 methylation machinery, thereby generating an open chromatin environment (44). Additionally, oCGIs can exclude DNA methylation (44, 67), which may alter the binding of methylation-sensitive TFs (68, 69). Orphan CGI-related chromatin changes may themselves contribute to increased enhancer function and recruitment of transcriptional machinery. They may also make existing, previously inaccessible and unconstrained TFBSs at the locus available to TFs, adding new regulatory functions now subject to selection. Additionally, the open chromatin environment generated by novel oCGIs may act as a nucleation point for the evolution of additional TFBSs that contribute new regulatory information. Over time, TFBSs may accumulate and generate an enhancer with biological functions under evolutionary constraint.

Our findings support that highly constrained enhancers have also gained and lost oCGIs (Fig. 3E). Therefore, we propose that oCGI gain may also contribute to the co-option of existing, constrained enhancers in novel biological contexts (Fig. 8B). An existing enhancer may be active in the developing limb, where it is bound by limb TFs that contribute to its function. In developing brain, however, the enhancer remains embedded within closed, inactive chromatin, and the limb TFBSs are not used. In our example, this constrained limb enhancer gains an oCGI and CpG-dependent mechanisms generate active, open chromatin in the brain where the region was previously closed. This new chromatin state may facilitate the evolution of binding sites for brain TFs, leading to a novel function for this enhancer in the brain.

Loss of oCGIs may also contribute to regulatory innovation by reducing enhancer activity. The tendency of CpG dinucleotides to mutate by deamination (38) makes oCGIs susceptible to decay. Decay could occur at weakly constrained enhancers, but also at highly constrained enhancers, since loss of regulatory activity may also lead to novel functions favored by selection. Additionally, we proposed above that oCGIs contribute to the initial opening of chromatin in a new context that allows existing TFBSs to become constrained and new TFBSs to emerge. It is possible that, once an enhancer has gained enough TFBSs to be strongly active, the CpG dinucleotides of the oCGI are no longer required for function (unless part of a TFBS) and begin to decay. In this way, oCGIs could act transiently to seed enhancers, then decay once no longer needed for function.

Turnover of oCGIs may also impact the evolution of poised enhancers, a class of enhancers whose function is dependent in part on oCGIs (70, 71). Poised enhancers are enriched near neurodevelopmental genes and are important for their activation during neural differentiation (72). Evolutionary turnover of oCGIs could therefore impact the function of poised enhancers and downstream neurodevelopmental processes. Recent work has identified poised enhancers in both mouse and human ESCs (73). We note that the enhancer we modeled in humanized mice, hs754, was identified as a poised enhancer only in human but not in mouse, in line with the presence of a large oCGI only in the human sequence. Further characterization of hs754 may reveal differences in poising, particularly at earlier developmental time points when neurodevelopmental genes are first activated, which is when poised enhancers are thought to function. Evolutionary changes in oCGI content that impact poised enhancer function during neurodevelopment may have implications for brain evolution, including in humans.

Our study identifies oCGI turnover as a novel mechanism affecting enhancer evolution in a variety of tissue contexts. Given this finding, oCGI content should be considered when assessing the mechanisms driving regulatory evolution across species and its impact on trait evolution. In the context of human and non-human primate evolution, several previous studies have focused on substitution events (15, 17, 21, 74) or deletions (27, 28), many of which are thought to alter TFBS content at enhancers. Our work highlights that an additional class of sequence change, oCGI turnover, is likely to reveal additional enhancers with lineage-specific functions, broadening the set of candidate regulatory innovations that may contribute to the evolution of novel traits.

## Methods

### Generating genome-wide CGI maps

We defined CGIs computationally (Fig. S1) using a program developed by Andy Law (75, 76). This program first scans the genome to identify all CpG dinucleotides. It then performs a windowing procedure to identify CGIs. Briefly, it selects the first CpG in the genome and adds downstream CpGs until the interval between two CpGs is at least 200 bp, when it performs a test for the following criteria: GC content of at least 50% and a ratio of observed over expected CpG dinucleotides of at least 0.6 (38). If the interval meets the criteria, it attempts to build a bigger CGI by continuing the process of adding a CpG and testing whether the criteria are met, then once the criteria are no longer met the interval is output as a CGI. Otherwise if the criteria are not met, the program drops the initial CpG and continues adding downstream CpGs until the length of 200 bp is met again, at which point it performs the test as above. This program generates more permissive CGI tracks than the default CGI tracks displayed by the UCSC Genome Browser, which imposes additional tests on CGI intervals to output only the largest, most CpG-dense CGIs (77).

We used the following genome versions in this study: rheMac10, calJac4, mm39, rn7, susScr11, canFam6, felCat9, and equCab3 for analyzing adult tissue data (34), and hg19, rheMac2, and mm9 for analyzing developing limb and cortex data (31, 36). Repeat-masked genomes were used for CGI identification.

### Filtering strategy to identify oCGIs

We restricted our analysis to oCGIs by excluding CGIs overlapping several categories of annotated gene-associated features including promoters (2kb upstream of transcription start sites, TSSs) and exons, as described in Figure S1 and Table S12, using NCBI and UCSC versions of RefSeq for each genome (45, 78). We also excluded CGIs falling in two additional annotated feature sets in the human and mouse genomes: promoters (2kb upstream of TSSs) annotated by the FANTOM Consortium based on CAGE (Cap Analysis of Gene Expression) data from several hundred human and mouse cell lines and tissues (46) and blacklist regions annotated by the ENCODE project in human and mouse (47). We further restricted our analysis to oCGIs with orthologous sequence in the human genome (hg38 for rheMac10, calJac4, mm39, rn7, susScr11, canFam6, felCat9, and equCab3, or hg19 for rheMac2 and mm9), which we identified using liftOver (78), that did not overlap features annotated in human (Fig. S1).

### Analysis of VISTA enhancer data

We downloaded bed files from the ENCODE Portal (https://www.encodeproject.org) containing annotated peaks for the four histone modifications used in this study (H3K27ac, H3K4me3, H3K4me2, H3K4me1) in five E11.5 mouse tissues (forebrain, midbrain, hindbrain, heart, and limb). We overlapped tested VISTA elements (https://enhancer.lbl.gov; see Results) with each peak set in each tissue using BEDTools intersect (79) and performed a separate analysis for VISTA elements that overlapped and did not overlap an oCGI in mouse (Fig. S2A). For each test, we then counted the number of VISTA elements falling into each of four categories based on their overlap with a ChIP-seq peak (overlap versus no overlap) and whether they showed transgenic reporter activity in the tissue predicted by the ChIP-seq data (reporter activity versus no reporter activity). We performed a Fisher’s exact test on the four categories in this contingency table to evaluate whether there was a significant association between the presence or absence of a ChIP-seq peak and reporter activity (Fig. S2B, Table S1). We corrected p-values across all tests using the Benjamini-Hochberg procedure (80). A result was considered significant if the q-value (BH-corrected p-value) was less than 0.05.

### Analysis of published ChIP-seq data

We downloaded FASTQ files for histone modification profiles obtained by ChIP-seq in adult tissues (34) from the ArrayExpress repository (accession number E-MTAB-7127). We downloaded FASTQ files for transcription factor profiles obtained by ChIP-seq in adult liver from the ArrayExpress repository (accession numbers E-MTAB-437 for CTCF, E-MTAB-1509 for FOXA1, HNF4A, HNF6/ONECUT1, and CEBPA). We excluded two species for which the available genomes were generated with less than 10X sequencing coverage: opossum (monDom5) and rabbit (oryCun2).

We mapped reads using Bowtie2 (81) (with settings specified by the flag “--very-sensitive) to unmasked genomes. We removed duplicate and multi-mapping reads using the Sambamba (82) commands “view” and “markdup.” We called peaks using MACS2 (83) with default settings, set to narrow peaks for H3K4me3, H3K27ac, and all transcription factors, and set to broad peaks using “--broad” for H3K4me1.

For histone modifications we discarded outlier replicates, defined as those with ≥ 20% fewer or ≥ 50% more peaks than the average of the other replicates for that species, tissue, and histone modification. For transcription factors we discarded replicates with low peak numbers, as was done previously with this data (60) for one replicate of rhesus CEBPA, rhesus HNF4A, rhesus FOXA1, and dog HNF6/ONECUT1. We also excluded one replicate with low peak numbers for the following species and TFs: dog FOXA1, rhesus HNF6/ONECUT1, rat HNF6/ONECUT1, and mouse FOXA1. We then identified reproducible peaks by taking the intersection of peaks in each replicate using BEDTools intersect.

For visualization, we used bamCoverage from deepTools (84) to generate bigWig files with the following settings: --normalizeUsing CPM --binSize 10 --extendReads 300 -- centerReads. These bigWig files summarize ChIP-seq signal data as extended read counts per million (CPM) in bins with a width of 10 bp. We generated bigBed files with peak intervals for each replicate (85).

For the histone modification ChIP-seq in developing cortex, we downloaded processed peak files and bigWig files from http://noonan.ycga.yale.edu/noonan_public/reilly2015/ (also available in the Gene Expression Omnibus under accession number GSE63649) (36). For species pair analysis, we matched time points between species as in Table S5. These bigWig files summarize ChIP-seq signal data as the number of sequenced fragments (extended to 300 bp) that overlap each base pair, normalized to 1 million aligned reads.

For the histone modification ChIP-seq in developing limb, we downloaded processed peak files and bigWig files from http://noonan.ycga.yale.edu/noonan_public/Limb_hub/ (also available from GEO under accession number GSE42413) (31). For species pair analysis, we matched time points between species as in Table S5. These bigWig files summarize ChIP-seq signal data as the number of sequenced fragments (extended to 300 bp) that overlap each base pair, normalized to 1 million aligned reads.

### Associating oCGIs and histone modification peaks

We measured the proportion of oCGIs in each species that overlap reproducible histone modification peaks using bedtools intersect (with flags “-wa -u”). Additionally, we performed a genomic reshuffling test to determine whether the overlap is greater than what would be expected if oCGIs and peaks were distributed independently throughout the genome (Fig. S3). For each species, we used bedtools shuffle (with flags “-chrom -noOverlapping”) to randomly redistribute oCGI intervals on each reference genome, excluding all regions with an annotated RefSeq feature (all promoters and all exons for protein-coding genes, lncRNAs, and other ncRNAs, plus introns for pseudogenes and features of an unknown type) and all regions falling in RepeatMasker regions because oCGIs were called on repeat-masked genomes. Since our oCGI sets are additionally filtered by lifting to the human genome and excluding human features, we also lifted each shuffled set to the human genome and filtered out oCGIs overlapping human features, then restricted the shuffled set in the original species based on its status in human. We then counted the percentage of this shuffled and filtered set overlapping a peak for each histone modification we examined in each tissue. We repeated the shuffling procedure 20,000 times and generated an expected value based on the mean percentage of shuffled oCGIs overlapping peaks for each tissue and histone modification across all 20,000 shuffling rounds. We calculated p-values by measuring the proportion of shuffling rounds in which the percentage of shuffled oCGIs overlapping peaks was more extreme than the observed percentage of oCGIs overlapping peaks. We corrected p-values across all species, tissues, and marks and a result was considered significant if the q-value (BH-corrected p-value) was less than 0.05.

We also measured the percentage of peaks that overlap oCGIs by first identifying intronic and intergenic peaks for each species, tissue, and histone modification, including lifting to human and filtering based on human feature annotations as for oCGIs. We then measured the percentage of peaks passing this filter that overlapped an oCGI in the original genome (Fig. S4).

We measured the histone modification levels in peaks with oCGIs and peaks without oCGIs (Fig. 1C, Fig. S5) by quantifying reads for adult tissue datasets using featureCounts from Subread (86) and normalizing reads per kilobase per million basepairs (RPKM). We quantified levels from bigWig files from developing tissue datasets using bigWigAverageOverBed, a measure analogous to RPKM.

### Analysis of sequence conservation and age

We downloaded 100-way Vertebrate phastCons elements in the human genome from the UCSC Genome Browser, generated using an alignment of 99 vertebrate species to human (hg38). Each phastCons element covers a specified interval in the human genome and is associated with a normalized LOD (log-odds) score reflecting the probability of the element being generated under the constrained phylo-HMM (phylogenetic hidden Markov model) compared to its probability under the non-constrained model (49). Higher LOD scores indicate a greater degree of inferred constraint. Throughout this study, we overlapped oCGIs and phastCons elements and reported several measures, including: maximum LOD score of overlapping phastCons elements, aggregate LOD scores of overlapping phastCons elements (the sum across all elements), and the proportion of bases covered by a phastCons element.

We also used an age segmentation map of the human genome to date oCGIs and histone modification peaks (50). This map dates intervals within the human genome based on the most distantly related species in the 46-Way MultiZ alignment (hg19) that has alignable sequence in that interval. The ages are as follows: Human, Ape, Primate, Eutheria, Theria, Mammalia, Amniota, Tetrapoda, Gnathostomata, and Vertebrata. Some intervals have an unknown age. We combined the most ancient three categories (Tetrapoda, Gnathostomata, and Vertebrata) into a category called “Older than Amniota.” The map was generated based on the human genome, so we infer sequence ages in the other species based on the age assigned to their orthologous human sequence. As a consequence, we converted the age of some oCGIs and peaks to “Unknown” if their assigned age was for a clade that did not include the species being analyzed; for example, if an oCGI in the pig genome was dated to age “Human,” we converted its age to “Unknown.”

### Identification of orthologous oCGIs across species pairs

For each species pair in the dataset (“species A” and “species B”), we identified oCGIs that are mappable between both species (using human as an intermediate), regardless of their oCGI status in both species. In other words, the underlying sequence was required to be present in both species, even if the oCGI was only called in one species. The procedure is described in Figure S10. We collected several additional pieces of information for each orthologous oCGI. We counted the number of CpG dinucleotides in each oCGI using faCount from the UCSC Genome Browser (Fig. 2C, Fig. S12). We also intersected the human coordinates of orthologous oCGIs between each species pair with phastCons elements (Fig. 2D, Fig. S14-15) and the age segmentation map of the human genome (Fig. 2E, Fig. S16), both of which are described above.

### Enrichment analysis of species-specific oCGIs and species-specific peaks

For each species pair, tissue, and histone modification or transcription factor, we intersected the set of species-specific and shared oCGIs with reproducible ChIP-seq peaks in both species in the pair, requiring 1 bp of overlap. We sorted species-specific (A-only, B-only) and shared oCGIs based on their overlap with species-specific (A-only, B-only) and shared peaks. It was also possible for an orthologous oCGI to overlap with no peak in either species; however, these sites were excluded from the analysis.

We determined enrichment and depletion in each category using a procedure described in Figure S17. We calculated p-values for enrichment and depletion using a permutation test (Fig. S17). A result was considered significant if the q-value (BH-corrected p-value) was less than 0.05.

We also performed a peak-centric version of this test (rather than oCGI-centric) (Fig. S24).

Instead of first identifying orthologous oCGIs between a species pair, we first identified orthologous histone modification or transcription factor peaks. We then counted their overlap with oCGIs and performed enrichment analysis and a permutation test as described above for the oCGI-centric analysis, although in this case the peak species-specificity labels were randomly permuted and the number falling into each oCGI category was counted. Statistical significance was determined as above, and a result was considered significant if the q-value (BH-corrected p-value) was less than 0.05.

### Association between oCGIs and HGEs

For the human developing cortex and limb datasets (31, 36), at each human time point (Table S5) we sorted all intronic and intergenic histone modification peaks based on their status as HGEs or as non-HGE human enhancers. We counted the proportion of HGEs with an oCGI species pattern in each of the following categories: human-only oCGI (absent in rhesus and mouse), rhesus-only oCGI (absent in human and mouse), mouse-only oCGI (absent in human and rhesus), oCGI shared between human and rhesus only (absent in mouse), oCGI shared between human and mouse (absent in rhesus), oCGI shared between rhesus and mouse (absent in human), oCGI in all three species, and oCGI in none of the three species. We compared these proportions (the observed values) to proportions in the non-HGE human enhancer set using a resampling test (Fig. 4B, Fig. S29, Table S6), described in Figure S28. A result was considered significant if the q-value (BH-corrected p-value) was less than 0.05.

### Mouse line generation and validation

The hs754 humanized mouse line was generated at the Yale Genome Editing Center by injecting the editing plasmid, purified Cas9 RNA, and sgRNA into C57BL/6J embryos, as previously described (87). The sequence of the CRISPR guide RNA was 5’ GAACCAAATATGGTGGGGAC. Coordinates of human and mouse sequences are in Table S7. F0 edited mice were backcrossed with C57BL/6J mice from Jackson Laboratory for several generations. All animal work was performed according to approved Yale IACUC protocols (#2020-07271, #2019-11167, #2022-11167).

Genotyping primers for the humanized and wild type locus are provided in Table S8, along with cloning primers for amplification of human and mouse genomic DNA for the generation of the editing construct and Sanger sequencing to verify the integrity of the edited locus. We amplified across both homology arms and the humanized region in both humanized and wild type mice, then cloned the products into pUC19 and used Sanger sequencing to verify each product. We also amplified 6.1 kb upstream and 3.8 kb downstream of the humanized locus, followed by Sanger sequencing, to establish that the extended locus was intact (Fig. S30B-C). We also established that the homozygous line carried two copies of the humanized haplotype using quantitative PCR (qPCR) (Fig. S30D). We performed copy number qPCR using genomic DNA from three humanized and three wild type individuals using Light Cycler 480 SYBR Green I Master Mix (Roche #04707516001). The biological replicates for each individual were run in triplicate and error bars show the standard deviation from these technical replicates. All Ct values were normalized to a control region on chromosome 5, and normalized again to the values for the first wild type individual. Copy number primers are provided in Table S8.

### Chromatin immunoprecipitation and ChIP-seq

We collected tissue from developing diencephalon at E11.5 and E17.5 from homozygous humanized crosses and homozygous wild type crosses. Each biological replicate came from 6 pooled embryos (at E11.5) or 3 pooled embryos (at E17.5) from a single litter, and each experiment involved two (at E11.5) or three (at E17.5) biological replicates. We crosslinked and sonicated tissue, then immunoprecipitated chromatin with antibodies for H3K27ac (Active Motif #91193, 15ug chromatin at E11.5 or 20ug chromatin at E17.5 and 5ug of antibody), H3K4me3 (Cell Signaling #9751S, 5ug chromatin and 5ug of antibody), or CTCF (Diagenode #C15410210-50, 15ug chromatin and 5ug of antibody) as previously described (88). Using the MODified peptide array from Active Motif (#13005 and #13006) we have found that the lot of H3K27ac antibody that we used is cross-reactive for histone H2B lysine 5 acetylation, in addition to H3K27 acetylation (Fig. S41). This modification has also been shown to predict active enhancers (89). Samples and their matched inputs were sequenced on an Illumina NovaSeq 6000 to generate paired 150 bp reads.

### RNA extraction and RNA-seq

We collected diencephalon tissue from 6 pooled embryos (at E11.5) or 3 pooled embryos (at E17.5) from a single litter per biological replicate, from 4 biological replicates per genotype. We purified RNA using the Qiagen miRNeasy Kit (#74106). The Yale Center for Genome Analysis prepared libraries using polyA-selection (Roche Kapa mRNA Hyper Prep Cat #KR1352) and sequenced them on an Illumina NovaSeq 6000 to generate paired 150 bp reads.

### Analysis of ChIP-seq data from the hs754 humanized mouse

We generated Bowtie2 indexes for the mouse genome (mm39) and for the humanized mouse genome, made by replacing the sequence of the mouse locus with the human sequence. Coordinates are shown in Table S7. We mapped ChIP-seq reads to the appropriate genome using Bowtie2 (v2.4.2) using the settings “--sensitive --no-unal.” We removed multi-mapping and duplicate reads using the Sambamba commands “view” and “markdup.” We called peaks using MACS2 with default settings set to narrow peaks for H3K27ac, H3K4me3, and CTCF.

For each time point and ChIP target, we generated a set of merged peaks across the genotypes in order to perform differential peak calling. The humanized allele of hs754 is 5531 bp, compared to 5141 bp for the replaced region in mouse, and this size discrepancy meant that an adjustment procedure was required in order to assign orthology and merge peaks between genotypes, which is described in Figure S42. We counted reads in all merged peaks using HTSeq (90), then called differential peaks in R using DESeq2 (91). Results for all peaks are shown in Table S9. A peak was considered significantly different if the q-value (BH-corrected p-value) was less than 0.05. We found several significant differential peaks on chromosome 19, which are all located within two known copy-number variants in the C57BL/6 mouse line (Fig. S43) (92). We hypothesize that in some comparisons, the samples used for wild type and humanized tissue contained different copy numbers of these loci. Because our testing for differential ChIP-seq peaks using DESeq2 uses raw read counts from the histone modification IP tracks, which are not normalized to input counts, the copy number variant led to calling these regions as differentially marked between the genotypes.

We generated bigWig files for visualization using used bamCoverage from deepTools with the following settings: --normalizeUsing CPM --binSize 10 --extendReads 300 --centerReads. These bigWig files show counts per million (CPM) in bins with a width of 10 bp. We also generated bigBed files with peak intervals for each replicate.

### Analysis of RNA-seq data from the hs754 humanized mouse

We generated a STAR index for mm39 using the unmasked genome and the basic gene annotation GTF from GENCODE release 31 (93). We mapped reads using STAR (v2.7.9a) and used “--quantMode GeneCounts” to directly output counts. We then used DESeq2 in R to perform differential expression analysis using an adjusted p-value cutoff of 0.05. Full results are shown in Table S10.

### Analysis of published RNA-seq data

In order to analyze differences in gene expression between species pairs, we analyzed genes annotated as 1:1 orthologs by Ensembl (94). This required mapping reads to Ensembl genomes for counting based on Ensembl GTF files. However, in order to integrate gene expression data with the orthologous oCGI sets which were called in UCSC genome versions, we restricted this analysis to species for which Ensembl release 106 offered GTF files built on the same NCBI genome assemblies as the UCSC genome versions (rhesus macaque, mouse, rat, pig, cat, and horse; see Table S11). We converted the chromosome names in these Ensembl GTF files to be compatible with UCSC chromosome names by adding “chr” in front of the numbers used by Ensembl.

We downloaded FASTQ files for RNA-seq from the ArrayExpress repository (E-MTAB-8122). We mapped reads using STAR to unmasked genomes (rheMac10, mm39, rn7, susScr11, felCat9, and equCab3) with the setting --sjdbOverhang 149 (95). We specified the Ensembl GTF files described above at this mapping step. STAR output the counts for each gene in each species, which we used to calculate TPM (transcripts per million) (96), using the R package GenomicRanges to calculate exon lengths for each gene from GTF files (97).

### Association between species-specific oCGIs in species-specific histone modification peaks and gene expression

Within each species, we defined the regulatory domain of each gene annotated in the Ensembl GTF using GREAT (57) in order to associate the gene with its putative enhancers. First, we assigned a basal regulatory domain to each protein-coding gene, which was 5 kb upstream and 1 kb downstream of the TSS. We then extended each basal regulatory domain to the next nearest basal regulatory domain upstream and downstream, up to a maximum distance of 1 Mb. We then sorted genes into four categories based on species-specific oCGIs in species-specific peaks that fell into their regulatory domains (performing a separate analysis for each species pair, tissue, and histone modification): the “A-only set” of genes associated with an A-only oCGI in an A-only peak, the “B-only set” of genes associated with a B-only oCGI in a B-only peak, the “mixed set” of genes associated with both types of oCGIs which was excluded from the analysis, and the “background set” containing all other genes (Fig. S35A). Note that this assignment procedure considers only species-specific oCGIs in species-specific peaks, ignoring all other oCGIs and peaks that may be associated with the gene.

We then compared the TPM ratio (TPM in Species A / TPM in Species B) for every gene in the A-only set and the B-only set to the background set using a resampling test (Fig. S35B-E). A result was considered significant if the q-value (BH-corrected p-value) was less than 0.05.

### Analysis of GC-biased gene conversion tracts

GC-biased gene conversion tracts were defined using phastBias (66), which finds regions where weak-to-strong substitutions are enriched compared to strong-to-weak substitutions. The program was run separately for each of the nine species in the analysis, taking that species as the foreground branch, using the setting “--output-tracts”. A whole-genome alignment for all nine species was extracted from a 120-mammal alignment (98) using maf_parse. The neutral rate was determined based on four-fold degenerate sites. These sites were extracted from the 120-mammal alignment using msa_view with settings “--4d” and “--features” from the human Gencode annotation (v41). Branch lengths were determined using phyloFit with the setting “--subst-mod REV” (99).

## Data and code availability

ChIP-seq and RNA-seq data for the humanized mouse model have been deposited under GEO accession GSE231307. All code for the study is available at GitHub: https://github.com/NoonanLab/Kocher_et_al_oCGI_turnover

## Supplemental Material

Supplemental Figures 1-43

Supplemental Tables 1-12

## Acknowledgements

This work was supported by a grant from the National Institute of General Medical Sciences (NIGMS) (R01 GM094780, to J.P.N.), a grant from the Eunice Kennedy Shriver National Institute of Child Health and Human Development (R01 HD102030 to J.P.N) and funds from the Yale School of Medicine (to J.P.N). A.A.K. was supported by an NSF Graduate Research Fellowship (DGE-1122492). E.V.D. was supported in part by NIGMS training grant T32 GM007499. K.M.Y. was supported by an NSF Graduate Research Fellowship (DGE-1752134). S.U. was supported in part by a Research Fellowship (352711928) from the Deutsche Forschungsgemeinschaft (DFG). M.B. was supported by an NIH F32 Postdoctoral Fellowship (NICHD) (F32 HD108935). This research program and related results were also made possible by the support of the NOMIS foundation (to J.P.N.). We thank members of the Noonan lab, B. Lesch and lab members, A. Louvi, G. Wagner and I-H. Park for their feedback on the manuscript, and L. Pennacchio for assistance with interrogating the VISTA Enhancer Browser database.

## Contributions

A.A.K., E.V.D., and J.P.N. conceived of and designed the study; A.A.K. performed all computational analyses related to oCGIs, with assistance from S.U. and K.M.Y.; E.V.D. designed the humanized mouse line and performed validation experiments; E.V.D. and T.N. generated the humanized mouse line; A.A.K. performed humanized mouse line validation experiments, ChIP-seq experiments, and RNA-seq experiments with assistance from M.F.R.L. and M.B.; A.A.K. analyzed data from the humanized mouse line; A.A.K. and J.P.N. wrote the manuscript with input from all authors.

