## Supplemental Figures for "CpG island turnover events predict evolutionary changes in enhancer activity"

#### Identification of oCGIs in Each Species

**Call CGIs in each masked genome**

Run tool built by Andy Law to call CGIs using the following criteria:

1. At least 200bp
2. GC content ≥ 50%
3. Observed / Expected CpG dinucleotides ≥ 0.6

Merge CGIs within 200bp

↓

**Filter to oCGIs**

Identify annotated features in NCBI and UCSC versions of RefSeq (using NCBI biotypes):

- Protein-coding genes
- lncRNAs
- Other ncRNAs
- Pseudogenes

Remove CGIs overlapping the following portions of annotated features:

- Promoters (2kb upstream of any transcription start site, TSS) for all feature types
- Exons for protein-coding genes, lncRNAs, and other ncRNAs
- Entire body including exons and introns of pseudogenes and features with unknown biotypes

Remove CGIs overlapping additional features for human and mouse:

- Promoters (2kb upstream of TSSs) annotated by the FANTOM consortium using CAGE (Cap Analysis of Gene Expression)
- ENCODE blacklist regions

↓

**Filter to oCGIs with orthology in human**

Lift to human (liftOver using minMatch = 0.3)

Remove CGIs overlapping human RefSeq, FANTOM, and ENCODE annotations

Lift back to original genome and keep only oCGIs that intersect with their original coordinates

**Figure S1. Pipeline for identifying oCGIs in nine mammalian species**
Flow chart of the steps involved in identifying oCGIs (*left*) and details of each step (*right*). See Table S12 for a description of how NCBI biotypes were used to classify features.

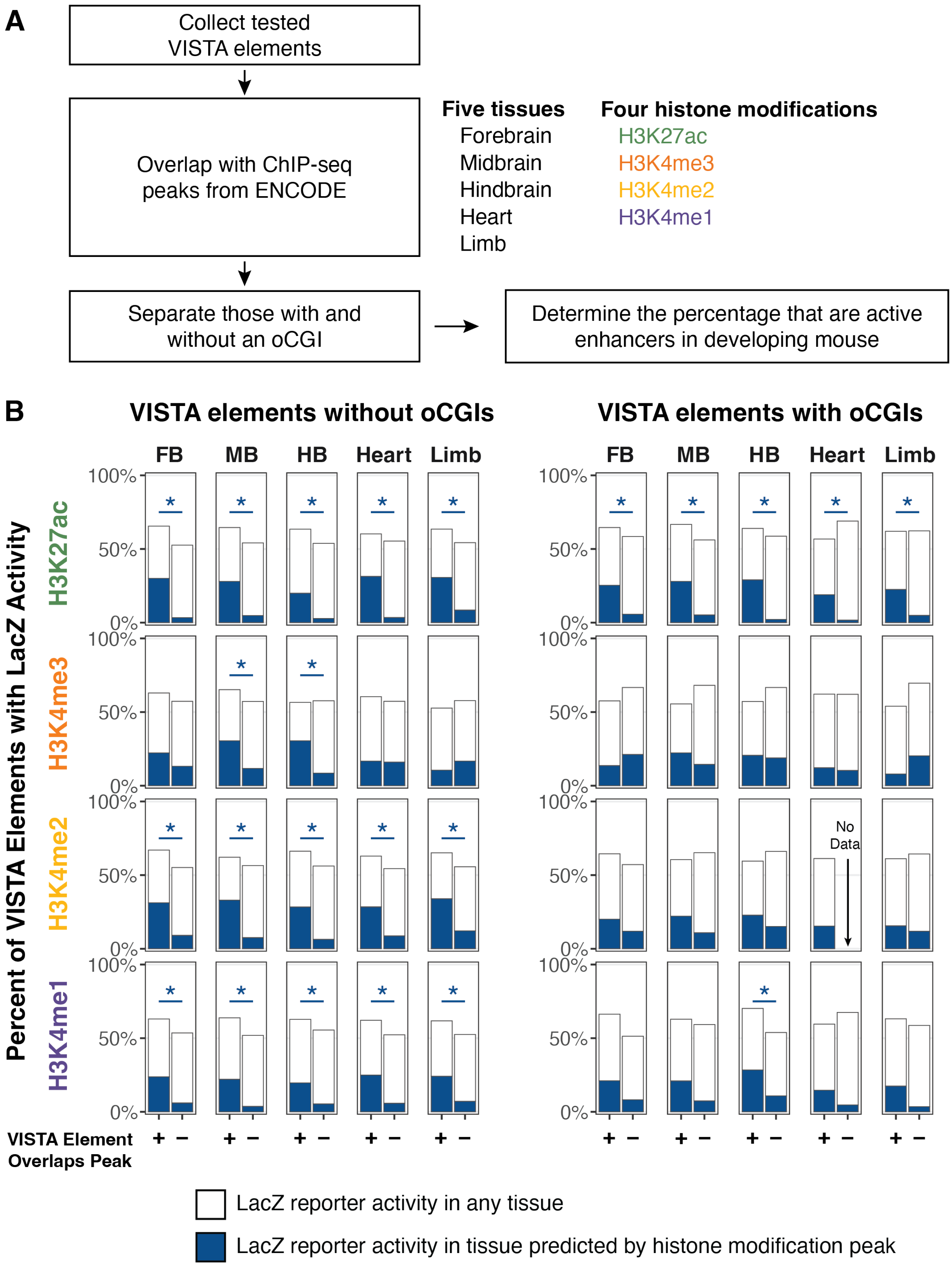

A
Collect tested VISTA elements
Overlap with ChIP-seq peaks from ENCODE
Separate those with and without an oCGI
Determine the percentage that are active enhancers in developing mouse
Five tissues
Forebrain
Midbrain
Hindbrain
Heart
Limb
Four histone modifications
H3K27ac
H3K4me3
H3K4me2
H3K4me1
B
VISTA elements without oCGIs
VISTA elements with oCGIs
FB
MB
HB
Heart
Limb
100%
50%
0%
*
No Data
Percent of VISTA Elements with LacZ Activity
VISTA Element Overlaps Peak
+
−
LacZ reporter activity in any tissue
LacZ reporter activity in tissue predicted by histone modification peak

**Figure S2. Histone modification peaks predict LacZ reporter assay activity**

(A) Workflow for testing whether histone modification peaks are predictive of transgenic reporter activity (VISTA Enhancer Browser) using ChIP-seq data from the ENCODE consortium across five tissues in embryonic day 11.5 (E11.5) mouse. (B) Percent of tested VISTA elements that are active enhancers in the tissue predicted by ChIP-seq peaks (dark blue) or active in another tissue (white). Each panel shows a separate bar for VISTA elements that overlap a histone modification peak in the specified tissue (*left bar*) or VISTA elements that do not overlap a peak in that tissue (*right bar*). A separate analysis was performed for VISTA elements that do not overlap an oCGI (*left panel*) and VISTA elements that do overlap an oCGI (*right panel*). Stars indicate a significant association between the presence or absence of a ChIP-seq peak and the number of VISTA elements that have reporter activity in the tissue predicted by the peak ($q < 0.05$, Fisher's exact test, BH-corrected).

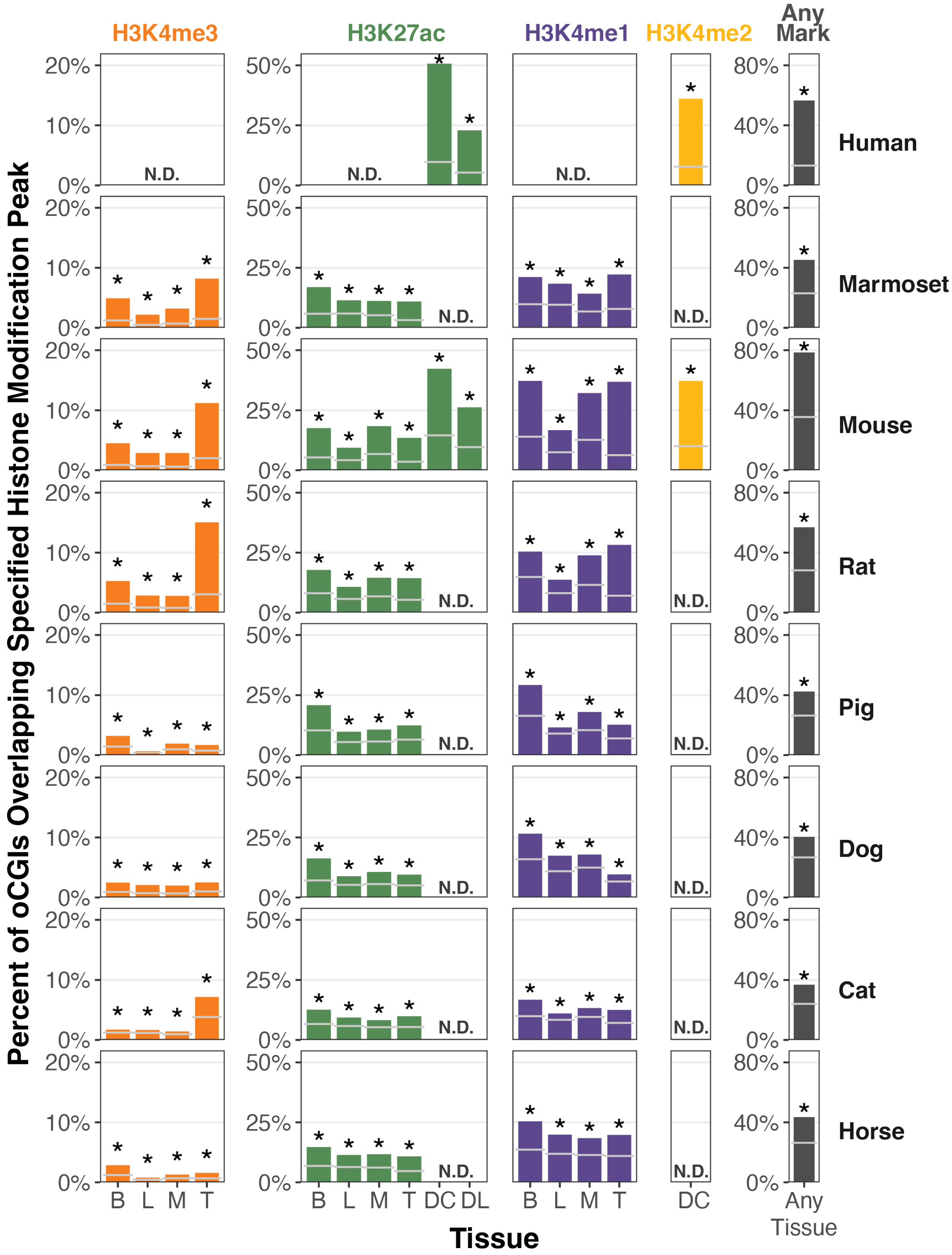

H3K4me3
H3K27ac
H3K4me1
H3K4me2
Any Mark
Human
Marmoset
Mouse
Rat
Pig
Dog
Cat
Horse
N.D.
20%
10%
0%
50%
25%
80%
40%
B L M T
B L M T DC DL
DC
Any Tissue
Tissue
Percent of oCGIs Overlapping Specified Histone Modification Peak

**Figure S3. oCGIs are enriched for active histone modifications in all species**
Percent of oCGIs overlapping a peak for each histone modification and in each tissue in all species except rhesus macaque (which is shown in Figure 1B). B = adult brain, L = adult liver, M = adult muscle, T = adult testis, DC = developing cortex, DL = developing limb. Gray horizontal lines indicate the expected overlap from a permutation test where oCGIs were reshuffled on the intronic and intergenic genome (see Methods). Stars indicate significance ($q < 0.05$ from this permutation test, BH-corrected). N.D. = no histone modification data available.

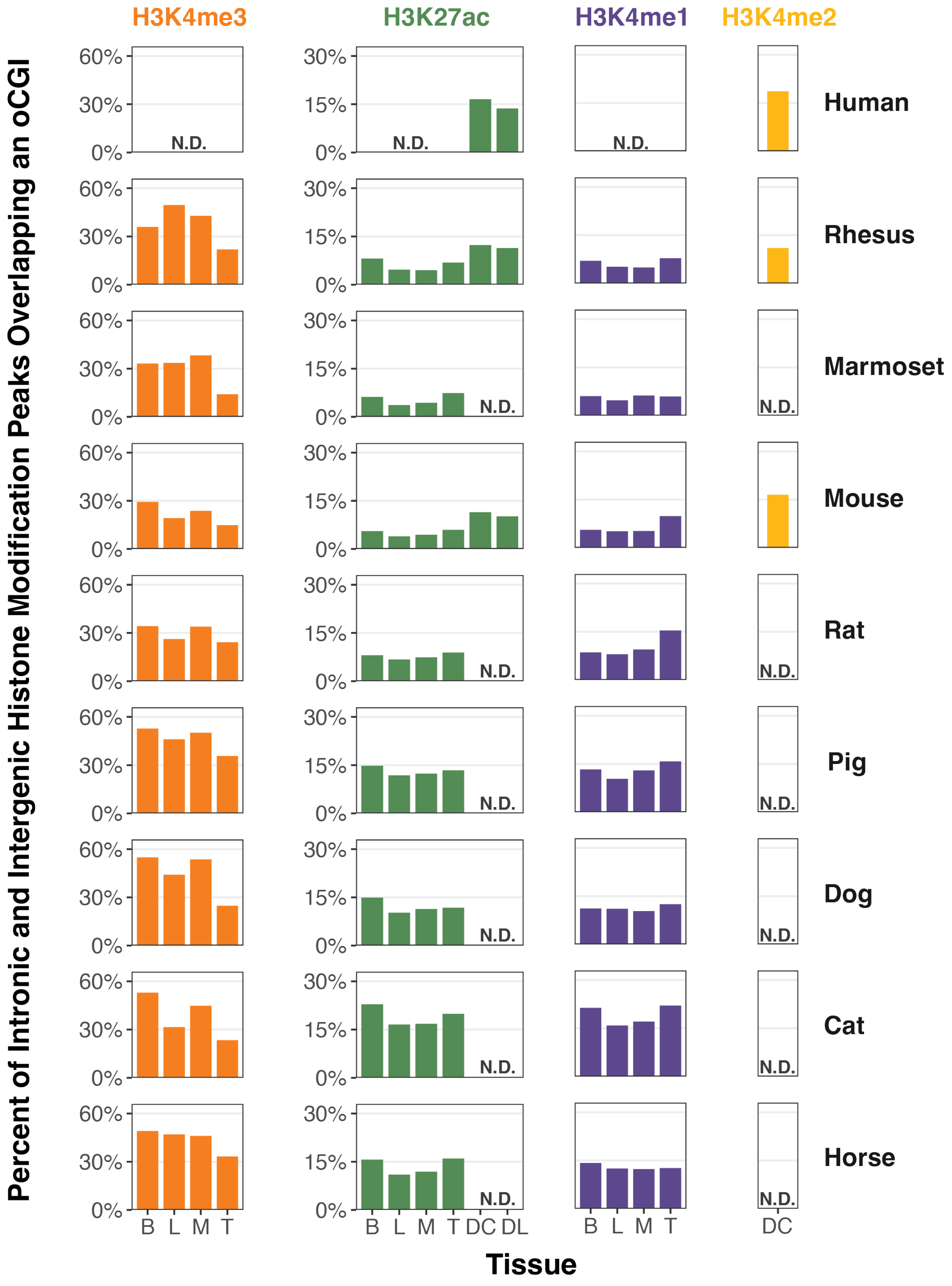

H3K4me3
H3K27ac
H3K4me1
H3K4me2
Percent of Intronic and Intergenic Histone Modification Peaks Overlapping an oCGI
60%
30%
0%
15%
N.D.
Human
Rhesus
Marmoset
Mouse
Rat
Pig
Dog
Cat
Horse
B
L
M
T
DC
DL
Tissue

**Figure S4. Percent of histone modification peaks that overlap oCGIs**
Percent of peaks for each histone modification in each tissue and species that overlap an oCGI. B = adult brain, L = adult liver, M = adult muscle, T = adult testis, DC = developing cortex, DL = developing limb. N.D. = no histone modification data available.

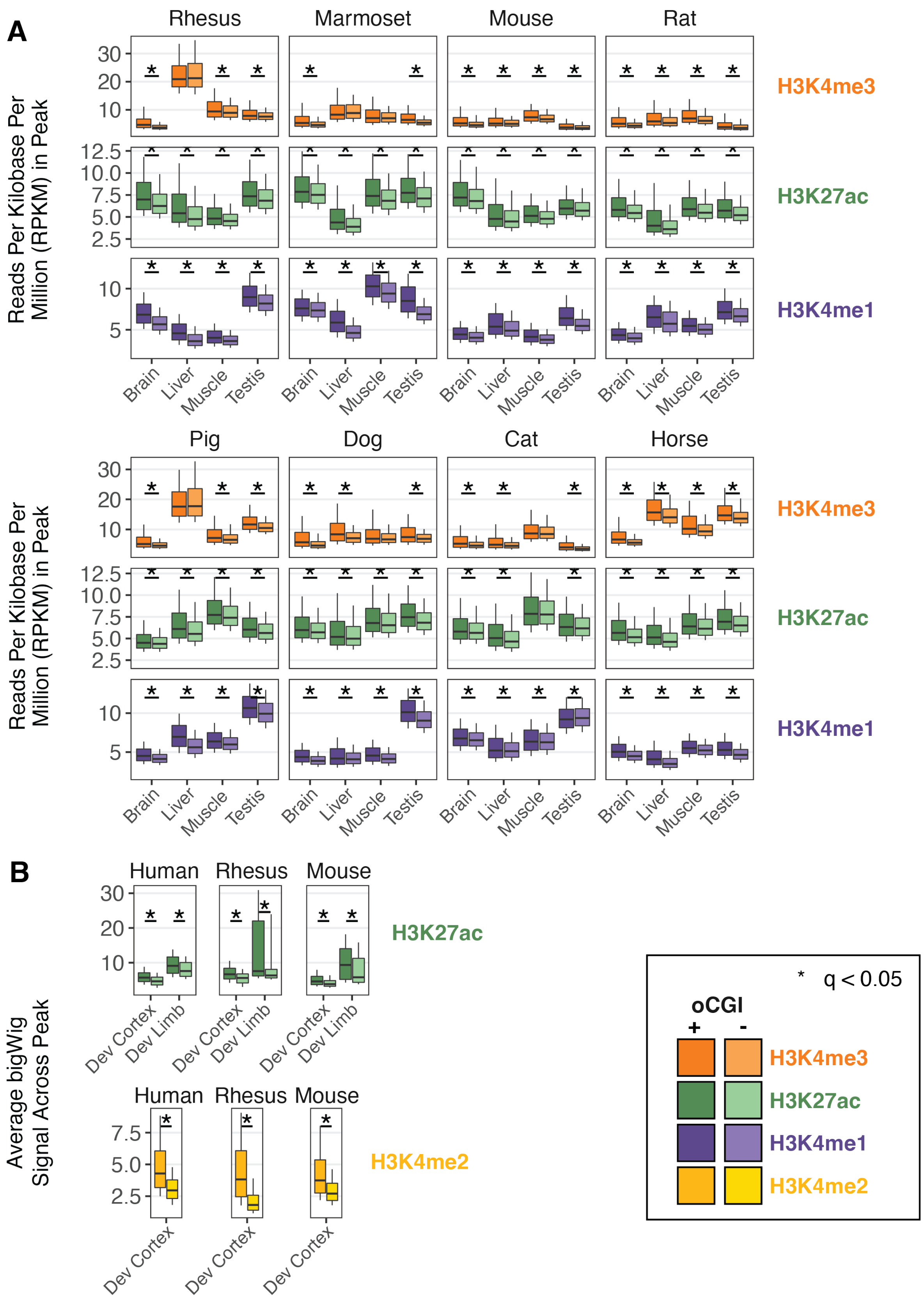

A
Rhesus
Marmoset
Mouse
Rat
Reads Per Kilobase Per Million (RPKM) in Peak
H3K4me3
H3K27ac
H3K4me1
Brain
Liver
Muscle
Testis
Pig
Dog
Cat
Horse
B
Human
Rhesus
Mouse
Average bigWig Signal Across Peak
Dev Cortex
Dev Limb
H3K4me2
* q < 0.05
oCGI
+
-

**Figure S5. Histone modification peaks with oCGIs have higher histone modification levels**
(A) RPKM (reads per kilobase per million mapped reads) for peaks with and without an oCGI from adult tissues. Box plots show the interquartile range and median, and whiskers indicate the 90% confidence interval. Stars indicate a significant difference between peaks with and without an oCGI ($q < 0.05$, Wilcoxon rank-sum test, BH-corrected). (B) Average bigWig signal (analogous to RPKM) for peaks with and without an oCGI from developing tissues. Box plots show the interquartile range and median, and whiskers indicate the 90% confidence interval. Stars indicate a significant difference between peaks with and without an oCGI ($q < 0.05$, Wilcoxon rank-sum test, BH-corrected).

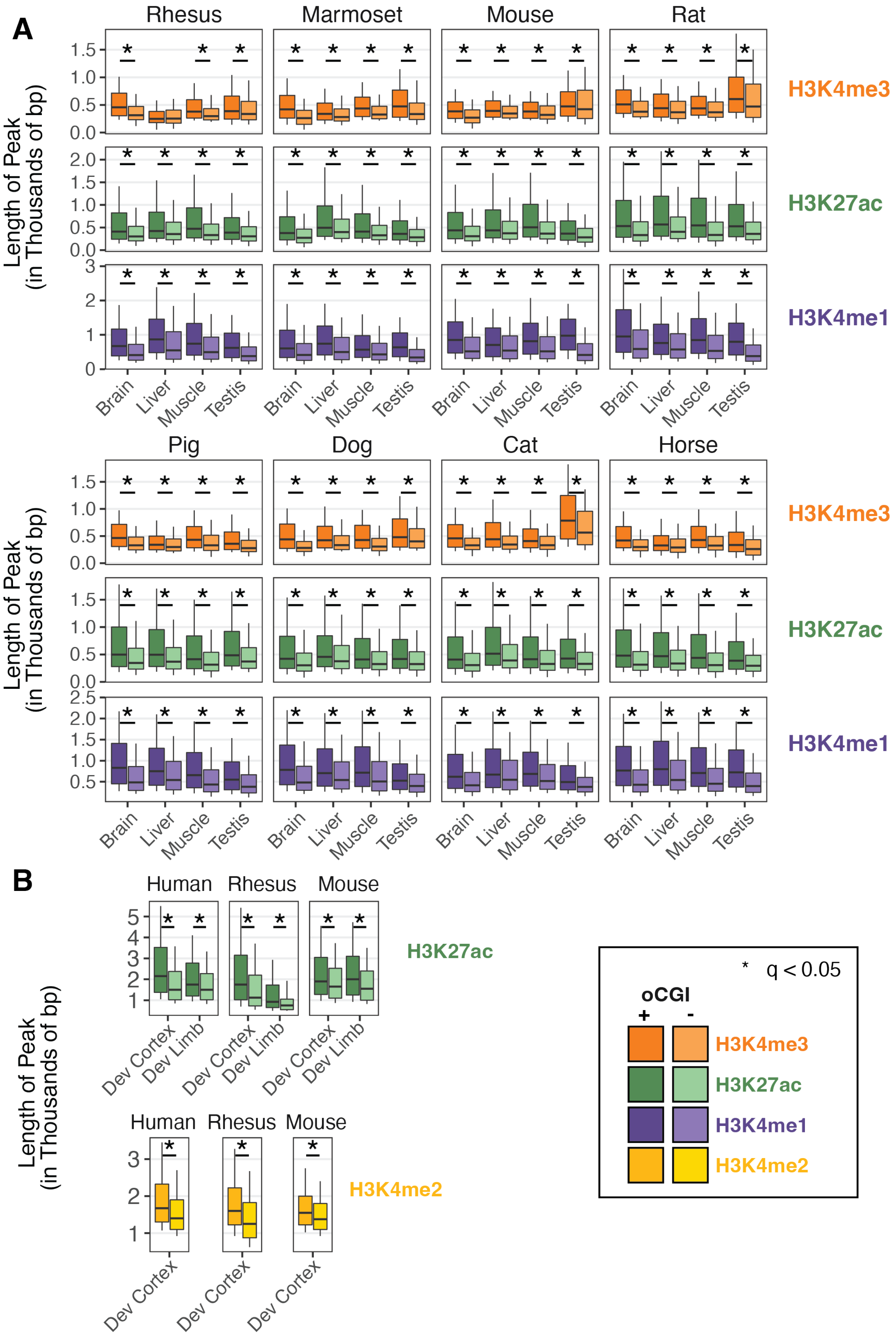

A
Rhesus
Marmoset
Mouse
Rat
Pig
Dog
Cat
Horse
Length of Peak
(in Thousands of bp)
H3K4me3
H3K27ac
H3K4me1
Brain
Liver
Muscle
Testis
B
Human
Rhesus
Mouse
Dev Cortex
Dev Limb
H3K4me2
* q < 0.05
oCGI
+
-

**Figure S6. Histone modification peaks with oCGIs are longer**

(A) Length of peaks with and without an oCGI for each histone modification in adult tissues. Box plots show the interquartile range and median, and whiskers indicate the 90% confidence interval. Stars indicate a significant difference between peaks with and without an oCGI ($q < 0.05$, Wilcoxon rank-sum test, BH-corrected). (B) Length of peaks with and without an oCGI for each histone modification in developing tissues. Box plots show the interquartile range and median, and whiskers indicate the 90% confidence interval. Stars indicate a significant difference between peaks with and without an oCGI ($q < 0.05$, Wilcoxon rank-sum test, BH-corrected).

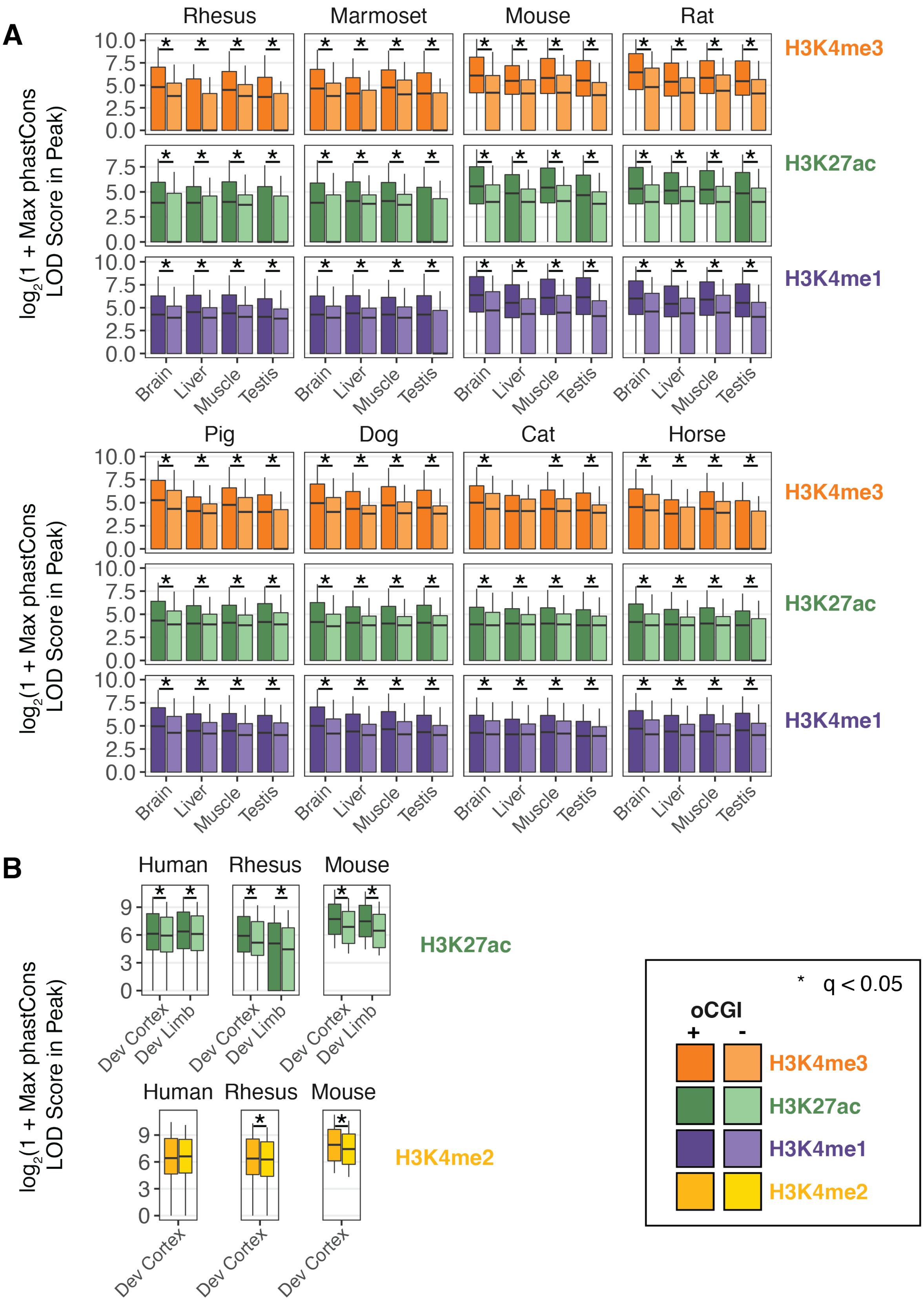
A
Rhesus
Marmoset
Mouse
Rat
Pig
Dog
Cat
Horse
log2(1 + Max phastCons LOD Score in Peak)
H3K4me3
H3K27ac
H3K4me1
Brain
Liver
Muscle
Testis
B
Human
Rhesus
Mouse
Dev Cortex
Dev Limb
H3K4me2
* q < 0.05
oCGI
+
-

**Figure S7. Histone modification peaks with oCGIs are more constrained**

(A) Maximum phastCons LOD scores in peaks with and without an oCGI from adult tissues. Box plots show the interquartile range and median, and whiskers indicate the 90% confidence interval. Stars indicate a significant difference between peaks with and without an oCGI ($q < 0.05$, Wilcoxon rank-sum test, BH-corrected). (B) Maximum phastCons LOD scores in peaks with and without an oCGI from developing tissues. Box plots show the interquartile range and median, and whiskers indicate the 90% confidence interval. Stars indicate a significant difference between peaks with and without an oCGI ($q < 0.05$, Wilcoxon rank-sum test, BH-corrected).

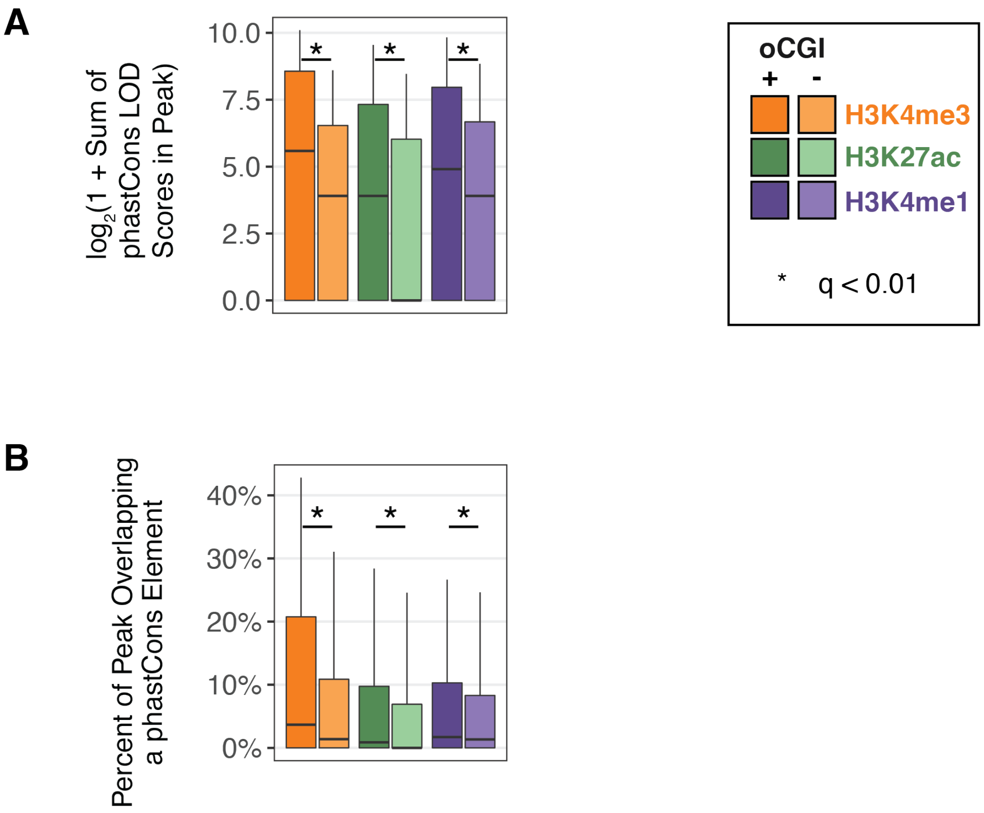


**Figure S8. Alternative measures of evolutionary constraint**

Results are shown for peaks with and without oCGIs for adult brain in rhesus. (A) Sum of phastCons LOD scores within histone modification peaks with and without an oCGI. Box plots show the interquartile range and median, and whiskers indicate the 90% confidence interval. Stars indicate a significant difference between peaks with and without an oCGI ($q < 0.05$, Wilcoxon rank-sum test, BH-corrected). (B) Percentage of bases in peak with and without an oCGI that overlap a phastCons element. Box plots show the interquartile range and median, and whiskers indicate the 90% confidence interval. Stars indicate a significant difference between peaks with and without an oCGI ($q < 0.05$, Wilcoxon rank-sum test, BH-corrected).

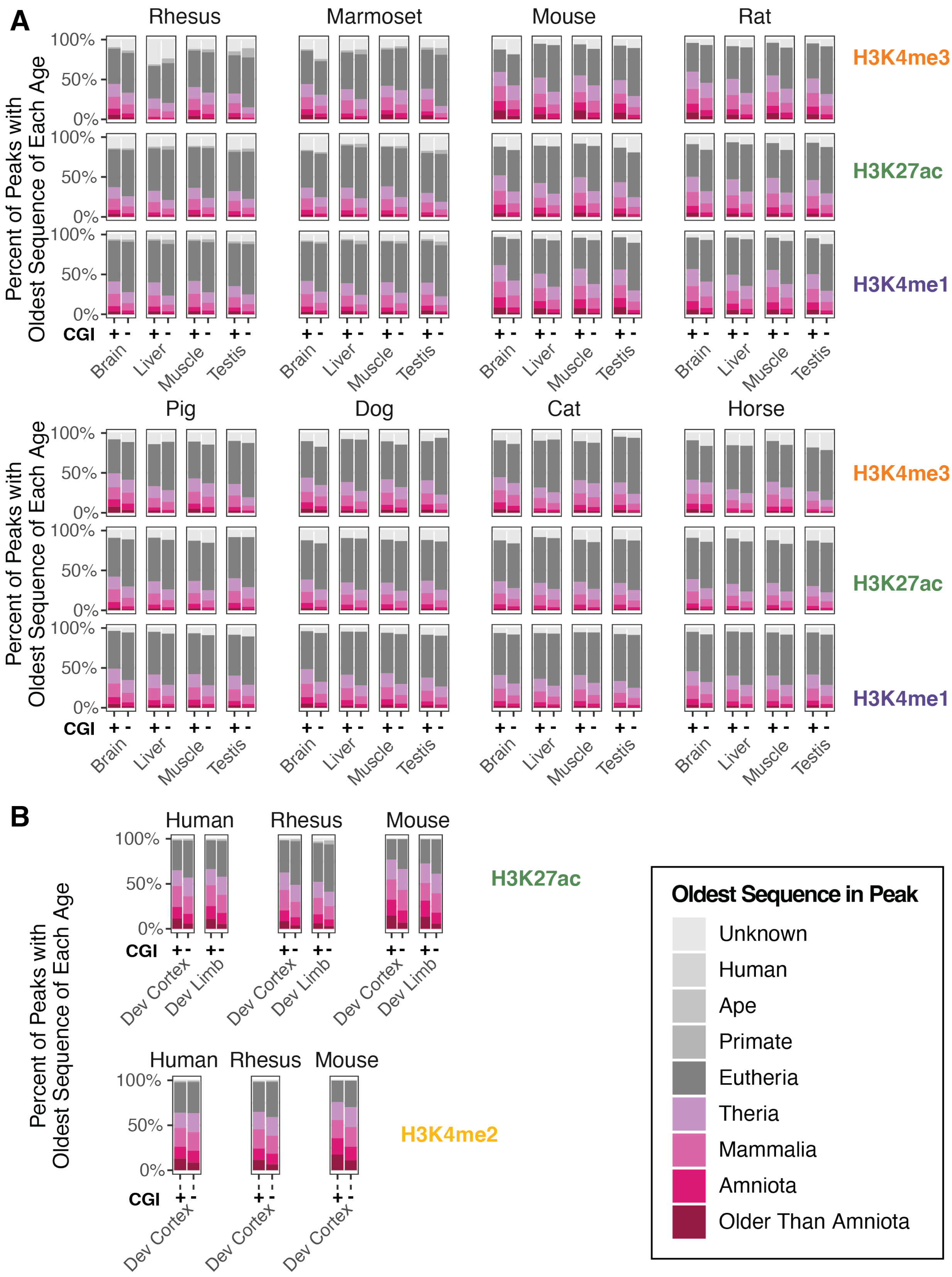
A
Rhesus
Marmoset
Mouse
Rat
Pig
Dog
Cat
Horse
Percent of Peaks with Oldest Sequence of Each Age
100%
50%
0%
H3K4me3
H3K27ac
H3K4me1
CGI
+ -
Brain
Liver
Muscle
Testis
B
Human
Rhesus
Mouse
H3K27ac
Dev Cortex
Dev Limb
H3K4me2
Oldest Sequence in Peak
Unknown
Human
Ape
Primate
Eutheria
Theria
Mammalia
Amniota
Older Than Amniota

**Figure S9. Histone modification peaks with oCGIs are present at older sequences**
(A) The percentage of peaks with and without an oCGI from adult tissues whose oldest sequence belongs to each age category (see legend). (B) The percentage of peaks with and without an oCGI from developing tissues whose oldest sequence belongs to each age category (see legend).

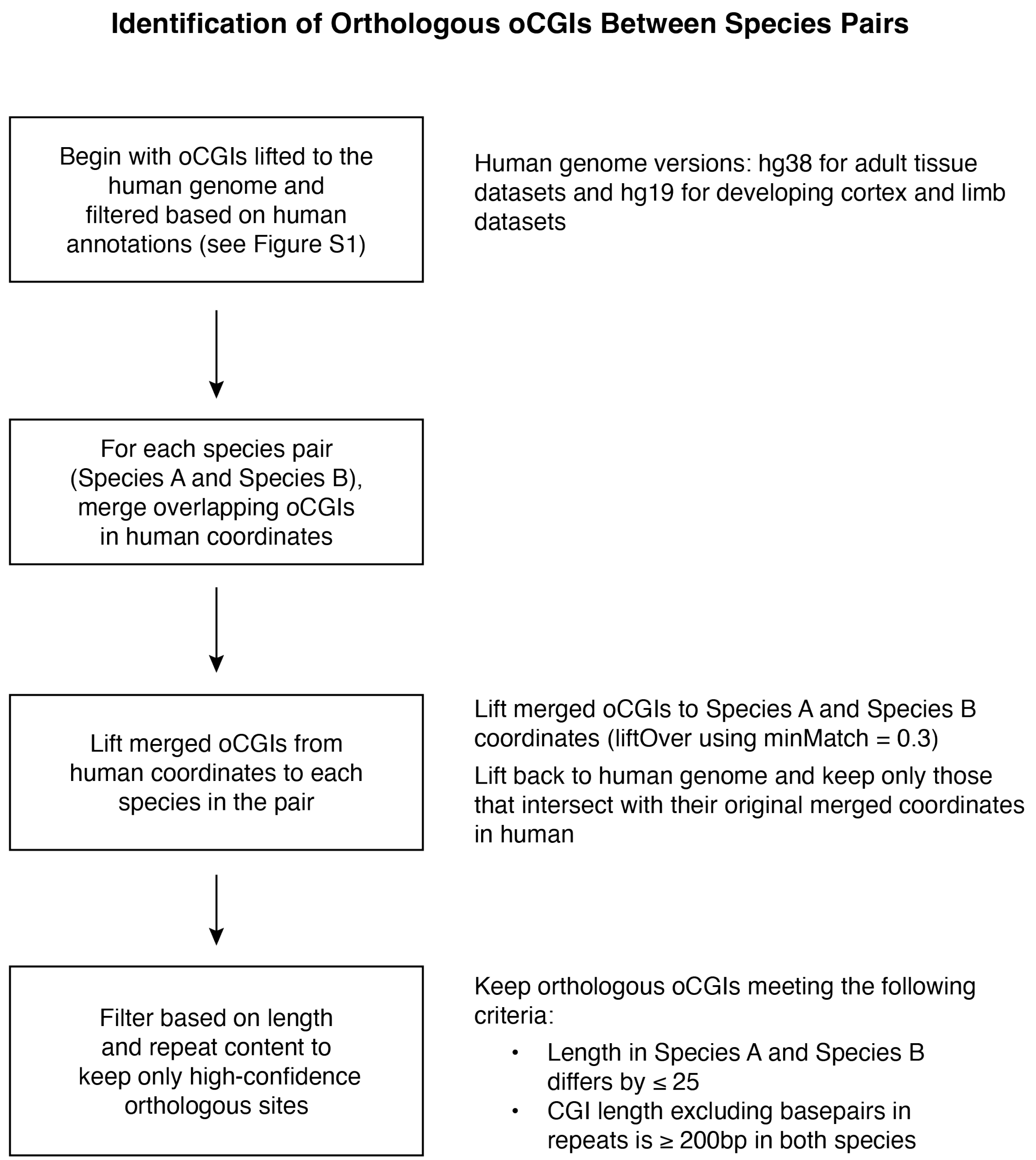


**Figure S10. Pipeline for identifying orthologous oCGIs between species pairs**
Flow chart of steps involved in identifying orthologous oCGIs between species pairs (*left*) and details of each step (*right*).

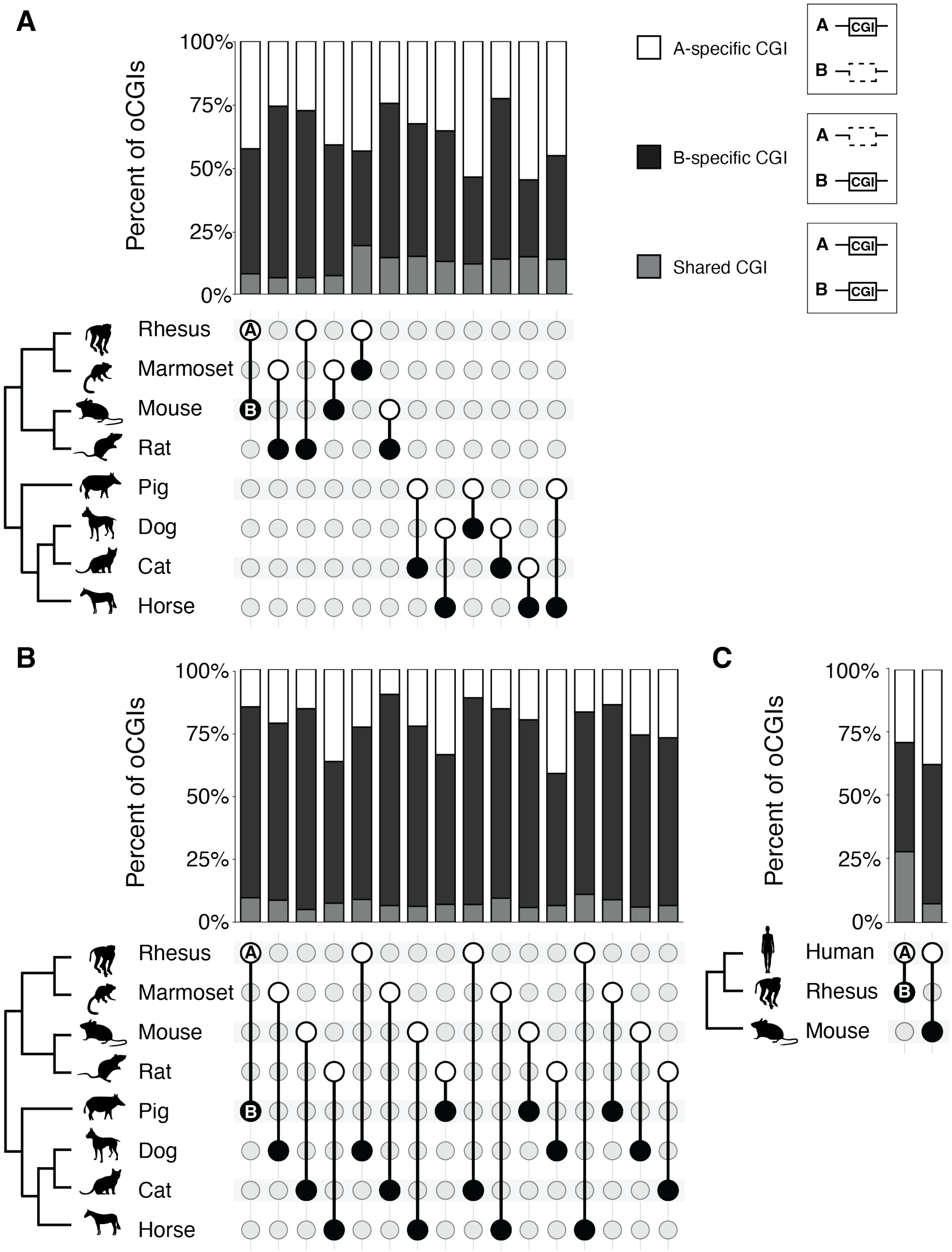
A
B
C
Percent of oCGIs
100%
75%
50%
25%
0%
A-specific CGI
B-specific CGI
Shared CGI
CGI
Human
Rhesus
Marmoset
Mouse
Rat
Pig
Dog
Cat
Horse

**Figure S11. Divergence of oCGIs across all species pairs**
Percent of oCGIs across species pairs (species A vs species B) that are A-only, B-only, or shared. In all cases species A is above species B on the phylogeny, with species A denoted by a white circle and species B denoted by a black circle. (A) More closely related species pairs. (B) More distantly related species pairs. (C) Species pairs involving human.

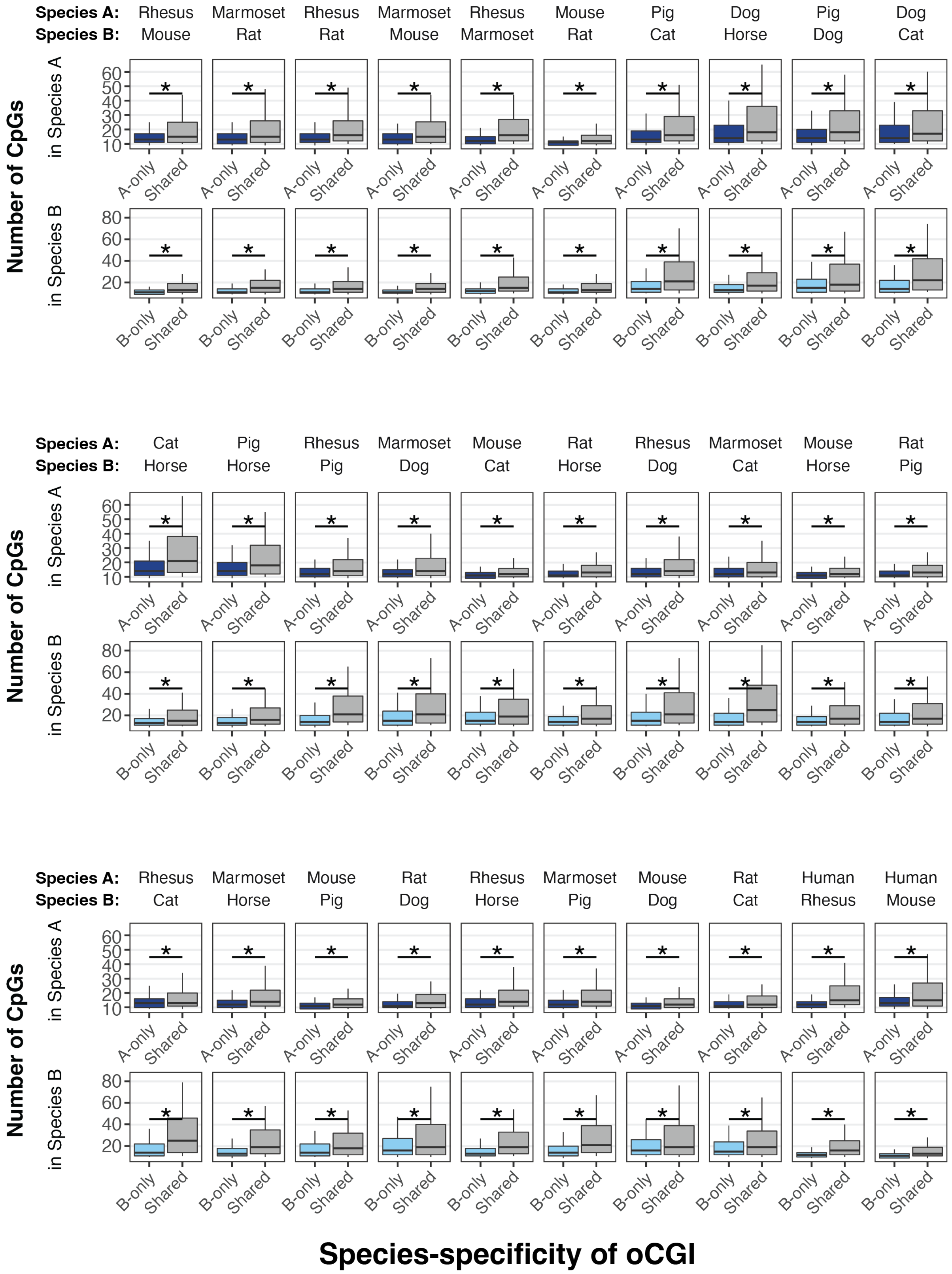
Species A:
Species B:
Rhesus Mouse
Marmoset Rat
Rhesus Rat
Marmoset Mouse
Rhesus Marmoset
Mouse Rat
Pig Cat
Dog Horse
Pig Dog
Dog Cat
Cat Horse
Pig Horse
Rhesus Pig
Marmoset Dog
Mouse Cat
Rat Horse
Rhesus Dog
Marmoset Cat
Mouse Horse
Rat Pig
Rhesus Cat
Marmoset Horse
Mouse Pig
Rat Dog
Rhesus Horse
Marmoset Pig
Mouse Dog
Rat Cat
Human Rhesus
Human Mouse
Number of CpGs
in Species A
in Species B
A-only
B-only
Shared
Species-specificity of oCGI

**Figure S12. Shared oCGIs contain more CpG dinucleotides**

Number of CpG dinucleotides in A-only (dark blue) or B-only (light blue) oCGIs compared to shared (gray) oCGIs across all species pairs considered. Box plots show the interquartile range and median, and whiskers indicate the 90% confidence interval. Stars indicate a significant difference between species-specific and shared oCGIs ($q < 0.05$, Wilcoxon rank-sum test, BH-corrected).

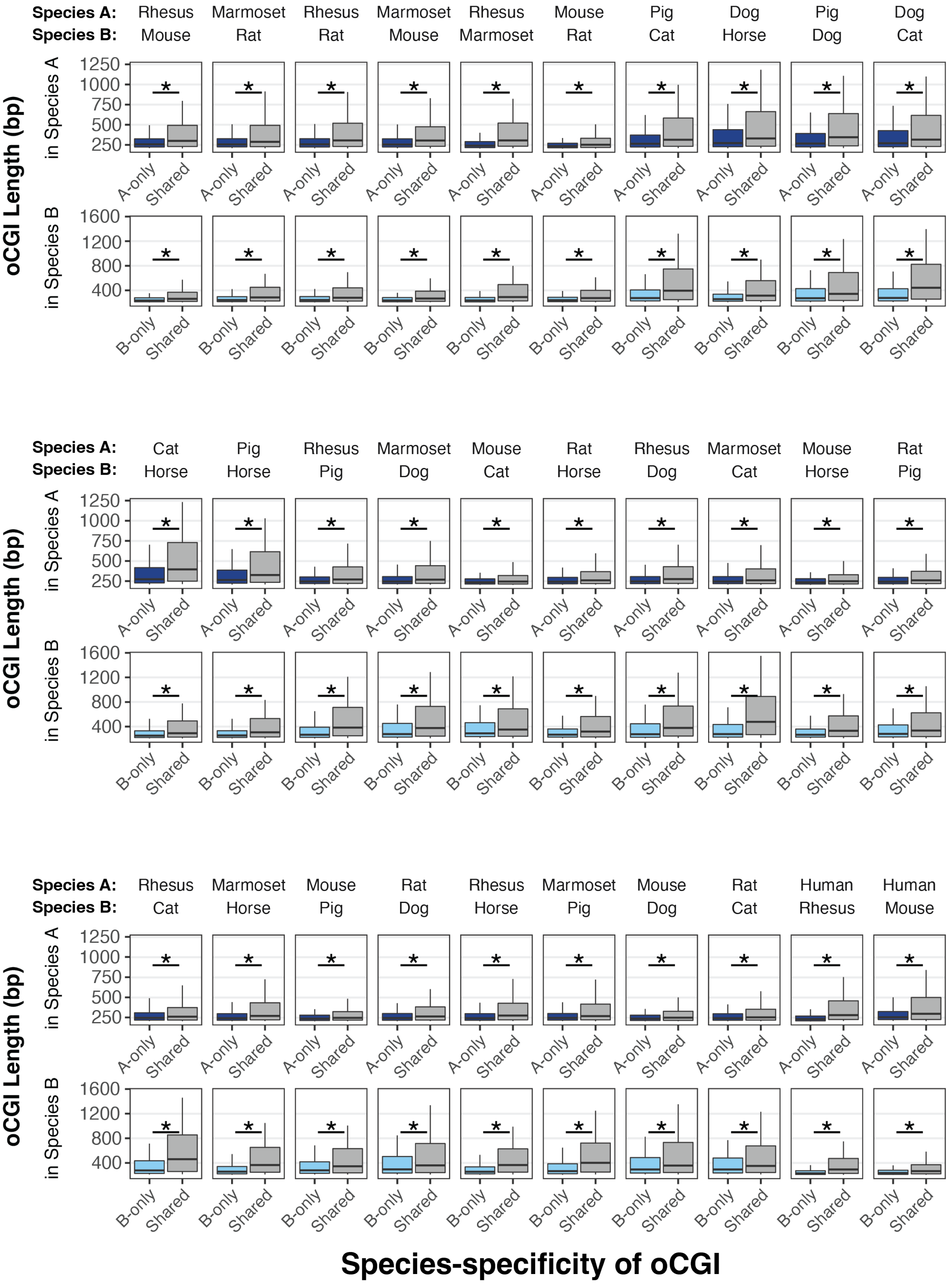
Species A:
Species B:
Rhesus Mouse
Marmoset Rat
Rhesus Rat
Marmoset Mouse
Rhesus Marmoset
Mouse Rat
Pig Cat
Dog Horse
Pig Dog
Dog Cat
Cat Horse
Pig Horse
Rhesus Pig
Marmoset Dog
Mouse Cat
Rat Horse
Rhesus Dog
Marmoset Cat
Mouse Horse
Rat Pig
Rhesus Cat
Marmoset Horse
Mouse Pig
Rat Dog
Rhesus Horse
Marmoset Pig
Mouse Dog
Rat Cat
Human Rhesus
Human Mouse
oCGI Length (bp)
in Species A
in Species B
1250
1000
750
500
250
1600
1200
800
400
*
A-only
B-only
Shared
Species-specificity of oCGI

**Figure S13. Shared oCGIs are longer**

Length of A-only (dark blue) or B-only (light blue) oCGIs compared to shared (gray) oCGIs across all species pairs considered. Box plots show the interquartile range and median, and whiskers indicate the 90% confidence interval. Stars indicate a significant difference between species-specific and shared oCGIs ($q < 0.05$, Wilcoxon rank-sum test, BH-corrected).

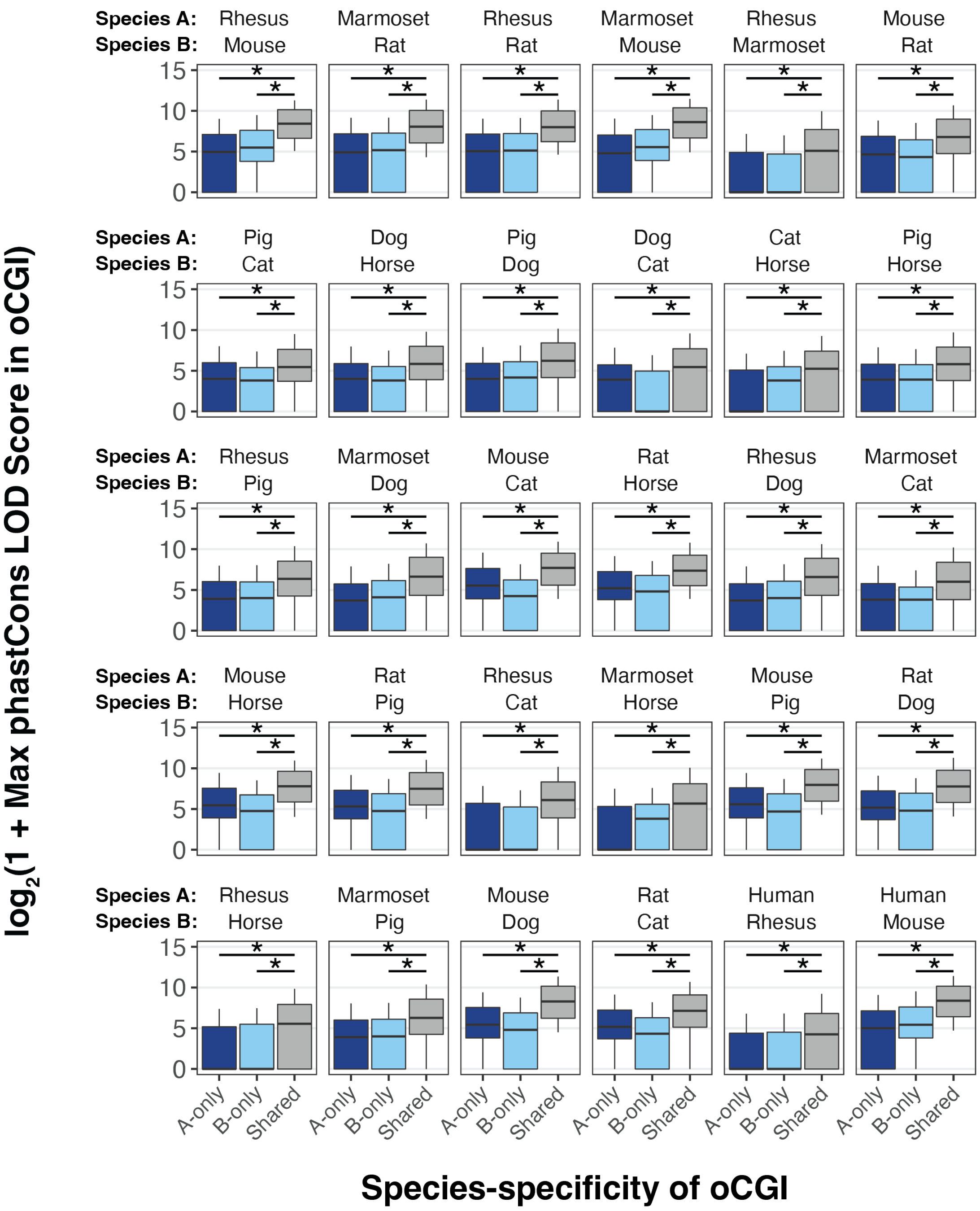


**Figure S14. Shared oCGIs have higher maximum phastCons LOD scores**
Maximum phastCons LOD scores in A-only (dark blue), B-only (light blue), and shared (gray) oCGIs across all species pairs considered. Box plots show the interquartile range and median, and whiskers indicate the 90% confidence interval. Stars indicate a significant difference between species-specific and shared oCGIs ($q < 0.05$, Wilcoxon rank-sum test, BH-corrected).

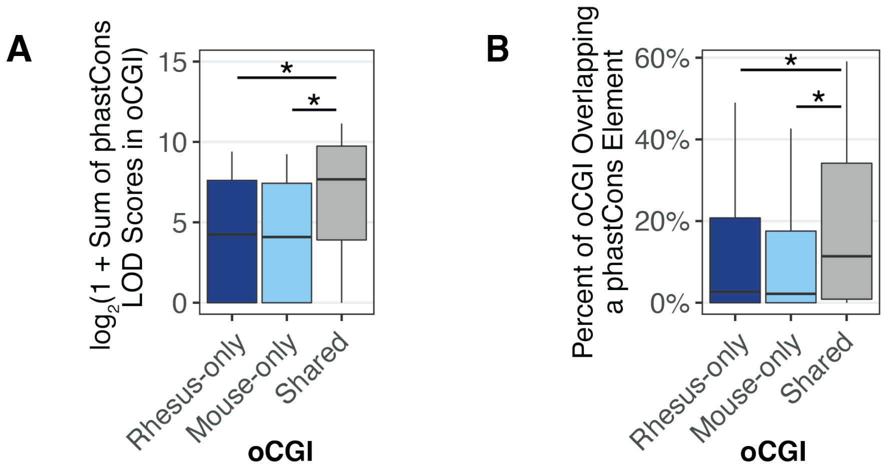


**Figure S15. Alternative measures of oCGI constraint**

Results are shown for the comparison between rhesus (species A) versus mouse (species B). (A) Sum of phastCons LOD scores within oCGIs that are rhesus-only, mouse-only, or shared. (B) Percent of bases that overlap a phastCons element within oCGIs that are rhesus-only, mouse-only, or shared. Box plots show the interquartile range and median, and whiskers indicate the 90% confidence interval. Stars indicate a significant difference between species-specific and shared oCGIs ($q < 0.05$, Wilcoxon rank-sum test, BH-corrected).

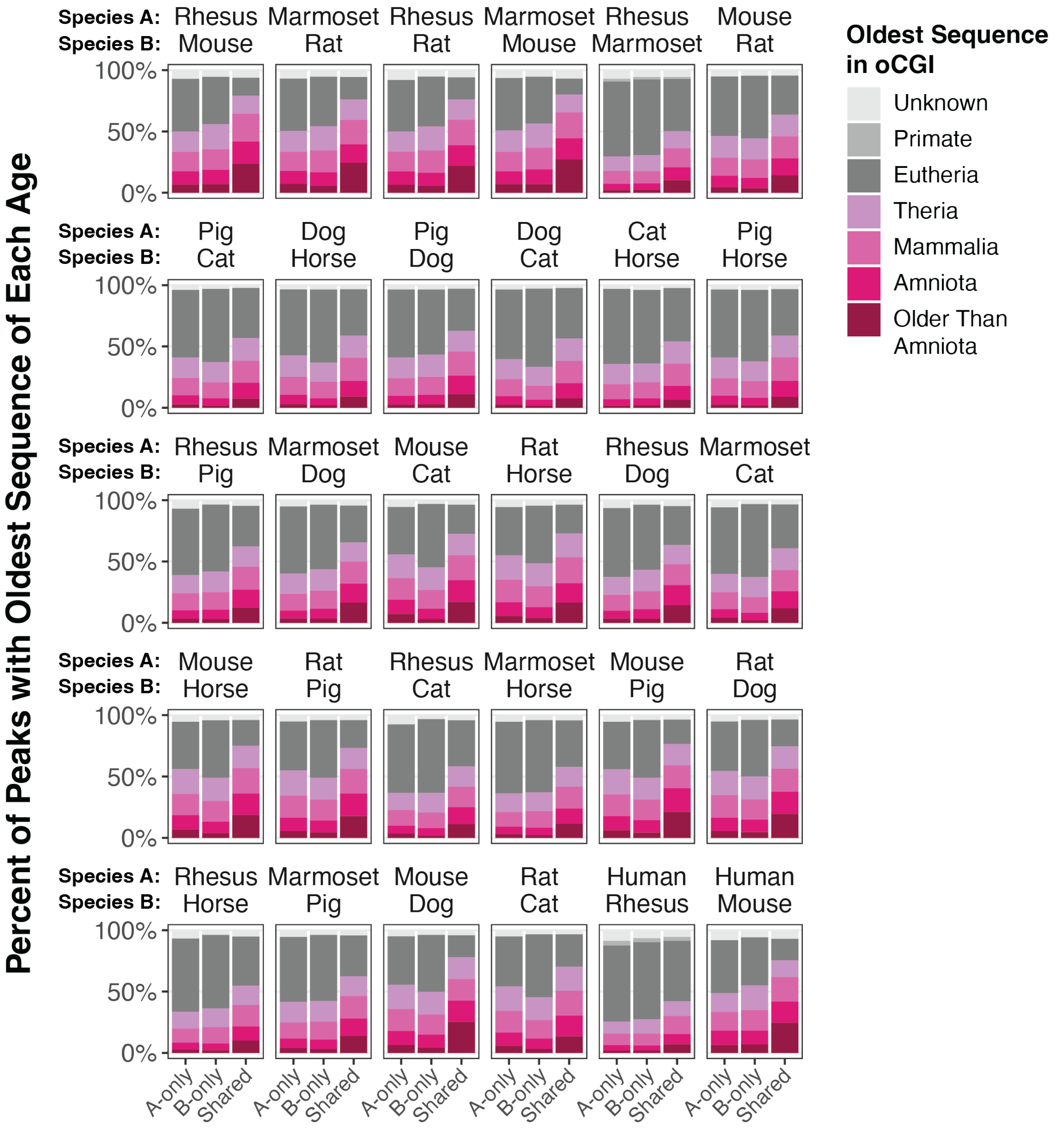


**Figure S16. Shared oCGIs are present at older sequences**
The percentage of A-only, B-only, and shared oCGIs whose oldest sequence belongs to each age category (see legend), across all species pairs considered.

A

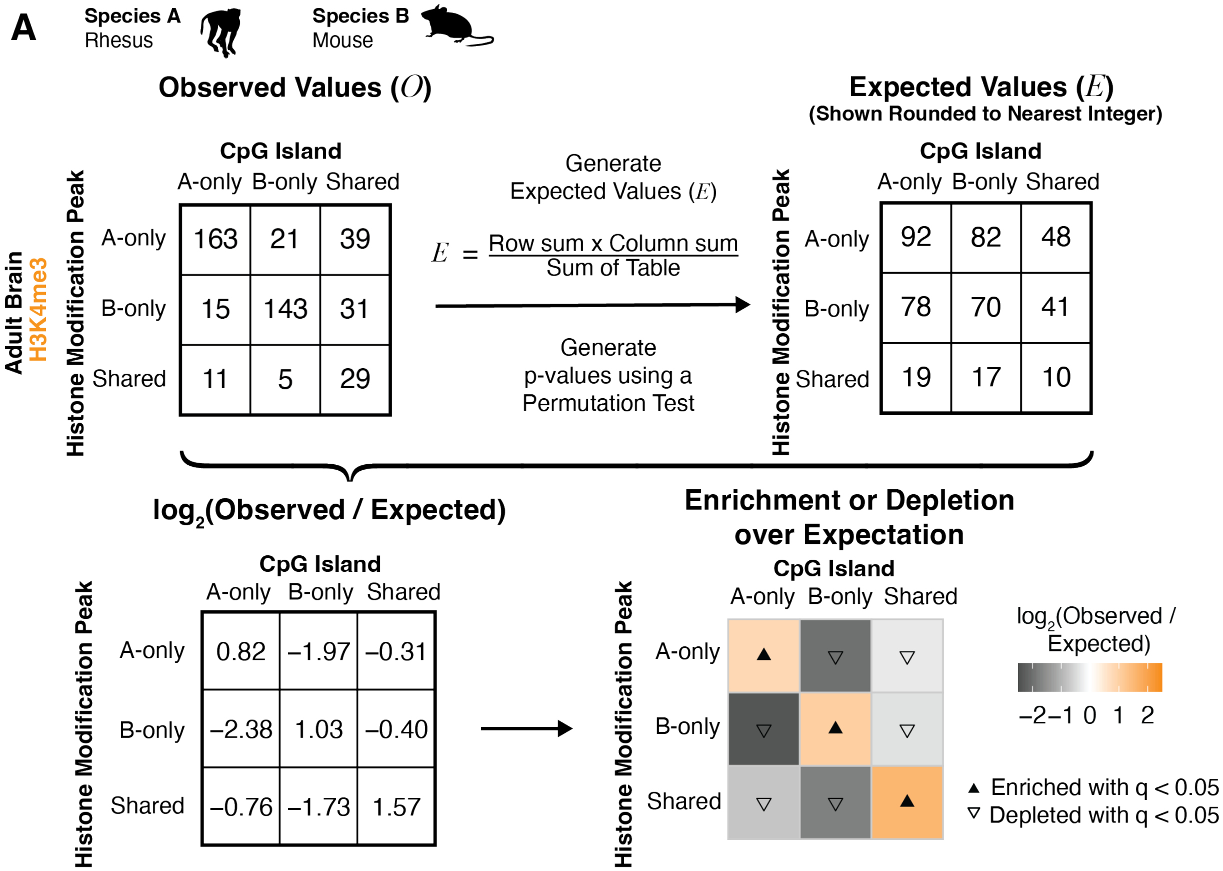


B

#### Permutation Test

1. Calculate the Expected Value, $E$, in each box.

$$E = \frac{\text{Row sum x Column sum}}{\text{Sum of Table}}$$

2. Using all orthologous oCGIs in the dataset, randomly reassign labels for species-specificity of the oCGI (A-only, B-only, Shared).

3. Sort the oCGIs with permuted labels into a new 3 x 3 box. Count the number of sites falling into each box.

4. Repeat $n$ = 20,000 times to establish a distribution of permuted values in each box.

5. Calculate p-value

   Number of times the permuted value is more extreme than the Observed Value $O$, divided by the number of resamples, $n$.

   If this number is 0, then p = 1 / $n$.

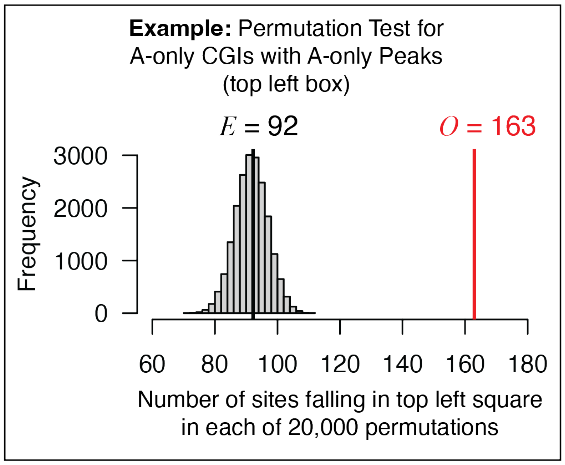

**Figure S17. Permutation test used for enrichment analysis**

(A) Determination of enrichment and depletion for each indicated comparison of species-specific and shared oCGIs and species-specific and shared peaks (rhesus macaque versus mouse, H3K4me3 peaks in adult brain). Observed Values and Expected Values (*top*) are used to generate $\log_2$-transformed ratios (*bottom left*), which are subsequently visualized as color gradients with each box colored according to the level of enrichment (orange) or depletion (gray) (*bottom right*) of genome-wide sites meeting the criteria for that box. The color bar illustrates the level of enrichment or depletion over expectation. The filled upward-pointing triangles denote significant enrichment and open downward-pointing triangles denote significant depletion ($q < 0.05$, permutation test, BH-corrected). (B) Explanation of permutation test to calculate statistical significance of the enrichment or depletion. The panel at right shows the distribution of values from 20,000 permutations for the top left box of the grid (A-only oCGIs with A-only peaks). The Observed Value of 163 (shown in red) is enriched compared to the Expected Value of 92, which is the mean of the 20,000 permuted values.

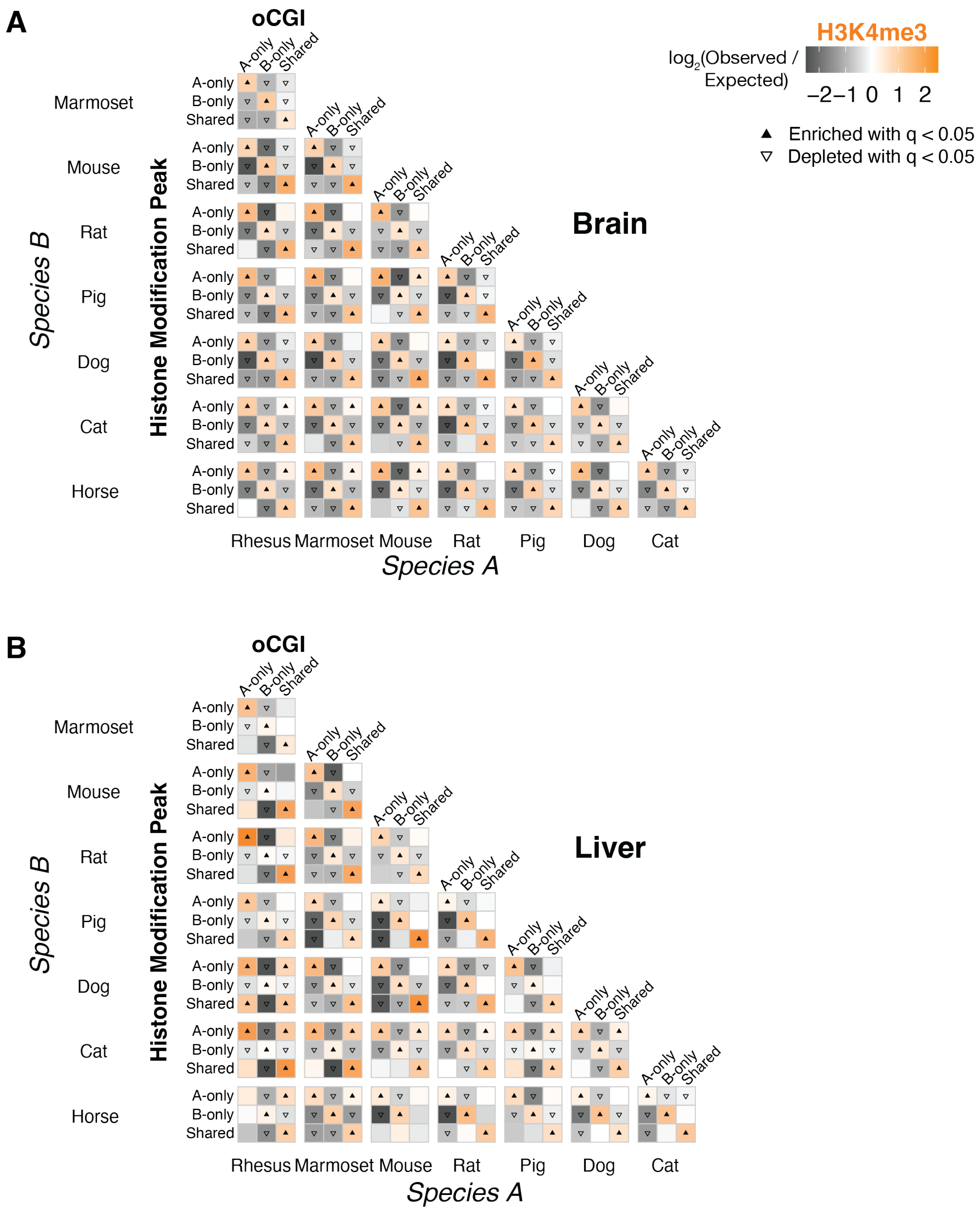
A
oCGI
H3K4me3
log2(Observed / Expected)
−2 −1 0 1 2
▲ Enriched with q < 0.05
▽ Depleted with q < 0.05
Brain
A-only
B-only
Shared
Marmoset
Mouse
Rat
Pig
Dog
Cat
Horse
Species B
Histone Modification Peak
Rhesus Marmoset Mouse Rat Pig Dog Cat
Species A
B
oCGI
Liver
Species B
Histone Modification Peak
Species A

**Figure S18. Species with oCGI predicts species with H3K4me3 peak (brain and liver)**
Enrichment and depletion in each indicated comparison of species-specific and shared oCGIs (*top:* A-only, B-only, Shared) and species-specific and shared peaks (*left*: A-only, B-only, Shared), compared to a null expectation of no association between oCGI turnover and peak turnover. Each 3 x 3 grid shows the results for a specific test examining oCGIs and their overlap with H3K4me3 in adult brain (A) or liver (B) within a given species pair. Each box in each grid is colored according to the level of enrichment over expectation (orange) or depletion (gray) of genome-wide sites that meet the criteria for that box. The color bar illustrates the level of enrichment or depletion over expectation. For visualization, values greater than 2.5 and less than -2.5 were set to 2.5 and -2.5, respectively. The filled upward-pointing triangles denote significant enrichment and open downward-pointing triangles denote significant depletion ($q < 0.05$, permutation test, BH-corrected, see Fig. S17 and Methods).

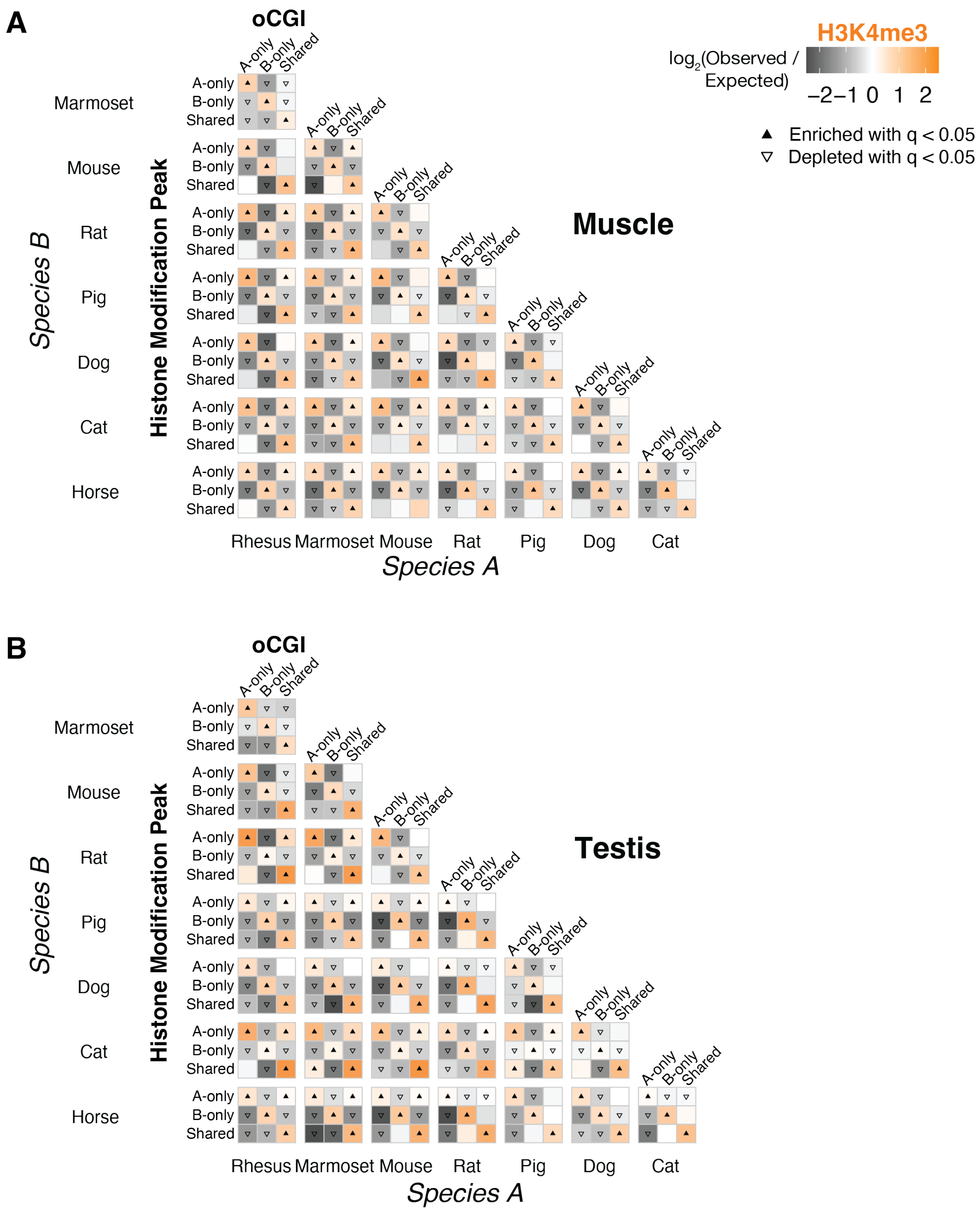
A
oCGI
H3K4me3
log2(Observed / Expected)
−2 −1 0 1 2
Enriched with q < 0.05
Depleted with q < 0.05
Muscle
Species B
Histone Modification Peak
Marmoset
Mouse
Rat
Pig
Dog
Cat
Horse
A-only
B-only
Shared
Rhesus
Marmoset
Mouse
Rat
Pig
Dog
Cat
Species A
B
oCGI
Testis
Species B
Histone Modification Peak
Species A

**Figure S19. Species with oCGI predicts species with H3K4me3 peak (muscle and testis)**
Enrichment and depletion in each indicated comparison of species-specific and shared oCGIs (*top:* A-only, B-only, Shared) and species-specific and shared peaks (*left*: A-only, B-only, Shared), compared to a null expectation of no association between oCGI turnover and peak turnover. Each 3 x 3 grid shows the results for a specific test examining oCGIs and their overlap with H3K4me3 in adult muscle (A) or testis (B) within a given species pair. Each box in each grid is colored according to the level of enrichment over expectation (orange) or depletion (gray) of genome-wide sites that meet the criteria for that box. The color bar illustrates the level of enrichment or depletion over expectation. For visualization, values greater than 2.5 and less than -2.5 were set to 2.5 and -2.5, respectively. The filled upward-pointing triangles denote significant enrichment and open downward-pointing triangles denote significant depletion ($q < 0.05$, permutation test, BH-corrected, see Fig. S17 and Methods).

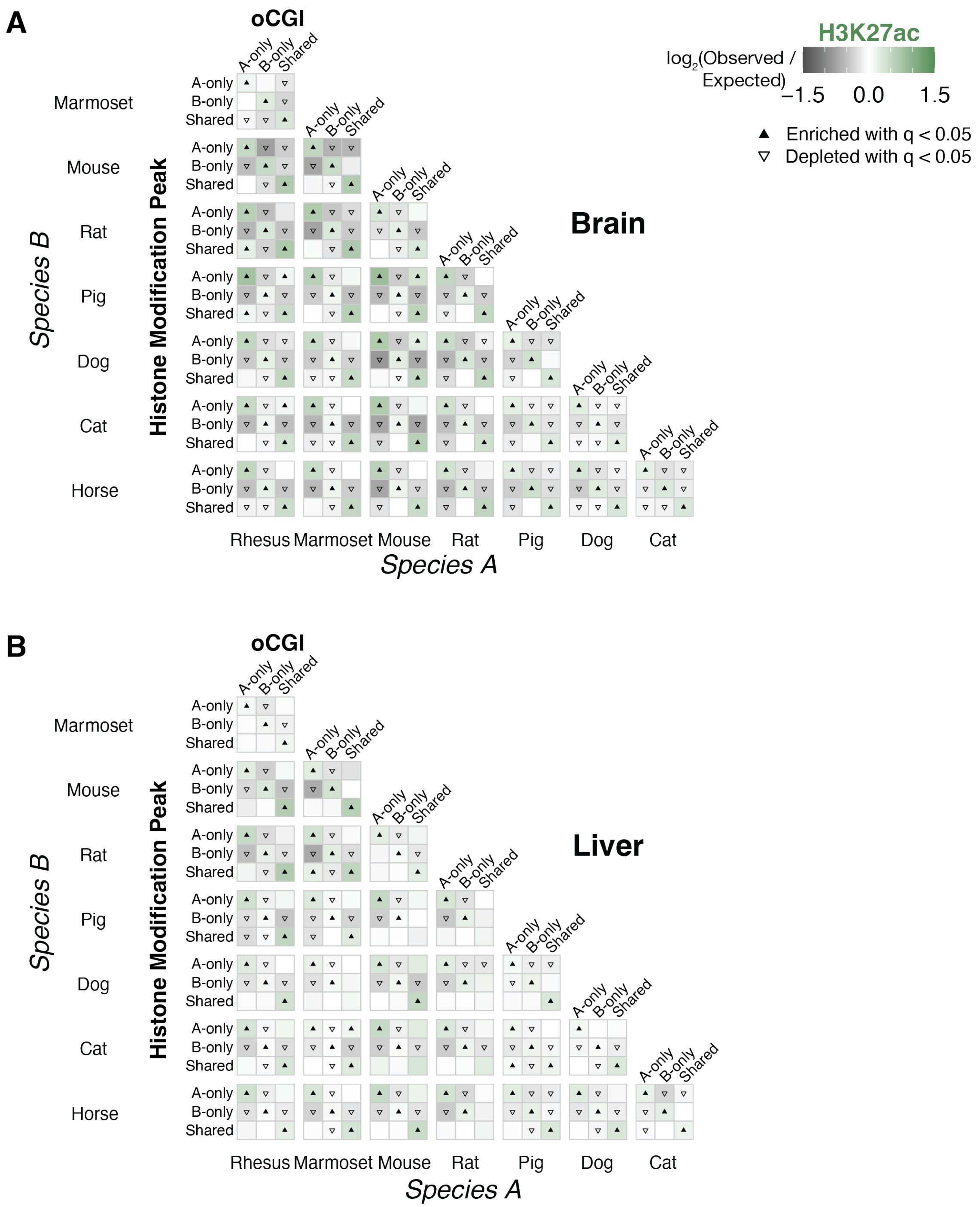
A
oCGI
H3K27ac
log2(Observed / Expected)
−1.5
0.0
1.5
Enriched with q < 0.05
Depleted with q < 0.05
Brain
Species B
Histone Modification Peak
Marmoset
Mouse
Rat
Pig
Dog
Cat
Horse
A-only
B-only
Shared
Rhesus Marmoset Mouse Rat Pig Dog Cat
Species A
B
oCGI
Liver
Species B
Histone Modification Peak
Species A

**Figure S20. Species with oCGI predicts species with H3K27ac peak (brain and liver)**

Enrichment and depletion in each indicated comparison of species-specific and shared oCGIs (*top:* A-only, B-only, Shared) and species-specific and shared peaks (*left*: A-only, B-only, Shared), compared to a null expectation of no association between oCGI turnover and peak turnover. Each 3 x 3 grid shows the results for a specific test examining oCGIs and their overlap with H3K27ac in adult brain (A) or liver (B) within a given species pair. Each box in each grid is colored according to the level of enrichment over expectation (green) or depletion (gray) of genome-wide sites that meet the criteria for that box. The color bar illustrates the level of enrichment or depletion over expectation. For visualization, values greater than 1.5 and less than -1.5 were set to 1.5 and -1.5, respectively. The filled upward-pointing triangles denote significant enrichment and open downward-pointing triangles denote significant depletion ($q < 0.05$, permutation test, BH-corrected, see Fig. S17 and Methods).

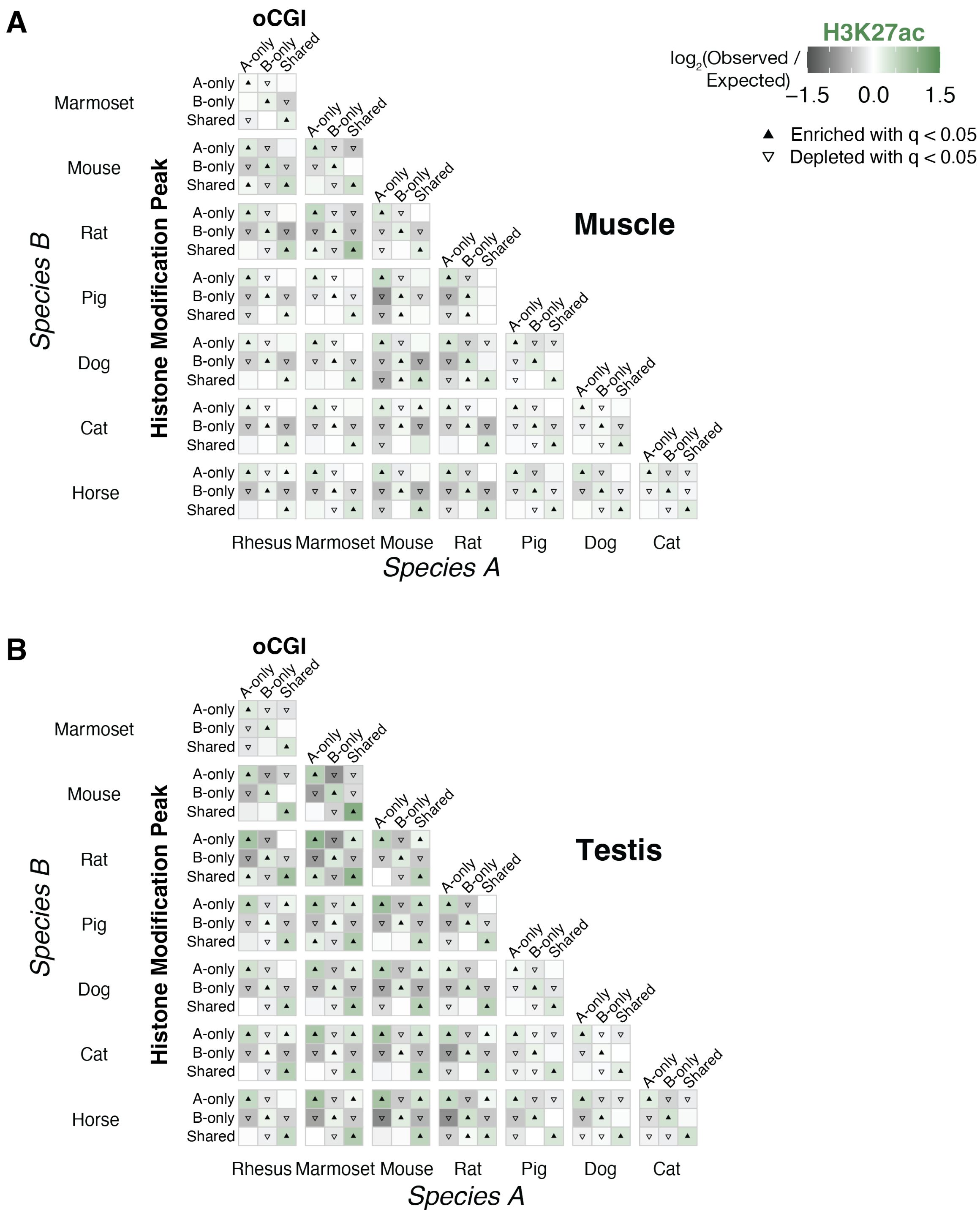
A
oCGI
H3K27ac
log2(Observed / Expected)
−1.5 0.0 1.5
Enriched with q < 0.05
Depleted with q < 0.05
Muscle
Species B
Histone Modification Peak
Marmoset
Mouse
Rat
Pig
Dog
Cat
Horse
A-only
B-only
Shared
Rhesus Marmoset Mouse Rat Pig Dog Cat
Species A
B
oCGI
Testis
Species B
Histone Modification Peak
Marmoset
Mouse
Rat
Pig
Dog
Cat
Horse
Rhesus Marmoset Mouse Rat Pig Dog Cat
Species A

**Figure S21. Species with oCGI predicts species with H3K27ac peak (muscle and testis)**
Enrichment and depletion in each indicated comparison of species-specific and shared oCGIs (*top:* A-only, B-only, Shared) and species-specific and shared peaks (*left*: A-only, B-only, Shared), compared to a null expectation of no association between oCGI turnover and peak turnover. Each 3 x 3 grid shows the results for a specific test examining oCGIs and their overlap with H3K27ac in adult muscle (A) or testis (B) within a given species pair. Each box in each grid is colored according to the level of enrichment over expectation (green) or depletion (gray) of genome-wide sites that meet the criteria for that box. The color bar illustrates the level of enrichment or depletion over expectation. For visualization, values greater than 1.5 and less than -1.5 were set to 1.5 and -1.5, respectively. The filled upward-pointing triangles denote significant enrichment and open downward-pointing triangles denote significant depletion ($q < 0.05$, permutation test, BH-corrected, see Fig. S17 and Methods).

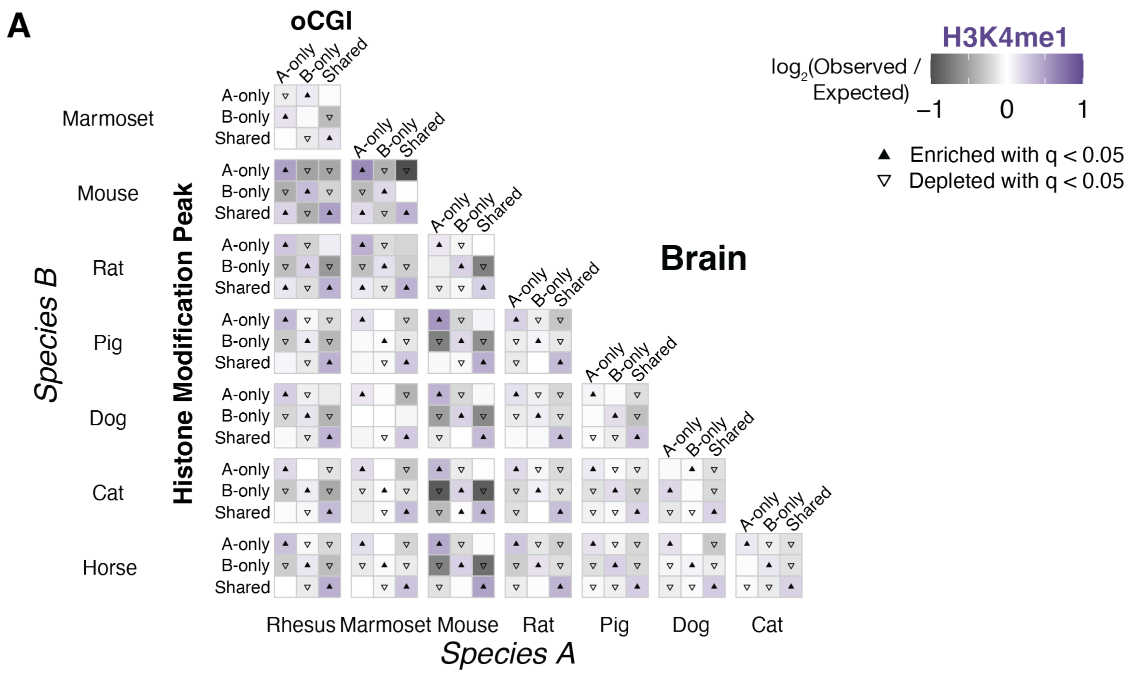
A
oCGI
H3K4me1
log2(Observed / Expected)
−1
0
1
Enriched with q < 0.05
Depleted with q < 0.05
Brain
A-only
B-only
Shared
Marmoset
Mouse
Rat
Pig
Dog
Cat
Horse
Species B
Histone Modification Peak
Rhesus
Marmoset
Mouse
Rat
Pig
Dog
Cat
Species A

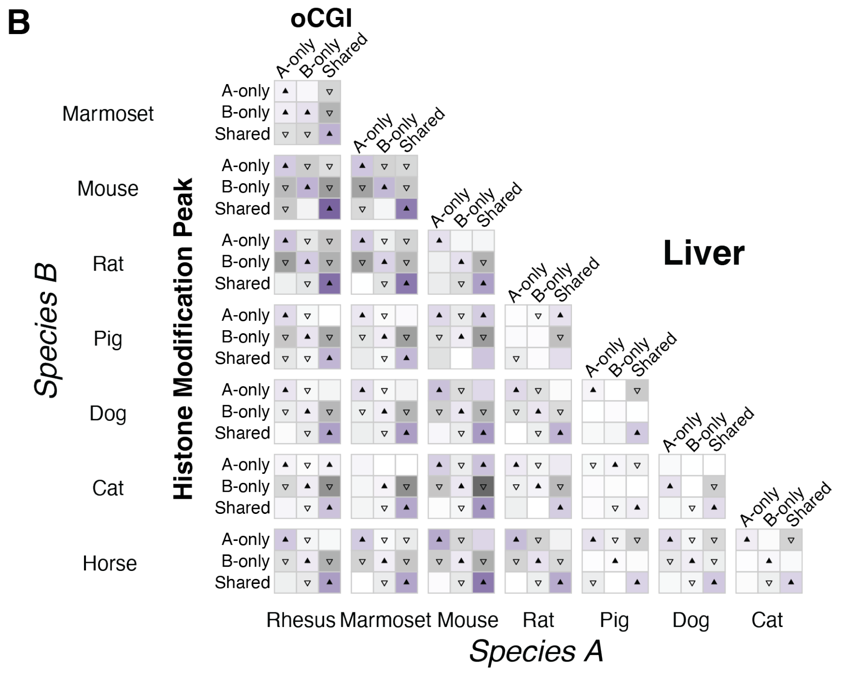
B
oCGI
Liver
A-only
B-only
Shared
Marmoset
Mouse
Rat
Pig
Dog
Cat
Horse
Species B
Histone Modification Peak
Rhesus
Marmoset
Mouse
Rat
Pig
Dog
Cat
Species A

**Figure S22. Species with oCGI predicts species with H3K4me1 peak (brain and liver)**
Enrichment and depletion in each indicated comparison of species-specific and shared oCGIs (*top:* A-only, B-only, Shared) and species-specific and shared peaks (*left*: A-only, B-only, Shared), compared to a null expectation of no association between oCGI turnover and peak turnover. Each 3 x 3 grid shows the results for a specific test examining oCGIs and their overlap with H3K4me1 in adult brain (A) or liver (B) within a given species pair. Each box in each grid is colored according to the level of enrichment over expectation (purple) or depletion (gray) of genome-wide sites that meet the criteria for that box. The color bar illustrates the level of enrichment or depletion over expectation. For visualization, values greater than 1 and less than -1 were set to 1 and -1, respectively. The filled upward-pointing triangles denote significant enrichment and open downward-pointing triangles denote significant depletion ($q < 0.05$, permutation test, BH-corrected, see Fig. S17 and Methods).

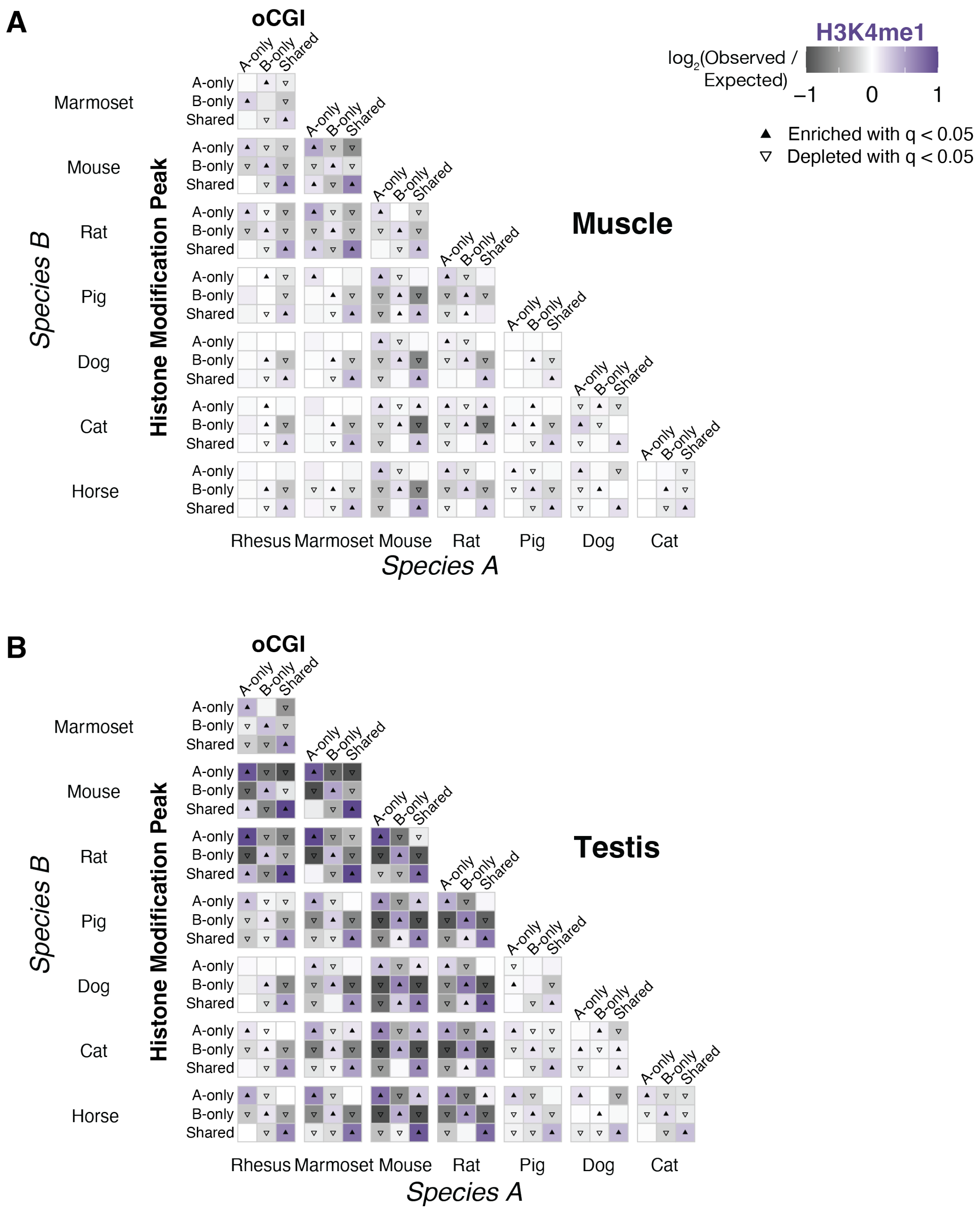
A
B
oCGI
H3K4me1
log2(Observed / Expected)
–1
0
1
Enriched with q < 0.05
Depleted with q < 0.05
Muscle
Testis
A-only
B-only
Shared
Marmoset
Mouse
Rat
Pig
Dog
Cat
Horse
Rhesus
Species A
Species B
Histone Modification Peak

**Figure S23. Species with oCGI predicts species with H3K4me1 peak (muscle and testis)**
Enrichment and depletion in each indicated comparison of species-specific and shared oCGIs (*top:* A-only, B-only, Shared) and species-specific and shared peaks (*left*: A-only, B-only, Shared), compared to a null expectation of no association between oCGI turnover and peak turnover. Each 3 x 3 grid shows the results for a specific test examining oCGIs and their overlap with H3K4me1 in adult muscle (A) or testis (B) within a given species pair. Each box in each grid is colored according to the level of enrichment over expectation (purple) or depletion (gray) of genome-wide sites that meet the criteria for that box. The color bar illustrates the level of enrichment or depletion over expectation. For visualization, values greater than 1 and less than -1 were set to 1 and -1, respectively. The filled upward-pointing triangles denote significant enrichment and open downward-pointing triangles denote significant depletion ($q < 0.05$, permutation test, BH-corrected, see Fig. S17 and Methods).

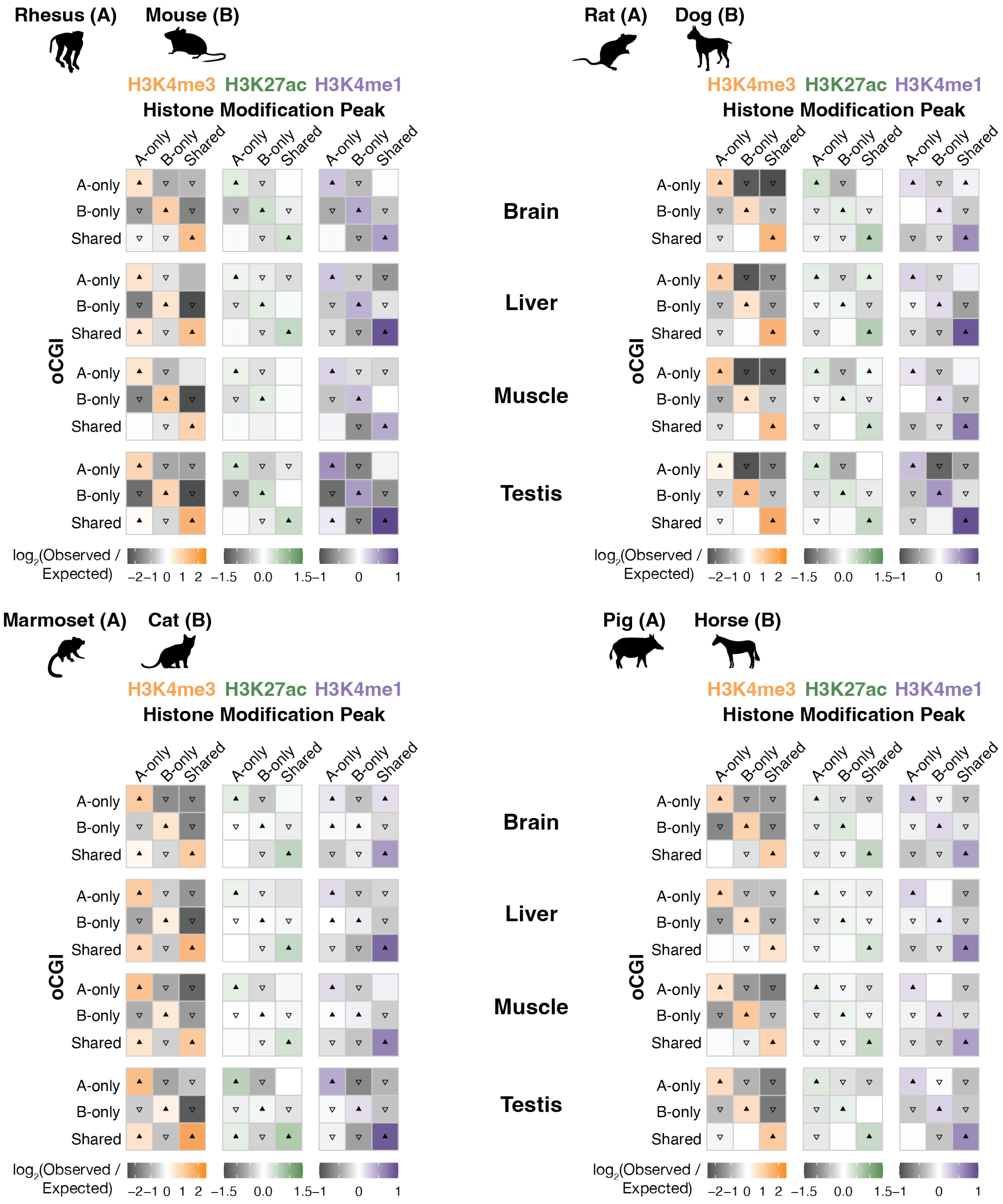

Rhesus (A)
Mouse (B)
Rat (A)
Dog (B)
Marmoset (A)
Cat (B)
Pig (A)
Horse (B)
H3K4me3 H3K27ac H3K4me1
Histone Modification Peak
A-only
B-only
Shared
Brain
Liver
Muscle
Testis
oCGI
log2(Observed / Expected)
−2 −1 0 1 2
−1.5 0.0 1.5
−1 0 1

**Figure S24. Species with peak predicts species with oCGI (peak-centric analysis)**
Enrichment and depletion in each indicated comparison of species-specific and shared peaks (*top:* A-only, B-only, Shared) and species-specific and shared oCGIs (*left*: A-only, B-only, Shared), compared to a null expectation of no association between peak turnover and oCGI turnover. Each 3 x 3 grid shows the results for a specific test examining H3K4me3, H3K27ac, or H3K4me1 peaks and their overlap with oCGIs in adult brain, liver, muscle, or testis within a given species pair. Each box in each grid is colored according to the level of enrichment over expectation (orange for H3K4me3, green for H3K27ac, purple for H3K4me1) or depletion (gray for all marks) of genome-wide sites that meet the criteria for that box. The color bars illustrate the level of enrichment or depletion over expectation. For visualization, values greater than the scale maximum (2.5 for H3K4me3, 1.5 for H3K27ac, and 1 for H3K4me1) and less than the scale minimum (-2.5 for H3K4me3, -1.5 for H3K27ac, and -1 for H3K4me1) were set to the maximum or minimum, respectively. The filled upward-pointing triangles denote significant enrichment and open downward-pointing triangles denote significant depletion ($q < 0.05$, permutation test, BH-corrected, see Fig. S17 and Methods).

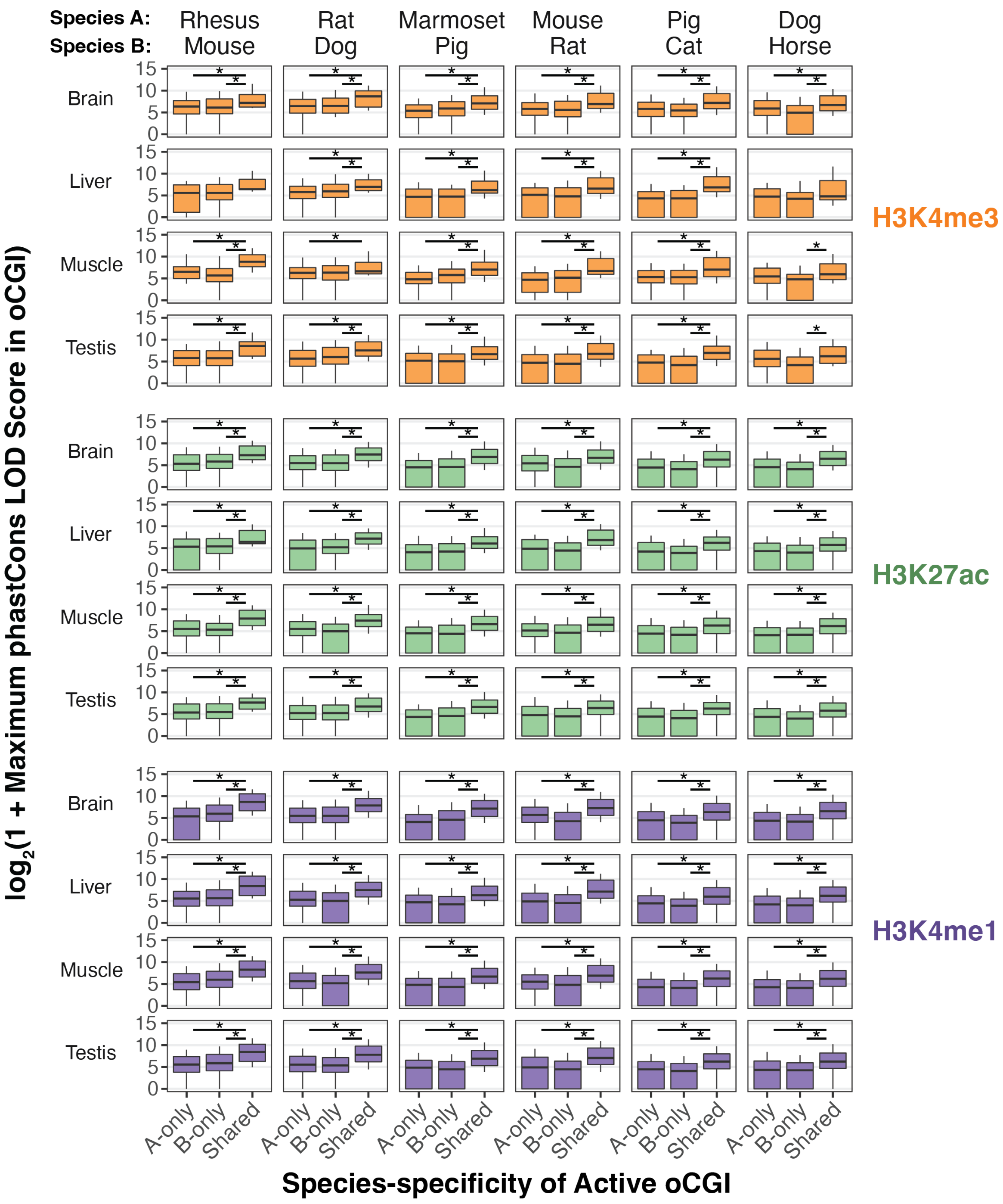


**Figure S25. Shared active oCGIs have higher maximum phastCons LOD scores**
Maximum phastCons LOD scores for species-specific oCGIs in species-specific peaks and shared oCGIs in shared peaks across all tissues and histone modifications for six species pairs. Box plots show the interquartile range and median, and whiskers indicate the 90% confidence interval. Stars indicate significant differences ($q < 0.05$ Wilcoxon rank-sum test, BH-corrected).

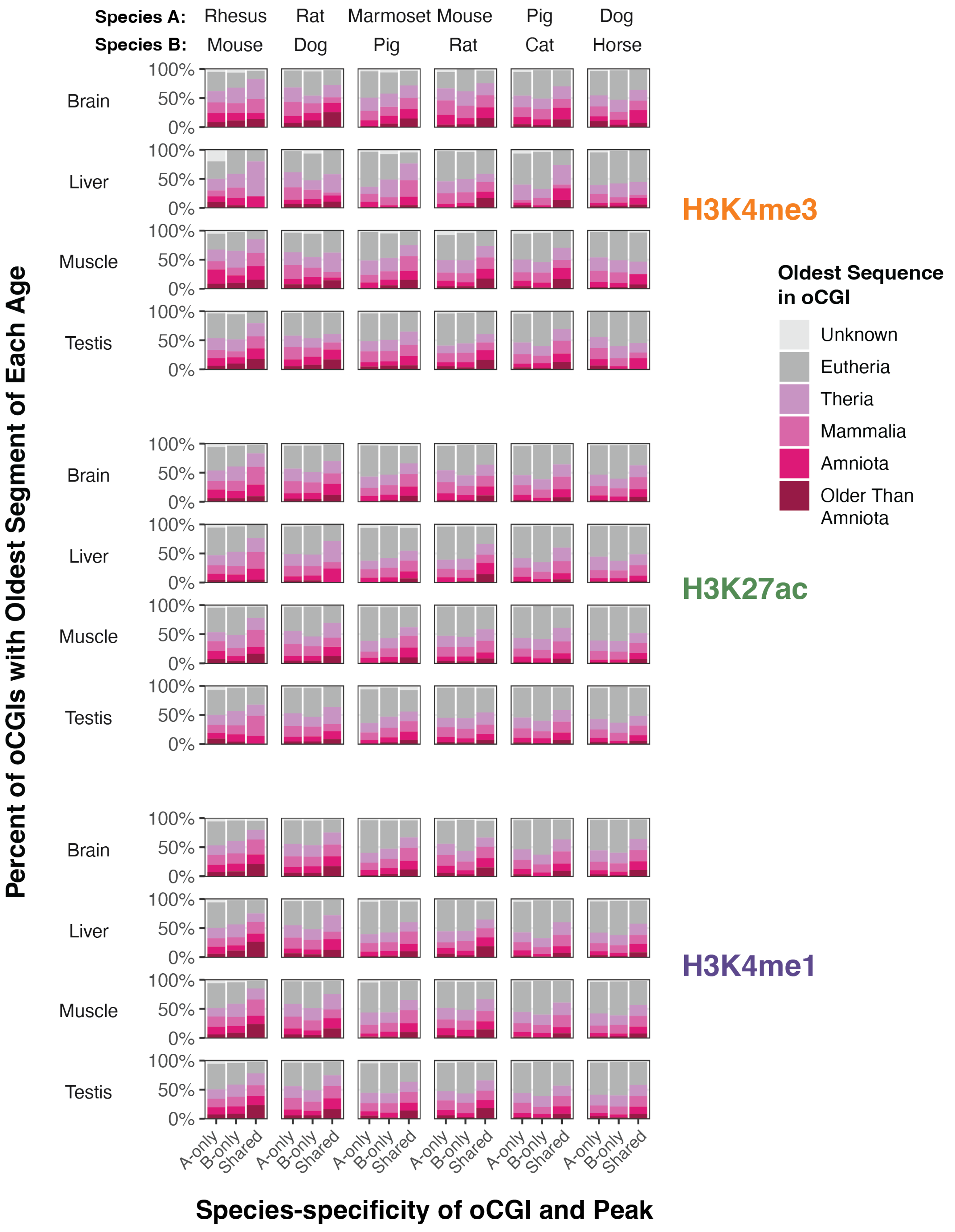


**Figure S26. Shared active oCGIs are present at older sequences**

The percent of species-specific oCGIs in species-specific peaks and shared oCGIs in shared peaks whose oldest sequence belongs to each age category (see legend). Results are shown for all tissues and histone modifications for six species pairs.

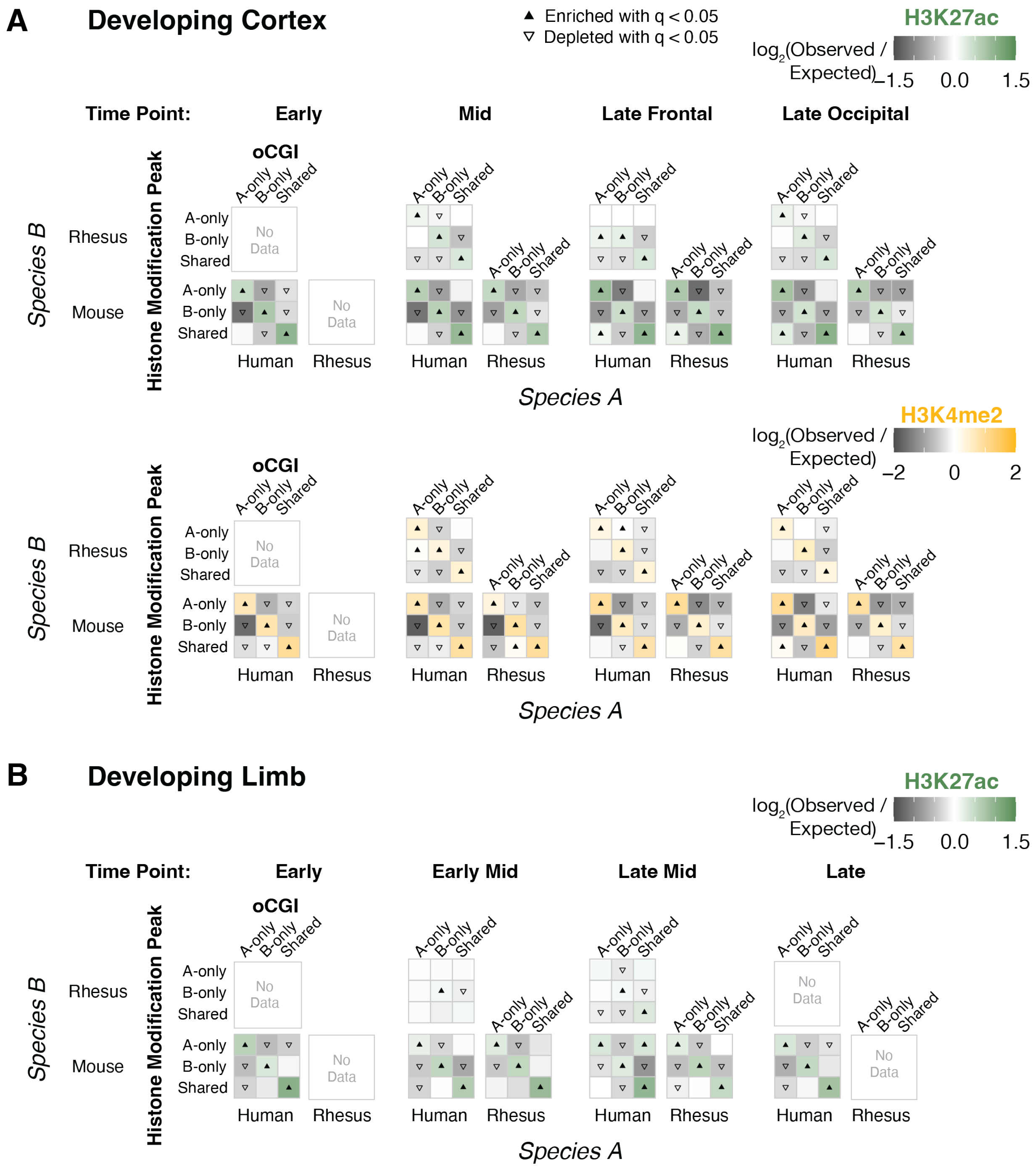

A
Developing Cortex
▲ Enriched with q < 0.05
▽ Depleted with q < 0.05
H3K27ac
log2(Observed / Expected)
−1.5
0.0
1.5
Time Point:
Early
Mid
Late Frontal
Late Occipital
oCGI
A-only
B-only
Shared
No Data
Species B
Rhesus
Mouse
Histone Modification Peak
Human
Rhesus
Species A
H3K4me2
−2
0
2
B
Developing Limb
Early Mid
Late Mid
Late

**Figure S27. oCGIs predict histone modification peaks in developing tissues**

(A) Enrichment and depletion in each indicated comparison of species-specific and shared oCGIs (*top:* A-only, B-only, Shared) and species-specific and shared peaks (*left*: A-only, B-only, Shared), compared to a null expectation of no association between oCGI turnover and peak turnover. Each 3 x 3 grid shows the results for a specific test examining oCGIs and their overlap with H3K27ac or H3K4me2 in developing cortex within a given species pair. Each box in each grid is colored according to the level of enrichment over expectation (green for H3K27ac, yellow for H3K4me2) or depletion (gray for both marks) of genome-wide sites that meet the criteria for that box. The color bars illustrate the level of enrichment or depletion over expectation. For visualization, values greater than the scale maximum (1.5 for H3K27ac, 2 for H3K4me2) and less than the scale minimum (-1.5 for H3K27ac, -2 for H3K4me2) were set to the scale maximum or minimum, respectively. The filled upward-pointing triangles denote significant enrichment and open downward-pointing triangles denote significant depletion ($q < 0.05$, permutation test, BH-corrected, see Fig. S17 and Methods). Developmental stages in each species for each time point are shown in Table S5. (B) Enrichment and depletion in each indicated comparison of species-specific and shared oCGIs (*top:* A-only, B-only, Shared) and species-specific and shared peaks (*left*: A-only, B-only, Shared), shown as in (A) but using H3K27ac peak data from developing limb.

#### Resampling Test to Measure Enrichment and Depletion of Each oCGI Species Pattern in HGEs Compared to non-HGEs

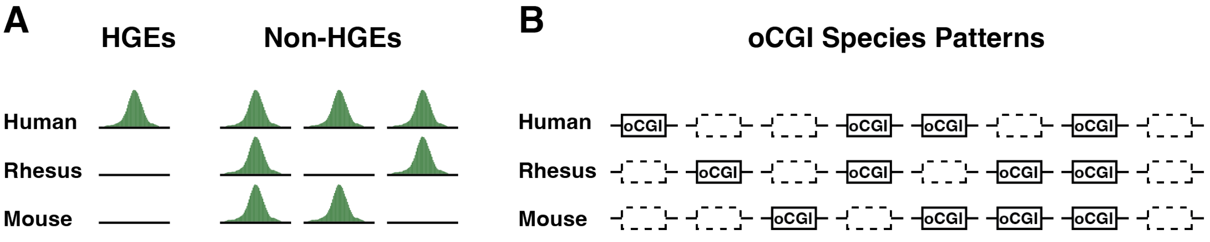


##### C Resampling Test

Performed Separately for Each Tissue and Time Point

1. Determine the percentage of HGEs, $P_{HGE}$, that belong to each category of oCGI species pattern.
2. Bin HGEs and non-HGEs by their total epigenetic signal in human, as measured by bigWigAverageOverBed. Bin sizes: 10 up to 2000, 100 up to the maximum total signal.
3. For the first bin, count the number HGEs, $n_{HGE}$, that fall in the bin. Randomly draw $n_{HGE}$ non-HGEs from the same bin.
4. Repeat step 3 across all bins, creating the "non-HGE Resampled Set" in which the total drawn non-HGEs is equal to the number of HGEs.
5. Calculate the percentage of the non-HGE Resampled Set that belong to each category of oCGI species pattern.
6. Repeat steps 3-5 for 20,000 resampling rounds to establish a distribution of percentages for each oCGI species pattern in the non-HGE Resampled Set, see panel (D).
7. For each oCGI species pattern, calculate the mean percentage of non-HGEs from all 20,000 resampling rounds that have that oCGI species pattern, $P_{non\text{-}HGE}$.
8. For each oCGI species pattern, plot $P_{HGE}$ and $P_{non\text{-}HGE}$ as bars (Fig. 4B) or calculate $\log_2(P_{HGE} / P_{non\text{-}HGE})$ (Fig. S29).
9. Calculate p-values for each oCGI species pattern

   Proportion of resampling rounds in which the percentage of the non-HGE Resampled Set with that oCGI species pattern is more extreme than $P_{HGE}$

   If this proportion is 0, then p = 1 / 20,000

**D**

Example: Resampling test comparing the percentage of HGEs and non-HGEs from 8.5 p.c.w. cortex that have a human-only oCGI

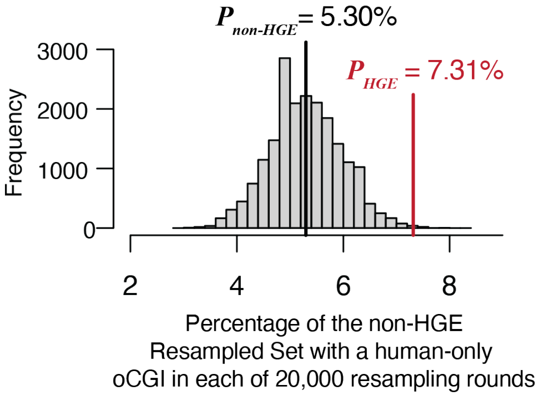

**Figure S28. Resampling test for analysis of oCGI species patterns in HGEs**

Explanation of resampling test to determine enrichment and depletion of each oCGI species pattern in HGEs compared to non-HGE human enhancers. (A) Schematic showing H3K27ac ChIP-seq peaks in human, rhesus macaque, and mouse for HGEs (*left*) compared to non-HGE human enhancers (*right*). (B) The oCGI species patterns considered in this analysis. Boxes with solid lines indicate oCGIs, and boxes with dashed lines indicate orthologous sequence that does not contain an oCGI. (C) Explanation of the resampling test used to compare HGEs to activity-matched non-HGEs. (D) Example results from a resampling test comparing the percent of HGEs that have a human-only oCGI to activity-matched non-HGE enhancers, using data from human cortex at 8.5 post-conception weeks (p.c.w.).

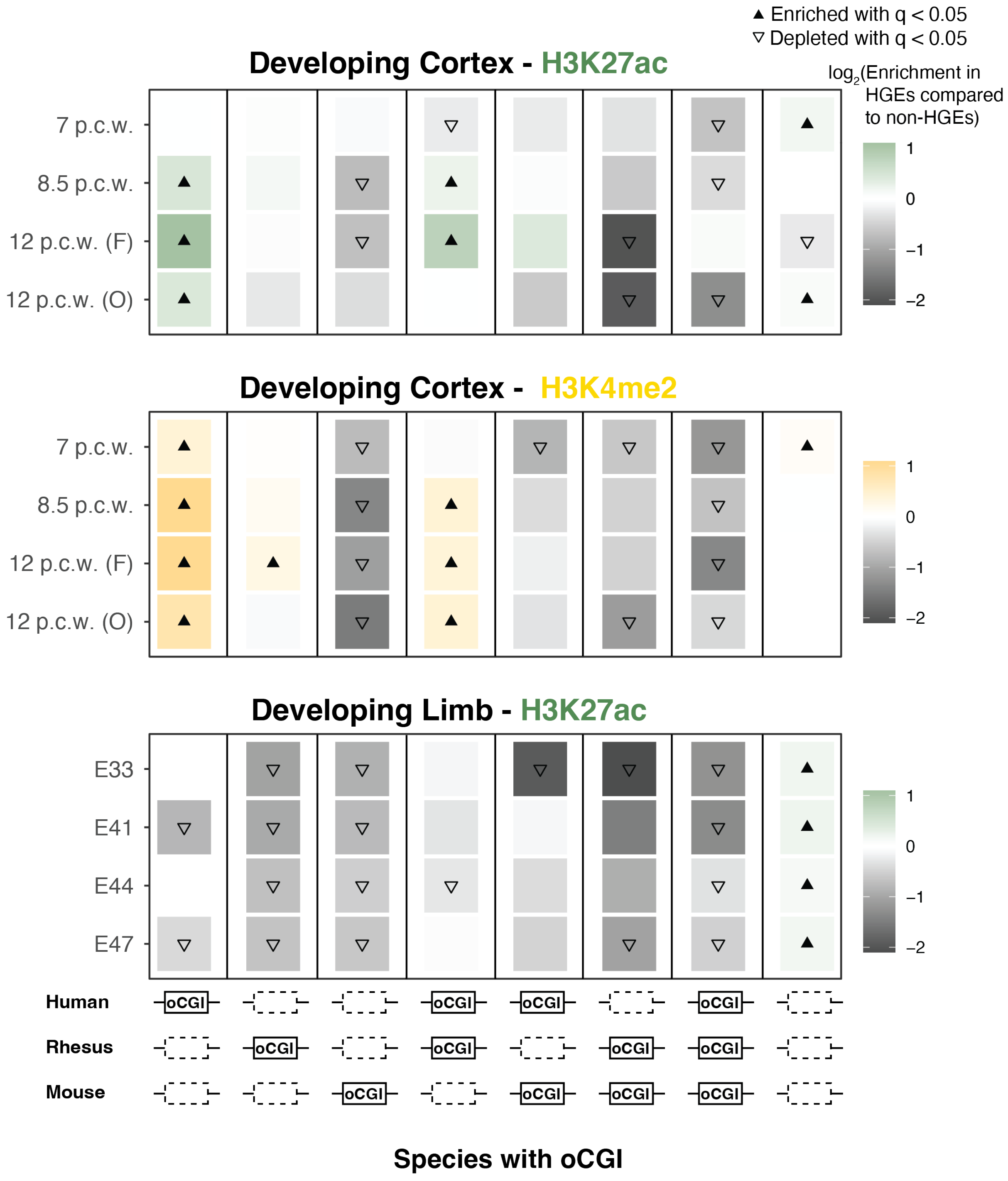
▲ Enriched with q < 0.05
▽ Depleted with q < 0.05
log2(Enrichment in HGEs compared to non-HGEs)
Developing Cortex - H3K27ac
7 p.c.w.
8.5 p.c.w.
12 p.c.w. (F)
12 p.c.w. (O)
1
0
−1
−2
Developing Cortex - H3K4me2
7 p.c.w.
8.5 p.c.w.
12 p.c.w. (F)
12 p.c.w. (O)
1
0
−1
−2
Developing Limb - H3K27ac
E33
E41
E44
E47
1
0
−1
−2
Human
Rhesus
Mouse
oCGI
Species with oCGI

**Figure S29. oCGI patterns enriched and depleted in HGEs**

Enrichment of each specified oCGI pattern (*bottom*) in HGEs compared to activity-matched non-HGE human enhancers, across all human tissues, marks, and time points. Boxes are colored according to their enrichment (green for H3K27ac, yellow for H3K4me2) or depletion (gray for both marks). The filled upward-pointing triangles denote significant enrichment and open downward-pointing triangles denote significant depletion ($q < 0.05$, resampling test comparing HGEs to activity-matched non-HGEs, BH-corrected; see Methods and Figure S28). p.c.w = post-conception weeks, F = frontal region, O = occipital region, E = embryonic day.

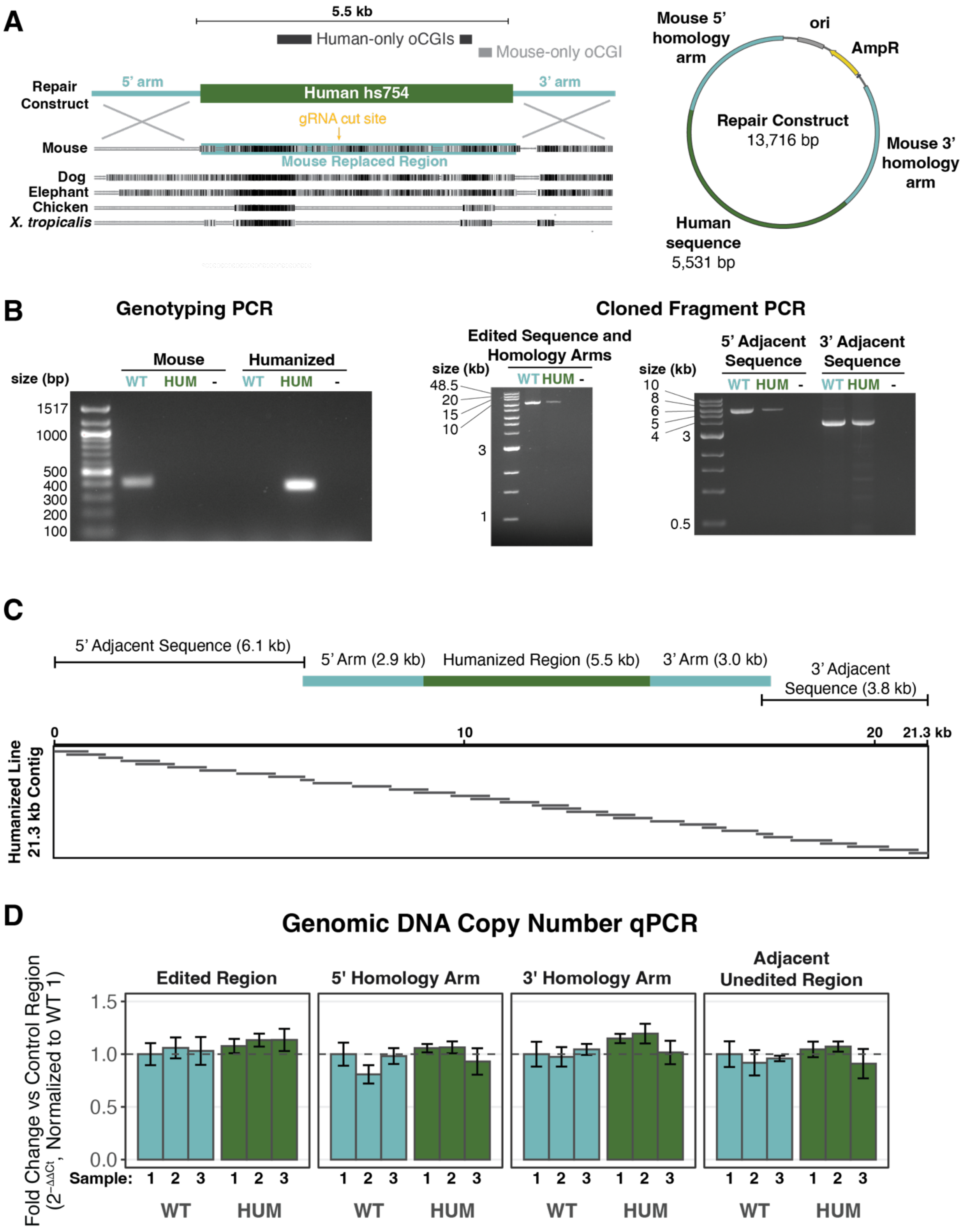
A
5.5 kb
Human-only oCGIs
Mouse-only oCGI
Repair Construct
5' arm
Human hs754
3' arm
gRNA cut site
Mouse
Mouse Replaced Region
Dog
Elephant
Chicken
X. tropicalis
Mouse 5' homology arm
ori
AmpR
Repair Construct
13,716 bp
Mouse 3' homology arm
Human sequence
5,531 bp
B
Genotyping PCR
Mouse
Humanized
size (bp)
WT HUM -
WT HUM -
1517
1000
500
400
300
200
100
Cloned Fragment PCR
Edited Sequence and Homology Arms
size (kb)
WT HUM -
48.5
20
15
10
3
1
5' Adjacent Sequence
3' Adjacent Sequence
size (kb)
WT HUM -
WT HUM -
10
8
6
5
4
3
0.5
C
5' Adjacent Sequence (6.1 kb)
5' Arm (2.9 kb)
Humanized Region (5.5 kb)
3' Arm (3.0 kb)
3' Adjacent Sequence (3.8 kb)
0
10
20
21.3 kb
Humanized Line 21.3 kb Contig
D
Genomic DNA Copy Number qPCR
Edited Region
5' Homology Arm
3' Homology Arm
Adjacent Unedited Region
Fold Change vs Control Region (2^-ΔΔCt, Normalized to WT 1)
1.5
1.0
0.5
0.0
Sample:
1 2 3 1 2 3
WT
HUM

**Figure S30. Generation and validation of the hs754 humanized mouse model hs754**

(A) *Left*: schematic of the humanized locus with human-based alignment to mouse, dog, elephant, chicken, and *Xenopus tropicalis* showing the humanized region, its highly conserved core sequence, and the location of the gRNA cut site. *Right*: the repair construct used to generate the humanized mouse line. (B) *Left*: Example genotyping PCR. Center: Amplification across the 5' homology arm, the humanized region, and the 3' homology arm. *Right*: Amplification of the mouse sequence adjacent to the 5' and 3' homology arms. (C) Alignment of Sanger sequencing reads across a 21.3-kb region centered on the humanized region on chromosome 13. (D) Genomic DNA copy number qPCR for the edited region, the homology arms, and an adjacent unedited region, normalized to a control region on chromosome 5. WT = wild type, HUM = humanized.

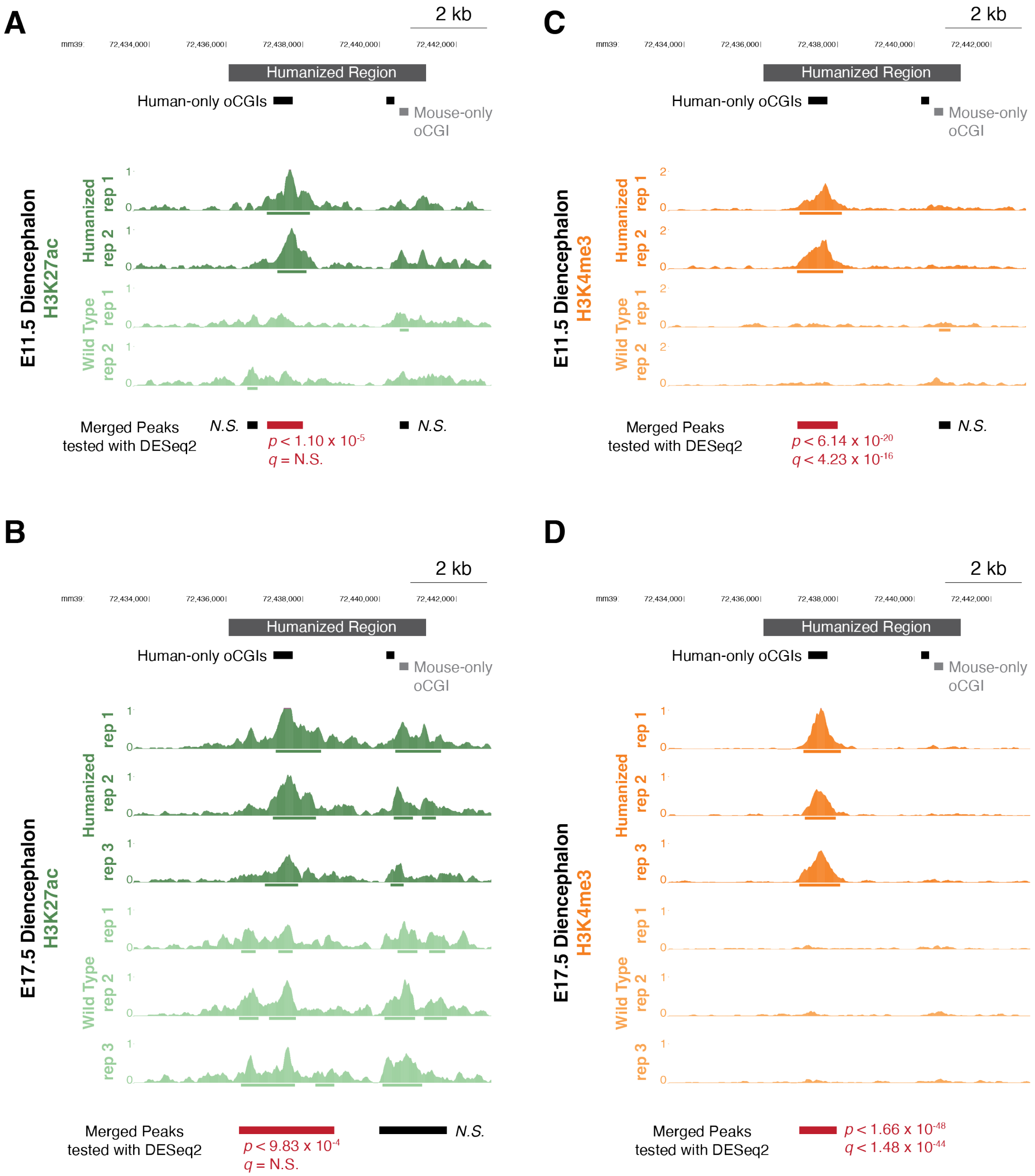
A
2 kb
mm39
72,434,000
72,436,000
72,438,000
72,440,000
72,442,000
Humanized Region
Human-only oCGIs
Mouse-only oCGI
E11.5 Diencephalon
H3K27ac
Humanized
rep 1
rep 2
Wild Type
rep 1
rep 2
Merged Peaks tested with DESeq2
N.S.
p < 1.10 x 10-5
q = N.S.
N.S.
C
2 kb
mm39
72,434,000
72,436,000
72,438,000
72,440,000
72,442,000
Humanized Region
Human-only oCGIs
Mouse-only oCGI
E11.5 Diencephalon
H3K4me3
Humanized
rep 1
rep 2
Wild Type
rep 1
rep 2
Merged Peaks tested with DESeq2
p < 6.14 x 10-20
q < 4.23 x 10-16
N.S.
B
2 kb
mm39
72,434,000
72,436,000
72,438,000
72,440,000
72,442,000
Humanized Region
Human-only oCGIs
Mouse-only oCGI
E17.5 Diencephalon
H3K27ac
Humanized
rep 1
rep 2
rep 3
Wild Type
rep 1
rep 2
rep 3
Merged Peaks tested with DESeq2
p < 9.83 x 10-4
q = N.S.
N.S.
D
2 kb
mm39
72,434,000
72,436,000
72,438,000
72,440,000
72,442,000
Humanized Region
Human-only oCGIs
Mouse-only oCGI
E17.5 Diencephalon
H3K4me3
Humanized
rep 1
rep 2
rep 3
Wild Type
rep 1
rep 2
rep 3
Merged Peaks tested with DESeq2
p < 1.66 x 10-48
q < 1.48 x 10-44

**Figure S31. Gain of H3K27ac and H3K4me3 associated with a human oCGI in a humanized mouse model, data for all replicates**

(A) H3K27ac levels for the humanized (dark green) and wild type (light green) mouse in embryonic day 11.5 (E11.5) diencephalon. Signal tracks display read counts per million in adjacent 10-bp bins. Bars under peaks show peak calls within each replicate. Merged peaks at the bottom show the union of peaks across all replicates and genotypes (shown in wild type coordinates), which were tested for differential levels between genotypes using DESeq2. Significance was determined using a Wald test to identify p-values, which were BH-corrected to q-values (both are shown). The humanized tracks have been shifted 190 bp to the left to align an orthologous base within the oCGI, due to overall differences in orthologous sequence lengths. (B) H3K27ac levels in E17.5 diencephalon, shown as in (A). (C) H3K4me3 levels in E11.5 diencephalon, shown as in (A) but with humanized tracks in dark orange and wild type tracks in light orange. (D) H3K4me3 levels in E17.5 diencephalon, shown as in (A) but with humanized tracks in dark orange and wild type tracks in light orange.

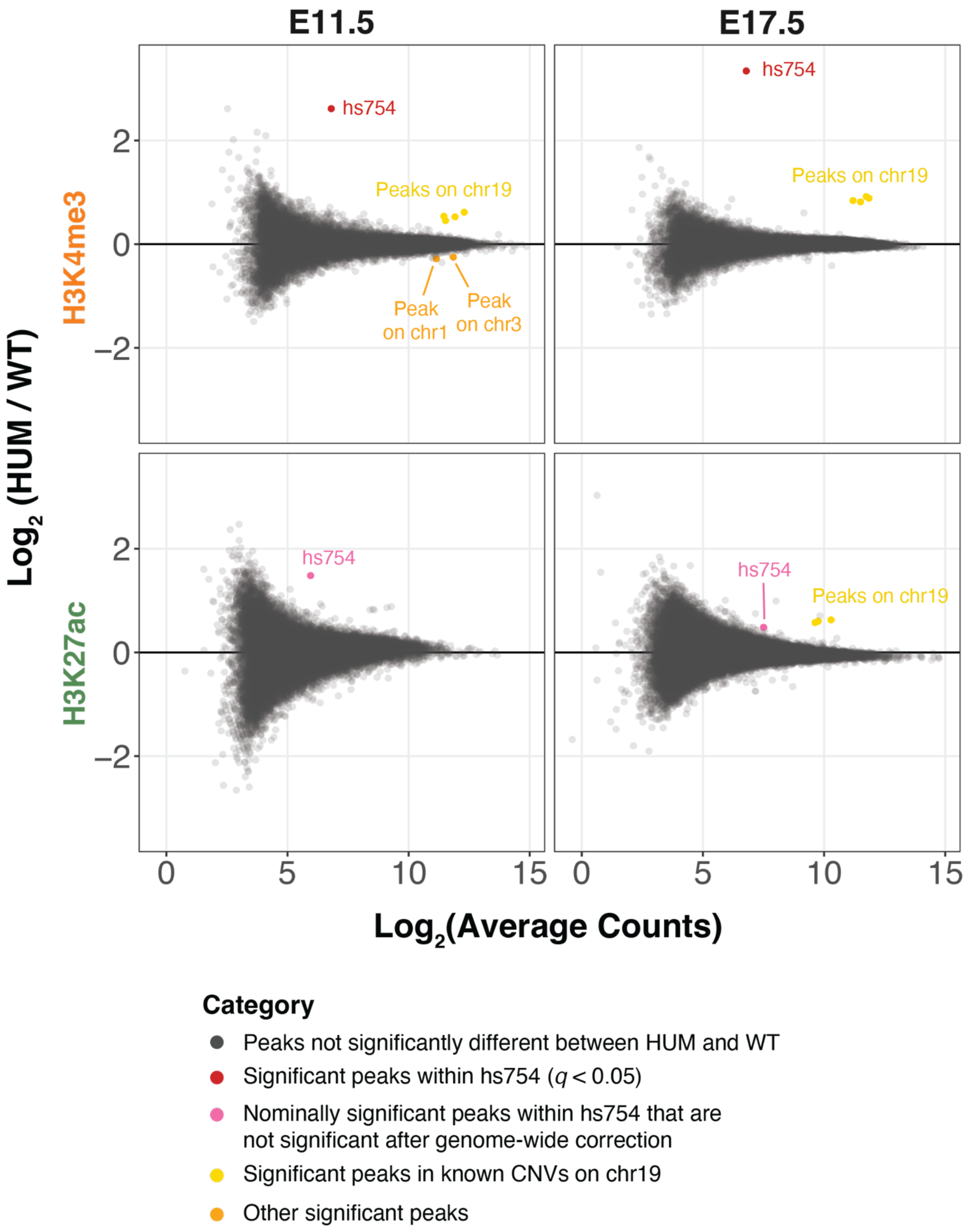
E11.5
E17.5
H3K4me3
H3K27ac
Log2 (HUM / WT)
Log2(Average Counts)
hs754
Peaks on chr19
Peak on chr1
Peak on chr3
2
0
−2
0
5
10
15
Category
Peaks not significantly different between HUM and WT
Significant peaks within hs754 (q < 0.05)
Nominally significant peaks within hs754 that are not significant after genome-wide correction
Significant peaks in known CNVs on chr19
Other significant peaks

**Figure S32. Genome-wide analysis of differential histone modification levels in the hs754 humanized mouse versus wild type**

$Log_2$-transformed average counts across all replicates and genotypes versus the $log_2$-transformed ratio of normalized reads between humanized (HUM) and wild type (WT) mouse diencephalon, at time points E11.5 (*left*) and E17.5 (*right*), for H3K4me3 (*top*) and H3K27ac (*bottom*). Differential peaks are indicated (see legend and Methods). See Fig S43 for additional information on the peaks on chr 19.

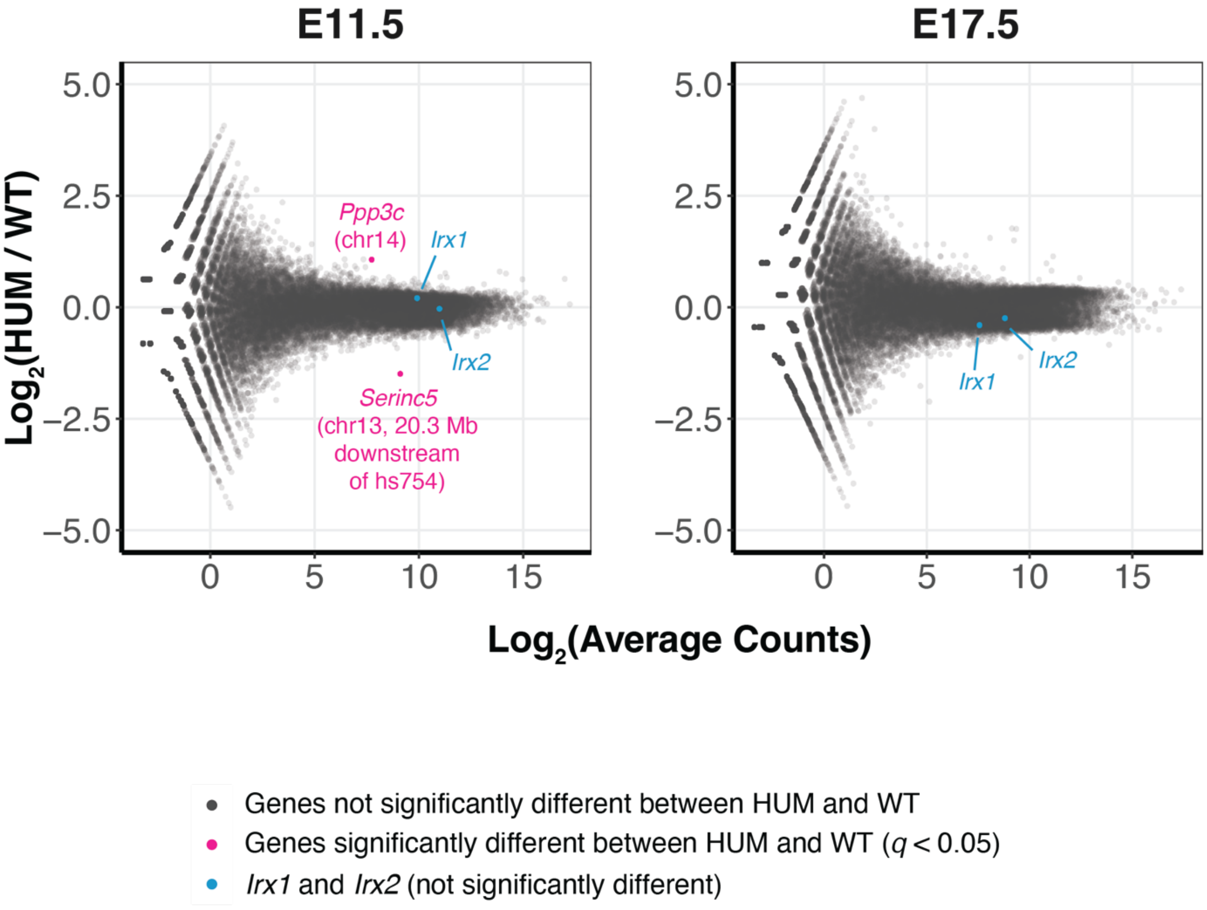


**Figure S33. Genome-wide analysis of differential gene expression in the hs754 humanized mouse versus wild type**
$Log_2$-transformed Average Counts across all replicates and genotypes versus the $Log_2$-transformed ratio of normalized reads between humanized (HUM) and wild type (WT) diencephalon at E11.5 (*left*) and E17.5 (*right*). All genes are shown in gray, with differentially expressed genes indicated in pink (see Methods). The putative target genes on either side of hs754 (*Irx1* and *Irx2*) are indicated in blue and are not called as differentially expressed.

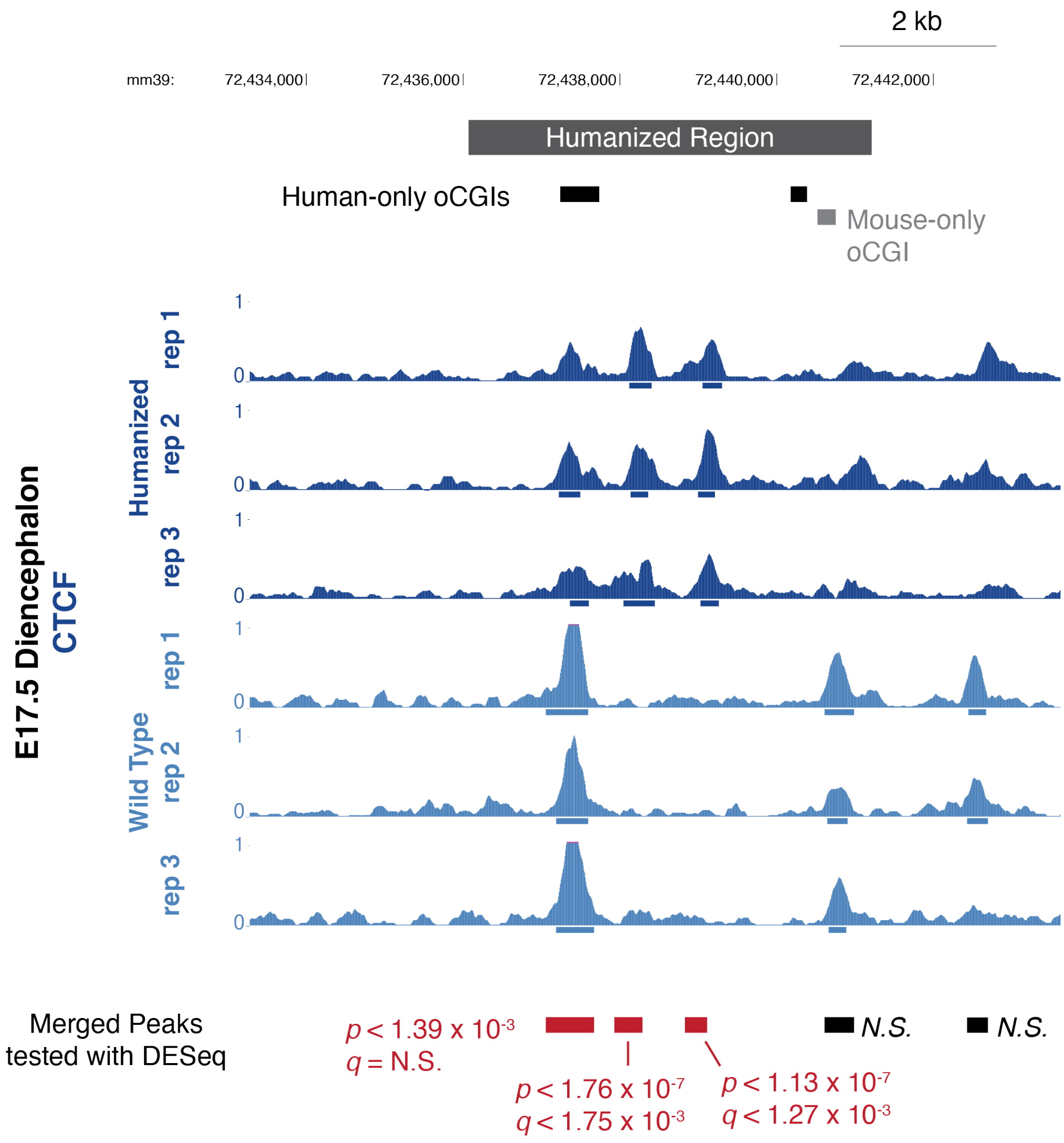


**Figure S34. The humanized locus recruits two additional CTCF binding events**
CTCF binding levels in developing diencephalon at embryonic day 17.5 (E17.5) from the humanized (dark blue) and wild type (light blue) mouse. Signal tracks display read counts per million in adjacent 10-bp bins. Bars under peaks show peak calls within each replicate. Merged peaks at the bottom show the union of peaks across all replicates and genotypes (shown in wild type coordinates), which were tested for differential signal between genotypes using DESeq2 (see Methods). Significance was determined using a Wald test to identify p-values, which were BH-corrected to q-values (both are shown). The humanized tracks have been shifted 190 bp to the left to align an orthologous base within the oCGI, due to overall differences in orthologous sequence lengths.

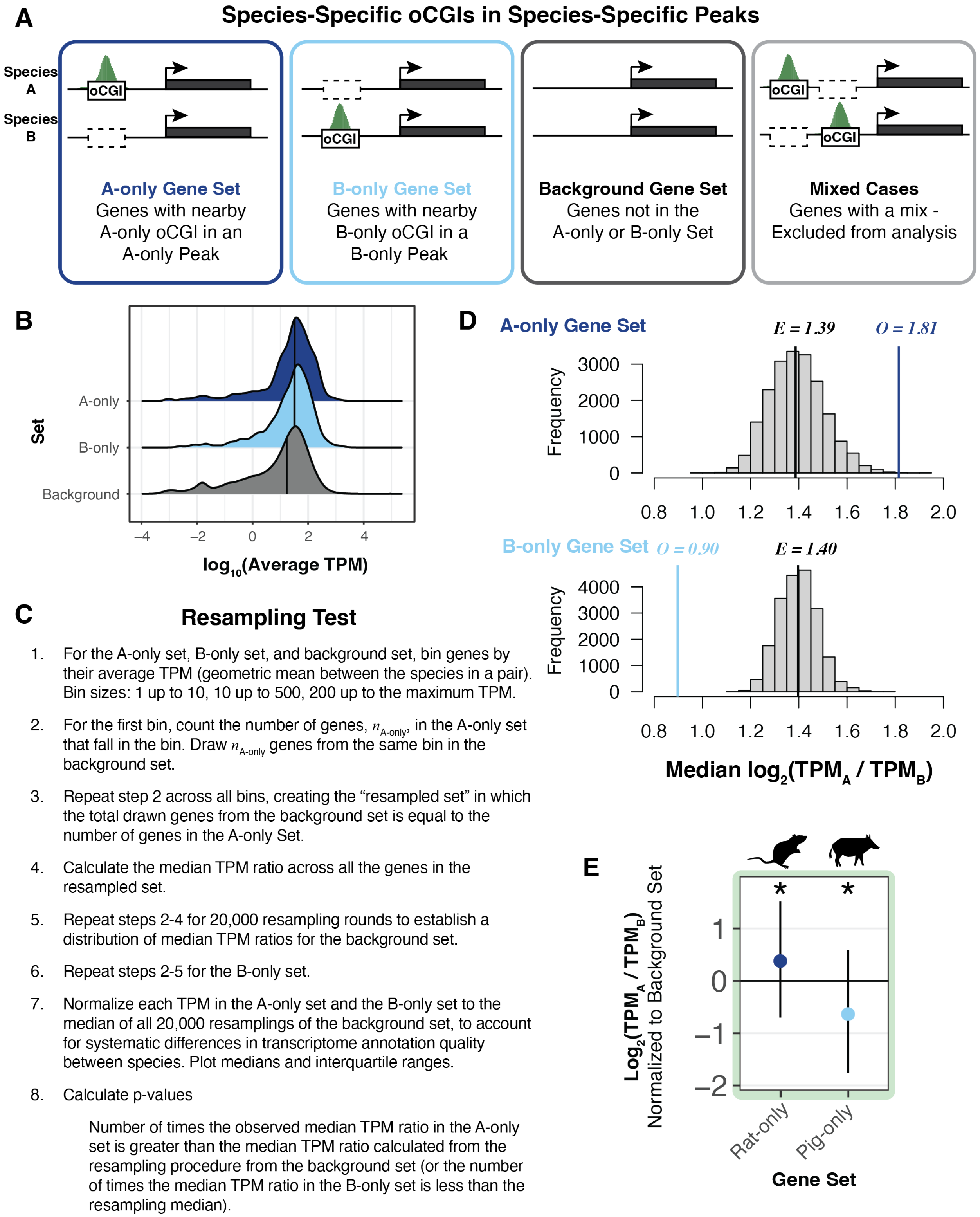
A
Species-Specific oCGIs in Species-Specific Peaks
Species A
Species B
oCGI
A-only Gene Set
Genes with nearby A-only oCGI in an A-only Peak
B-only Gene Set
Genes with nearby B-only oCGI in a B-only Peak
Background Gene Set
Genes not in the A-only or B-only Set
Mixed Cases
Genes with a mix - Excluded from analysis
B
Set
A-only
B-only
Background
–4
–2
0
2
4
log10(Average TPM)
C
Resampling Test
1. For the A-only set, B-only set, and background set, bin genes by their average TPM (geometric mean between the species in a pair). Bin sizes: 1 up to 10, 10 up to 500, 200 up to the maximum TPM.
2. For the first bin, count the number of genes, n_A-only, in the A-only set that fall in the bin. Draw n_A-only genes from the same bin in the background set.
3. Repeat step 2 across all bins, creating the “resampled set” in which the total drawn genes from the background set is equal to the number of genes in the A-only Set.
4. Calculate the median TPM ratio across all the genes in the resampled set.
5. Repeat steps 2-4 for 20,000 resampling rounds to establish a distribution of median TPM ratios for the background set.
6. Repeat steps 2-5 for the B-only set.
7. Normalize each TPM in the A-only set and the B-only set to the median of all 20,000 resamplings of the background set, to account for systematic differences in transcriptome annotation quality between species. Plot medians and interquartile ranges.
8. Calculate p-values
Number of times the observed median TPM ratio in the A-only set is greater than the median TPM ratio calculated from the resampling procedure from the background set (or the number of times the median TPM ratio in the B-only set is less than the resampling median).
D
A-only Gene Set
E = 1.39
O = 1.81
Frequency
3000
2000
1000
0
0.8
1.0
1.2
1.4
1.6
1.8
2.0
B-only Gene Set
O = 0.90
E = 1.40
Frequency
4000
3000
2000
1000
0
0.8
1.0
1.2
1.4
1.6
1.8
2.0
Median log2(TPMA / TPMB)
E
Log2(TPMA / TPMB)
Normalized to Background Set
*
*
1
0
–1
–2
Rat-only
Pig-only
Gene Set

**Figure S35. Analysis pipeline for species-specific active oCGIs and gene expression**

(A) Gene sets defined in this analysis based on their linear proximity to species-specific oCGIs in species-specific peaks. (B) Density plot showing the average TPM of genes in the A-only, B-only, and background sets using H3K27ac data from adult brain in rat versus pig. Medians are indicated with vertical black lines. The medians for the A-only and B-only set are higher than the background set, motivating the use of a binned resampling test to compare the A-only set and B-only set to the background set, as explained in the remainder of the figure. (C) Explanation of binned resampling test to compare the medians of the A-only set and the B-only set to expected values generated by expression-matched resampling from the background set. (D) Example resampling results for the comparison between rat and pig for genes associated with species-specific oCGIs in species-specific H3K27ac peaks in adult brain. Histograms show the distribution of median $log_2$-transformed TPM ratios across 20,000 resampling rounds, in each of which expression-matched genes were drawn from the background set. (E) Example results plot for the same comparison as in panel (D), between rat and pig using H3K27ac peaks from adult brain. All values in the rat-only set and pig-only set are normalized to the median of 20,000 resampling medians from the background set.

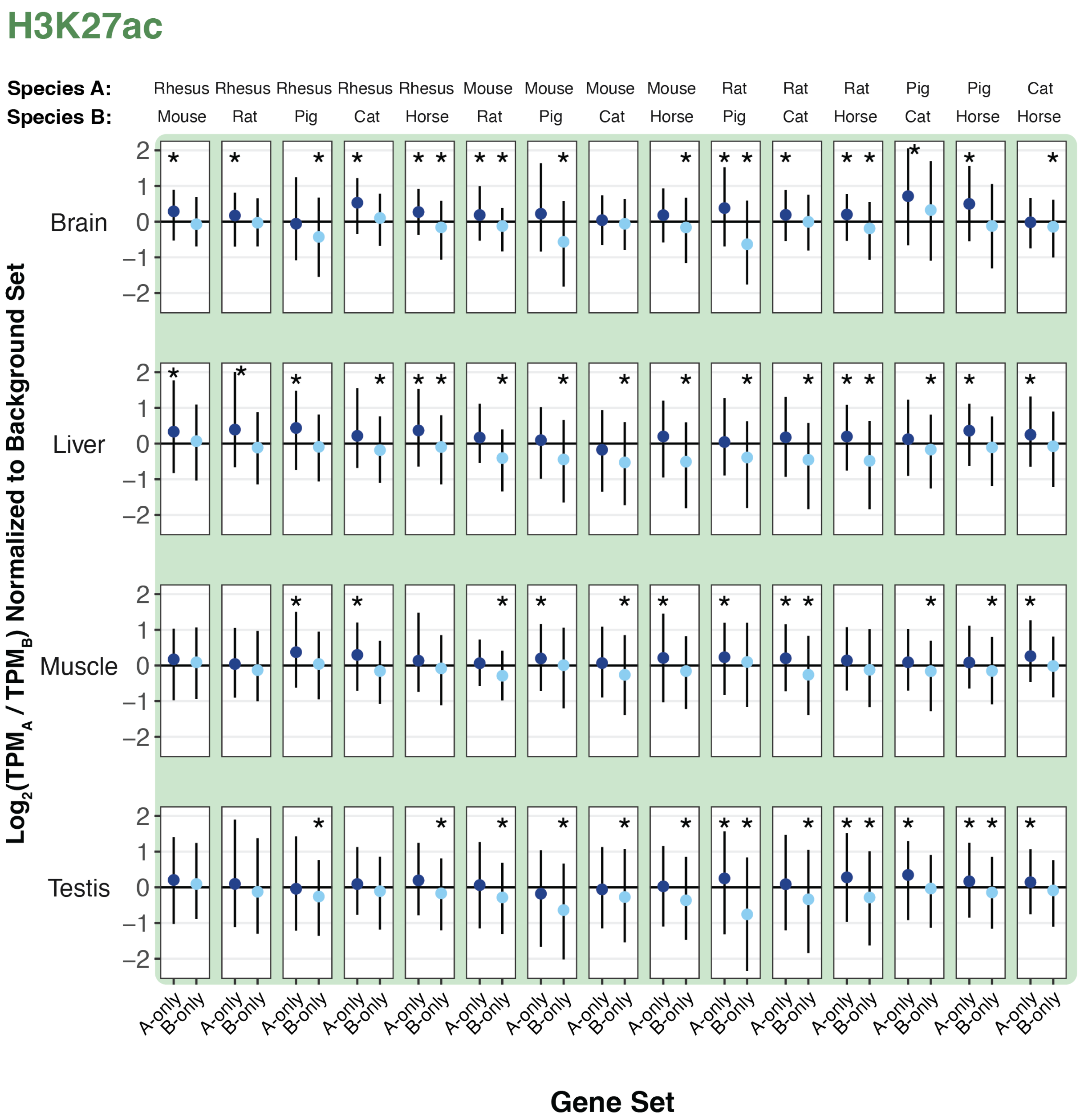


**Figure S36. Species-specific oCGIs in species-specific H3K27ac peaks are associated with gene expression changes**

The TPM ratio between species A and species B for genes in the A-only set and the B-only set. Results are shown for all species pairs and adult tissues, when genes are sorted based on nearby species-specific oCGIs in species-specific H3K27ac peaks. Points indicate median values for genes in the A-only set (dark blue) and genes in the B-only set (light blue). Lines indicate the interquartile range. All values in the A-only set and B-only set are normalized to the median of 20,000 resampling medians from the background set. Stars indicate a significant difference between the observed median and the expected median ($q < 0.05$, resampling test to compare to the background set, see Fig S35 and Methods).

## H3K4me1

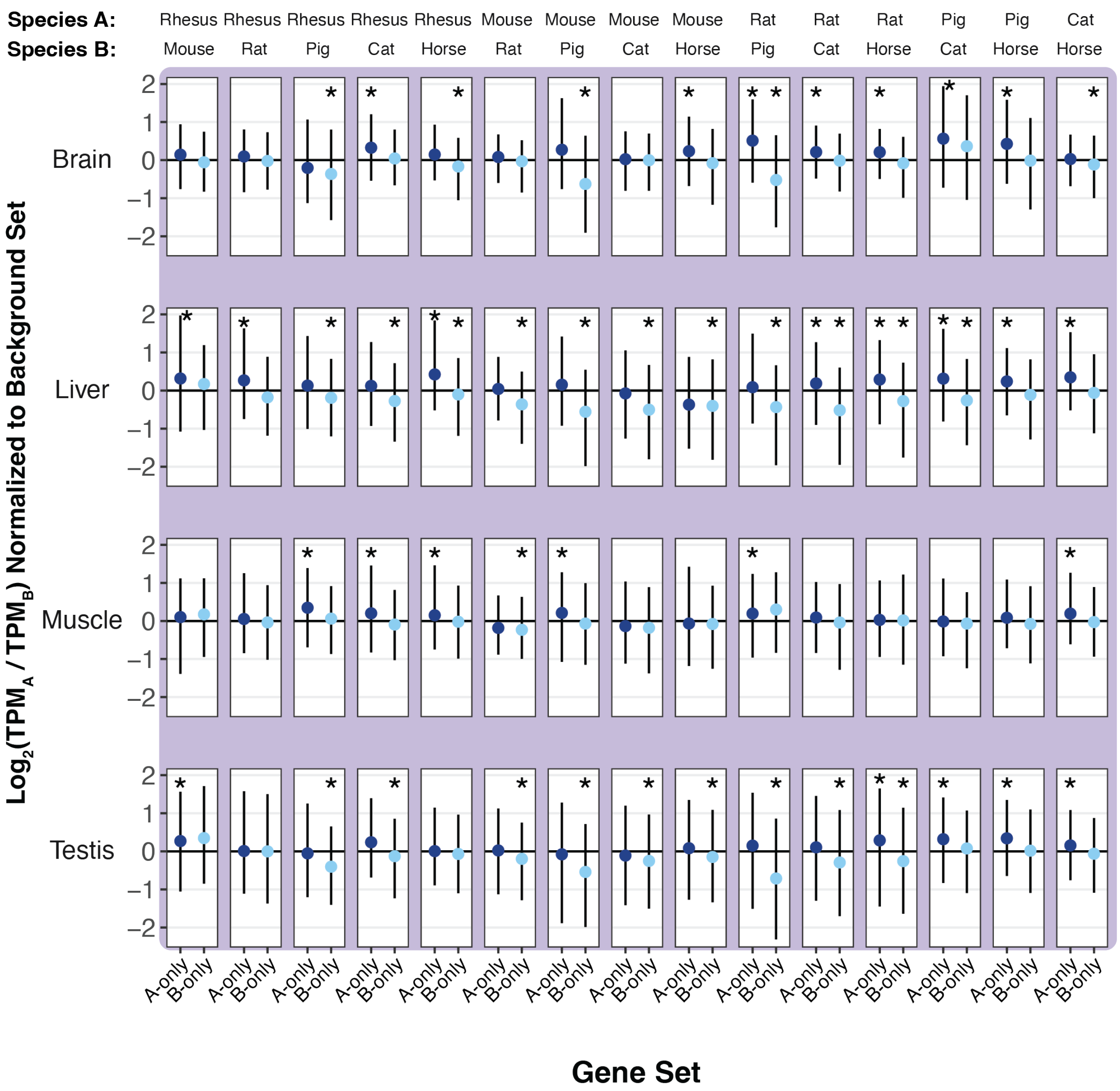


**Figure S37. Species-specific oCGIs in species-specific H3K4me1 peaks are associated with gene expression changes**

The TPM ratio between species A and species B for genes in the A-only set and the B-only set. Results are shown for all species pairs and adult tissues, when genes are sorted based on nearby species-specific oCGIs in species-specific H3K4me1 peaks. Points indicate median values for genes in the A-only set (dark blue) and genes in the B-only set (light blue). Lines indicate the interquartile range. All values in the A-only set and B-only set are normalized to the median of 20,000 resampling medians from the background set. Stars indicate a significant difference between the observed median and the expected median ($q < 0.05$, resampling test to compare to the background set, see Fig S35 and Methods).

## H3K4me3

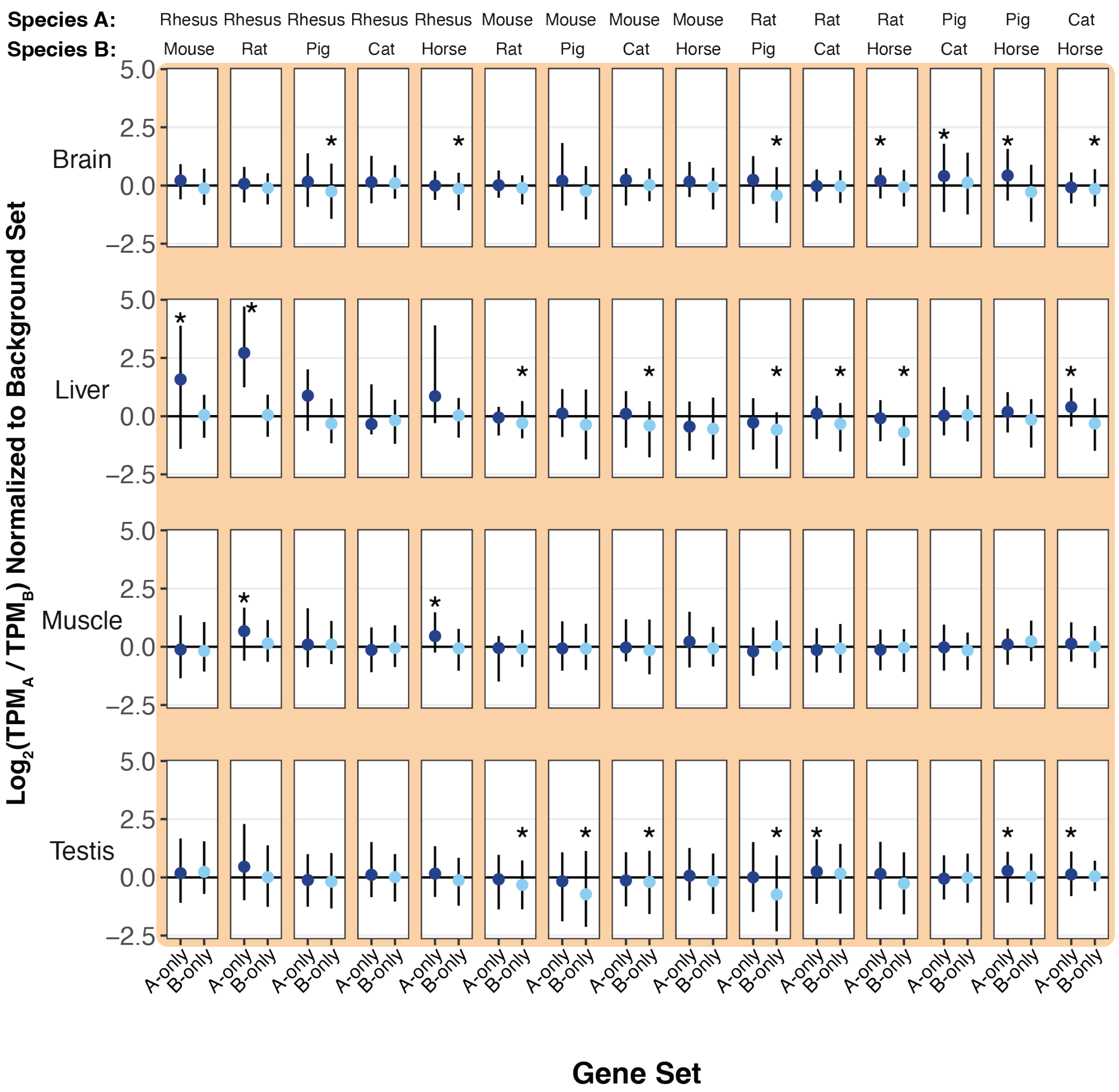


**Figure S38. Species-specific oCGIs in species-specific H3K4me3 peaks are associated with gene expression changes**

The TPM ratio between species A and species B for genes in the A-only set and the B-only set. Results are shown for all species pairs and adult tissues, when genes are sorted based on nearby species-specific oCGIs in species-specific H3K4me3 peaks. Points indicate median values for genes in the A-only set (dark blue) and genes in the B-only set (light blue). Lines indicate the interquartile range. All values in the A-only set and B-only set are normalized to the median of 20,000 resampling medians from the background set. Stars indicate a significant difference between the observed median and the expected median ($q < 0.05$, resampling test to compare to the background set, see Fig S35 and Methods).

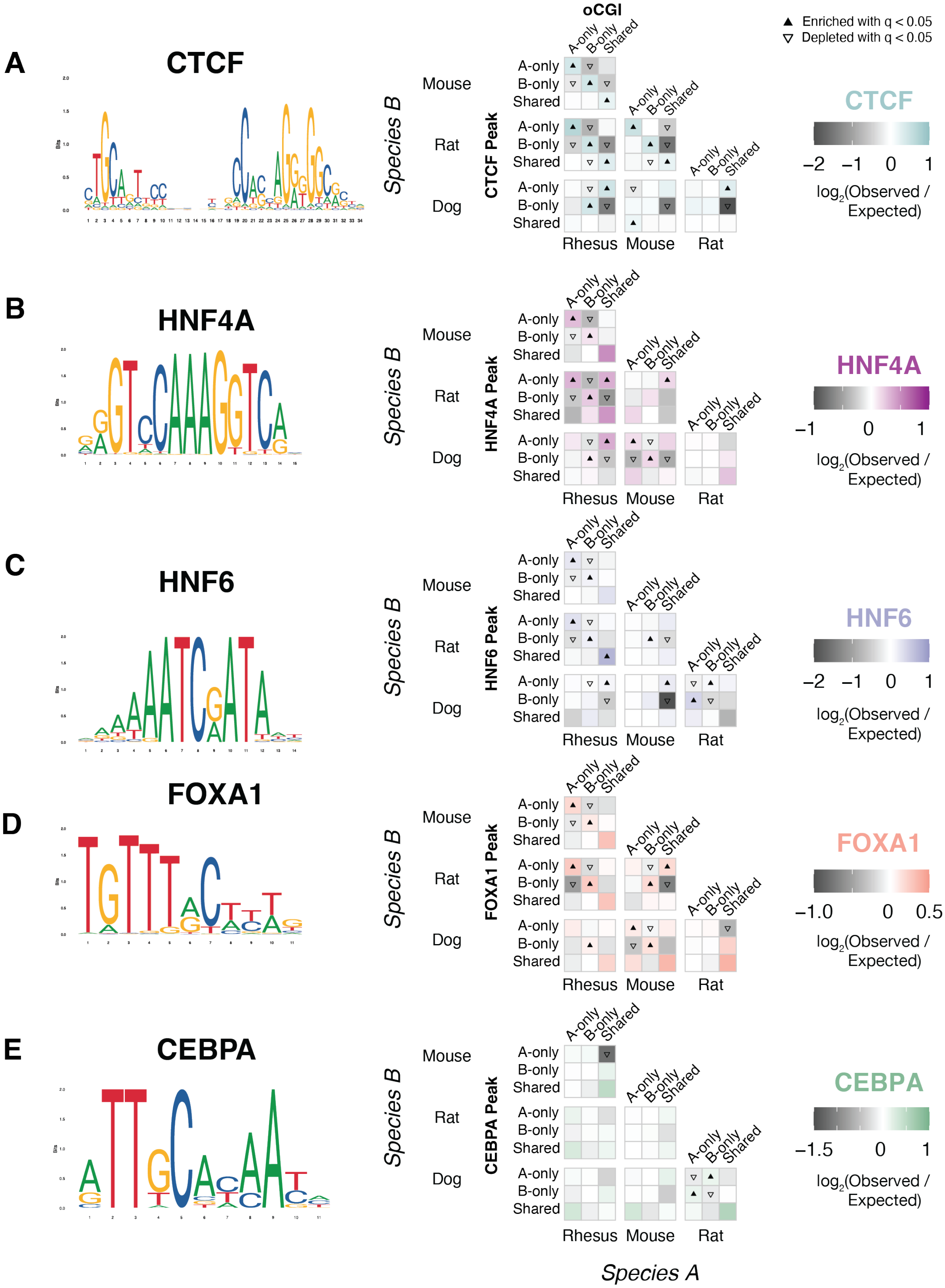
A
CTCF
B
HNF4A
C
HNF6
D
FOXA1
E
CEBPA
oCGI
Enriched with q < 0.05
Depleted with q < 0.05
A-only
B-only
Shared
Mouse
Rat
Dog
Species B
CTCF Peak
HNF4A Peak
HNF6 Peak
FOXA1 Peak
CEBPA Peak
Rhesus
Mouse
Rat
Species A
−2 −1 0 1
−1 0 1
−1.0 0 0.5
−1.5 0 1
log2(Observed / Expected)

**Figure S39. Enrichment and depletion results for all investigated TFs**

*Left*: consensus motifs for each TF. Motifs are from the JASPAR database. CTCF: MA1929.1, FOXA1: MA0148.1, HNF4A: MA1494.1, HNF6: MA0679.1, CEBPA: MA0102.3. *Right*: Enrichment and depletion in each indicated comparison of species-specific and shared oCGIs (*top:* A-only, B-only, Shared) and species-specific and shared TF peaks (*left*: A-only, B-only, Shared), compared to a null expectation of no association between oCGI turnover and peak turnover. Each 3 x 3 grid shows the results for a specific test examining oCGIs and their overlap with TF peaks in adult liver within a given species pair. Each box in each grid is colored according to the level of enrichment over expectation (teal for CTCF, magenta for HNF4A, purple for HNF6, red for FOXA1, and green for CEBPA) or depletion (gray for all TFs) of genome-wide sites that meet the criteria for that box. The color bars illustrate the level of enrichment or depletion over expectation. The filled upward-pointing triangles denote significant enrichment and open downward-pointing triangles denote significant depletion ($q < 0.05$, permutation test, BH-corrected; see Fig. S17 and Methods).

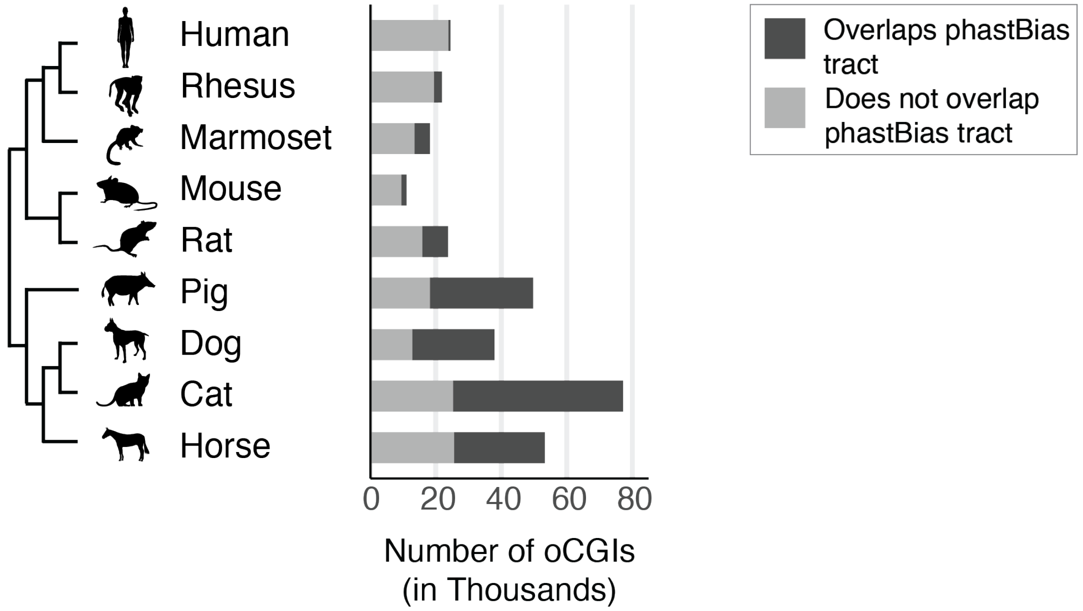


**Figure S40. GC-biased gene conversion tracts overlap many oCGIs**
Bar plots show the number of oCGIs in each species, with sites overlapping a phastBias tract in dark gray and sites not overlapping a phastBias tract in light gray.

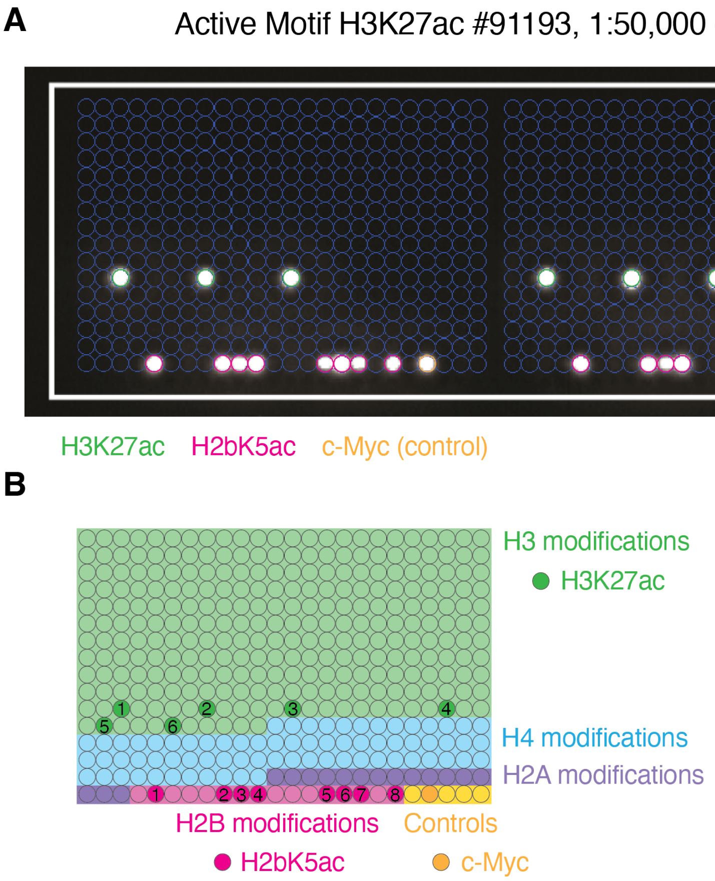


*H3 modifications:*

1. K27ac
2. K27ac + R26me2s
3. K27ac + R26me2a
4. K27ac + S28P
5. K27ac + S28P + R26me2s
6. K27ac + S28P + R26me2a

*H2B modifications:*

1. K5ac
2. K5ac + K12ac
3. K5ac + S14P
4. K5ac + K15ac
5. K5ac + K12ac + S14P
6. K5ac + K12ac + K15ac
7. K5ac + S14P + K15ac
8. K5ac + K12ac + S14P + K15ac

**Figure S41. Specificity of H3K27ac antibody used for humanized mouse ChIP-seq**
(A) Results of an analysis of H3K27ac antibody specificity using the MODified peptide array from Active Motif. Each blue-outlined circle shows the location of a peptide containing one or several posttranslational modifications. White signal shows peptides bound by the antibody of interest. Circles show the locations of peptides containing H3K27ac (green) or H2bK5ac (magenta), or the positive control antigen c-Myc (yellow). (B) Key showing locations of peptides from different histones (green background for histone H3, blue background for histone H4, purple background for histone H2A, pink background for histone H2B, yellow for control antigens). Circles filled with darker green show peptides with H3K27ac, and the additional modifications on these six peptides are listed to the right. Circles filled with darker pink show peptides with H2bK5ac, and the additional modifications on these eight peptides are listed below. Circle filled with darker yellow shows the location of the c-myc control antigen.

### Identifying Orthologous Peaks between Wild Type and Humanized Mouse

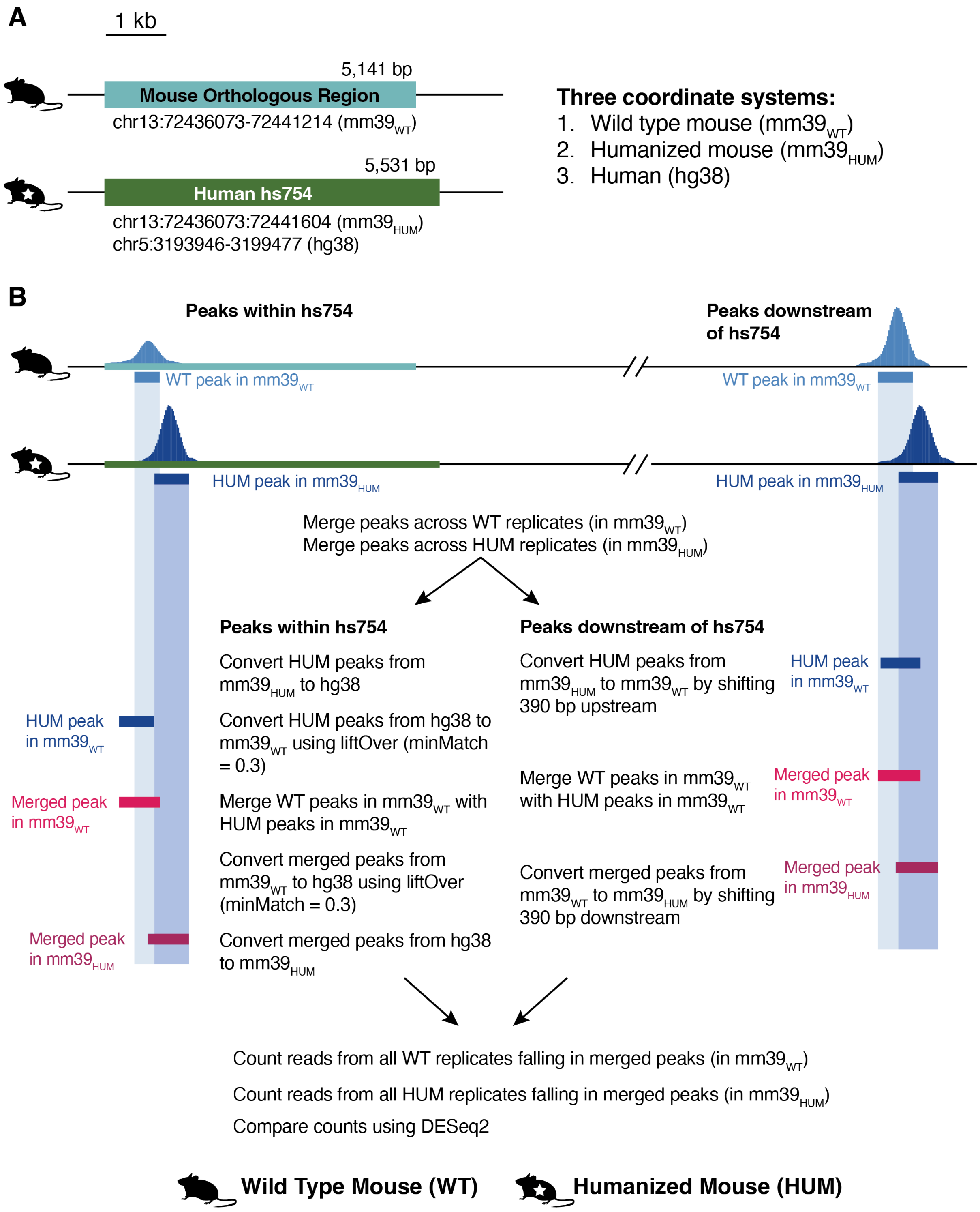

**Figure S42. Converting peak coordinates between wild type and humanized mouse**
(A) Schematic of the hs754 humanized locus and orthologous wild type locus, showing the size difference that requires a special procedure for comparing peaks between the two genotypes. Coordinates in the wild type mouse locus are referred to in mm39, denoted here as $mm39_{WT}$. Coordinates in the humanized mouse locus can be referred to either in humanized mouse coordinates ($mm39_{HUM}$) or human coordinates (hg38). (B) Procedure for comparing peaks between the wild type and humanized mouse loci, either within the humanized locus (*left*) or downstream of it on chromosome 13 (*right*).

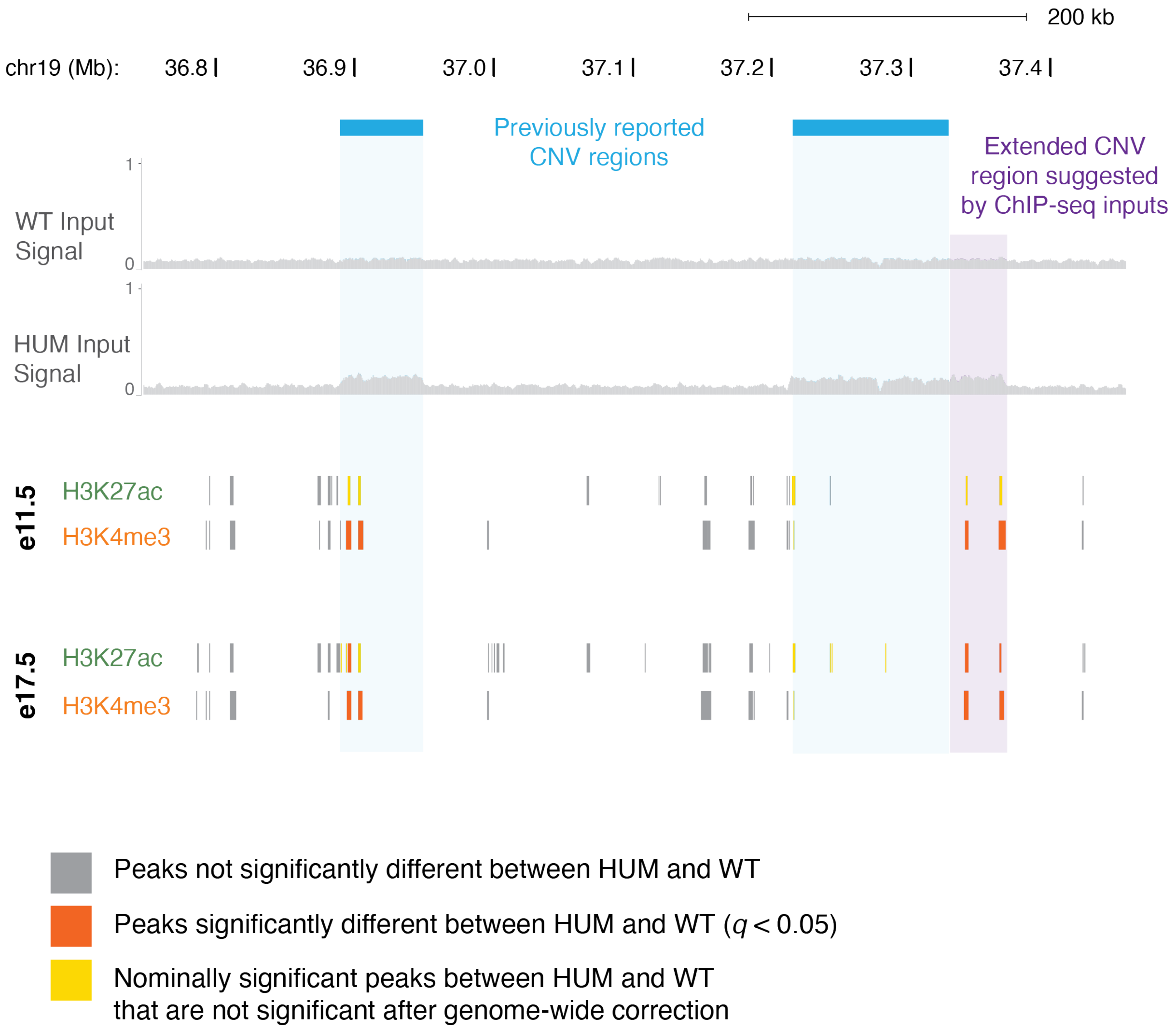


**Figure S43. Copy-number variants (CNVs) on chromosome 19**
Two genomic intervals on chromosome 19 containing known CNVs (Watkins-Chow & Pavan, 2008). ChIP input signal tracks from this study are shown in gray (from one replicate of wild type (WT) and humanized (HUM) E11.5 diencephalon) and suggest a duplicated signal in these regions (light blue), and in an extended region adjacent to the larger reported region (purple). All differential peaks on chromosome 19 that we called in this study fall within these CNV regions.
